# Dermal bone remodelling by invasive osteoblasts evolved in stem gnathostomes

**DOI:** 10.64898/2026.09.16.752034

**Authors:** Kate Weymouth-Crocker Jordan, Jason Downs, Xintao Zhang, Hiromi Yanagisawa, David E. Clouthier, Philip C.J. Donoghue, Georgy Koentges

## Abstract

The evolutionary origin of the human face can be traced to the earliest vertebrates with a dermal skeleton in which canonical skeletal tissues and cell types arose. To elucidate its developmental evolution we performed genetic mosaic labelling, 3D single cell analysis and fluorescent matrix birthdating *in vivo*. We discover a common lineage origin of endothelial cells and osteoblasts and *de novo* vasculogenesis (not angiogenesis) as the dominant mechanism. Inside the spongy (cancellous) layer, invasive OPN+/RUNX2+ osteoblasts establish two orthogonal collagen scaffolds, remodel them by directional secretion of matrix metalloproteinases and employ controllers of hydroxyapatite crystal resorption previously considered to be osteoclast-specific. Such intercalary biomineralization, comprising cellular sheet and volumetric biomineral expansion, renders all layers malleable even during postnatal stages. We trace this mechanism to bone ontogenies of the earliest skeletonizing vertebrates and resolve cell/tissue homologies controversial for almost two centuries. Our new model also has implications for bone matrix bioengineering and cancer osteomimicry that might replay some of these ontogenetic processes.

## Introduction

Dermal bone is a shared primitive feature of vertebrates with a mineralized skeleton. Three layers are traditionally recognized, a superficial compact dermis layer (D), a spongy layer and a deep compact layer. The fossil record of the earliest dermal skeleton comprises a dizzying array of (often acellular) matrix histologies, incompatible with the axiomatic model of apposition. Epithelio-mesenchymal interactions shaping the D layer were considered to dominate the development of the underlying bone, leading to the odontode theory^1-6^. Odontode theory treats the supporting cancellar and compact dermal bone as a passive bystander, governed by dermis angiogenesis^7,8^ which has not been experimentally verified. Most extant vertebrate skull bones bear no odontodes (dermal denticles composed of dentine and enameloid) and some ancestral vertebrates have odontodes but no underlying bone, suggesting that odontodes and dermal bone are phylogenetically and developmentally distinct ^9^. Similarly, while sutures are perceived as central places of cranial bone growth in mammals^10^, they cannot account for dermal bone growth in the majority of primitive vertebrates.

The molecular data in the accompanying paper established a 3-layered architecture underneath the dermis (D) layer: a compact layer L1, the spongy L2 and the compact L3. Our discovery of L1 as a generative layer provides the opportunity to re-examine fossil evidence. In mice a trilayered bone organization (L1,2,3) derives from a bilayered configuration (L1,3), dominated by the radial growth of the spongy L 2. Here we examine the molecular architecture and growth mechanism of this spongy layer 2 in relation to vasculature and biomineralization. We identify a key cell type responsible and describe the ultrastructural hallmarks of its behaviour in pristinely preserved placoderm and heterostracan ontogenetic stages which shed light onto the ancestry of this mechanism.

## Results

### Vasculogenesis in dermal bone growth

Current models of dermal bone formation posit that the characteristic architecture of dermal bone has been shaped ontogenetically by developing around preceding vasculature that sprouts angiogenetically from primary dermis vessels^7,8,11^. To validate this we determined the ontogenetic and spatial relationship between Runx2+ osteoblasts and CD31+ endothelial cells early in L2. We generated a new recombinase reporter with a membrane-bound vGFP enabling us to visualize the full cellular morphology of neural crest (NC) cells (see ESM5). 3D analysis of the frontal/clavicular bones at E15 reveals that existing vessels of the D layer are not connected to L2 as an angiogenetic mechanism would require (Fig.1a). Instead, we find large numbers of individualized CD31+ cells of NC origin (Fig.1a,b), commensurate with *de novo* vasculogenesis within L2. Each CD31+ NC cell within L2 is paired with a RUNX2+ NC cell, a phenomenon not discernible within L1(Fig.1b,c). These cells retain their close spatial relationship later when CD31+ cells coalesce to form endothelia and RUNX2+ cells form an osteoblastic surface. The NC origin of the CD31+ endothelial cells is surprising as previous fate maps had posited that all cranial endothelia are mesodermal^12,13^. In fully developed spongy bone a significant fraction of (vWF+/CD31+) endothelia are NC in origin (white in Fig1d-f), while others are purely mesodermal or of mixed origin (Fig.1g,h). 3D analysis reveals that dermis vessels and rosettes already co-exist at E13 (fig.1i,j), However, the connection between α-SMA+ dermis vessels and bone vessels develops later in time - it is only discernable after E17 and mediated via rosettes. In 3D view a dermis vessel is seen connected to some rosettes, while other rosettes remain unconnected (Fig1l).

**Figure 1:**
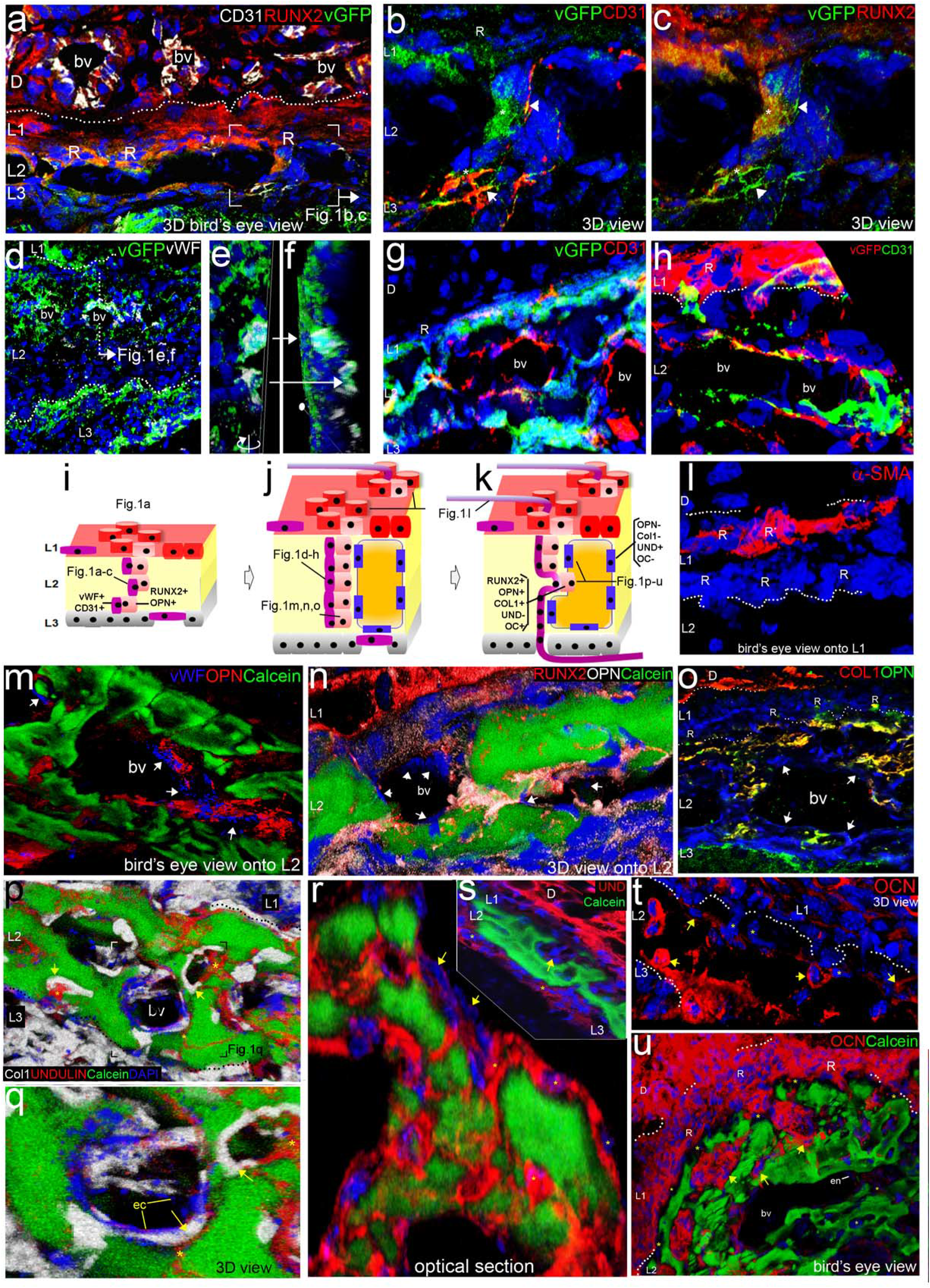
Emergence of the vascular-osteoblastic-mineral interface within L2. (**a**) Contrasting the dermis (D), which contains fully matured vasculature (blood vessels, BV), immature frontal bone (L1 & L3) comprises Runx2+ osteoblasts and individualised CD31+endothelial cells, the rosettes (R) are physically disconnected from dermis BVs. (**b**,**c**) Lacz+ neural crest-derived Runx2+ osteoblasts (*) and neural crest-derived CD31+ endothelia (arrow) enter the L2 space via rosettes (R) in a pair-wise manner. (**d-g**) Inside L2 the endothelia form BVs via vasculogenesis, and mature vasculature is of both neural crest (**d**, and cross sections **e**,**f**) and mesodermal (arrows in **g**) origin at E18. (**h**) The osteoblastic-endothelial interface, established by the spatial relationship of Runx2+ and CD31+ cells, is maintained even in the mature condition (E18). (**i-k**) Schematic of a detailed analysis of cellular identities during L2 maturation: osteoblast (pink) and endothelial cell (purple) pairs enter L2 via rosettes, inside L2 they form *de novo* sheets. Matrix (gold) is packaged into discrete mineral islands by a second cell population that is Runx2-/OPN-/Col1-/Undulin+ (blue). Once mineralised, areas are remodelled by Runx2+/OPN+/Col1+/Undulin-OBs. (**l**) A connection of the (α-SMA+) vasculature of the dermis and frontal-bone is established secondarily (*)post-birth (at P2) via rosettes. (**m-u**) Osteoblasts within mineralised (Calcein+) L2 are of two populations: immediately adjacent to the vasculature (BV) are OBs whose OPN is polarised towards the matrix (**m**). These cells are also Runx2+ (**n**) and Coll1+ (**o**). (**p-q**) OPN+/Runx2+/Coll1+ OBs (arrow) are a separate population from the Undulin+ OBs (*). (**r-s**) Undulin+ cellular extensions parcel the matrix (**r**, 3D reconstruction of several slices), while individual Undulin-cells invade the parcelled matrix (arrow in **s**). (**t-u**) These invading cells (arrow) are also OCN+ in the immature (**t**) and mature (**u**) frontal bone while the cells packaging the matrix are OCN-(*). **a-c, e-n, p-t**, 63-100x; **d**,**o** 10-20x. Nuclear DAPI, blue (all, bar **m**).

### Cell population diversity at the biomineral interface

We studied the cell population diversity of L2 *in vivo* at the molecular level and at single cell resolution. 3D reconstruction and volume rendering allowed us to ascertain whether a cell migrating through an environment containing a given factor also produces it.

Cells expressing OPN, an inhibitor of hydroxyapatite (HAP) crystal growth, partition vWF+ endothelia from the biomineral (Fig.1m). These OPN+ cells are also RUNX2+ (Fig.1n) and because OPN+ and Col1 immuno-reactivity is identical within L2 (Fig.1n,o), these three markers define a single dominant OPN+/RUNX2+/Col1+ osteoblastic cell type. Interestingly, a second cell type directly encases the mineral in more mature regions, This is Col1-negative but positive for Undulin (Fig.1p-s, asterisk), a FACIT collagen^14^ promoting adjacent Col1 motility^15^. The contiguous Undulin+ cellular sheet around the biomineral (Fig.1r,s) is broken locally by intercalating Runx2+/OPN+/Collagen1+/Undulin-osteoblasts (yellow arrows in Fig.1r,s,t) that are also positive for osteocalcin (OC) (Fig.1t,u) a known blocker of mineral nucleation^16^. UNDULIN and OC are mutually exclusive inside adjacent cells (Fig.1p,q).

### Osteoblasts secrete and decay two anisotropic collagen scaffolds

To understand how Runx2+/OPN+ NC cells could enact the observed biomineral shape changes we examined time courses of Collagen I (Fig.2a-f) and 2 (Fig.i-q) deposition and those of corresponding matrix metalloproteinases MMP9 (Fig.2e,f)^17^ and MMP13 (Fig.2l-q)^18^, instrumental cleavage of the respective collagens^19^.

**Figure 2:**
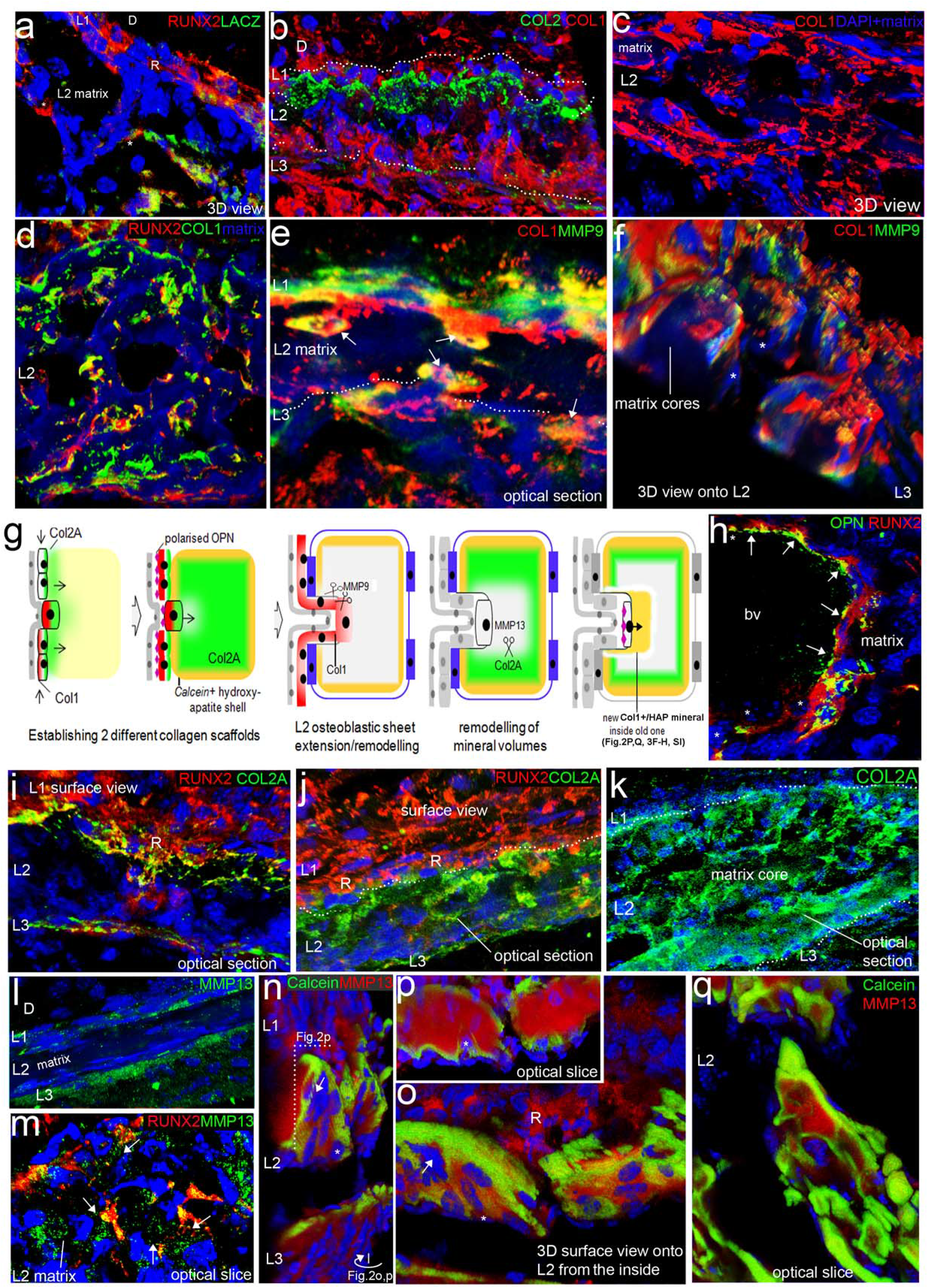
Runx2+ cells establish two anisotropic Collagen-MMP scaffolds for biomineral remodelling. (**a**) Runx2+ osteoblasts (*) dig into the L2 matrix, indicating they may have a role in remodelling. Matrix is laid along a collagen scaffold(**see ESM2**); (**b**) Collagen1 and Collagen 2 do not co-localize in 3D reconstructions of E16 frontal bone. (**c-d**) L2 matrix of mature frontal bone (evident by DAPI “sheen”) is not primarily Collagen 1-rich (**c**); instead Collagen1 is confined to areas along the borders of mineral and is expressed by cells that co-express Runx2 (**d**). (**e-f**) These Runx2+/Coll1+ osteoblasts dig into the mineralised matrix (arrows) by secretion of MMP9 which remains superficial to the MMP9-matrix core (*). (**g**) Schematic of the cellular deployment of Collagens and MMPs that facilitates the remodelling of previously established mineralized areas. (**h**) OPN+/Runx2+ osteoblasts at the border of vasculature (BVs) display polarised OPN (arrows), capping Calcein+ mineralization away from the BV lumen (see **ESM1**,**2**). (**i-k**) Runx2+ cells also secrete Collagen 2, which provides a progressively growing scaffold for the mineral matrix in the immature (**i-j**) and mature (**k**) frontal bone. (**l-m**) The Runx2+ cells also express and secrete MMP13 deep into the Collagen2+ matrix(arrow), rendering it malleable. (**n-q**) This results in a situation where MMP13 is found at the core of all mineralised areas, and invading cells are surrounded by a halo of MMP13 (arrow) which is not Calcein+. (**n-p**) 3D reconstructions (**n-o**) of area through which optical section **p** is taken, where a cell (*) can be seen depositing new calcein+ mineral (see ESM1.4) (**q**) The invasion and remodelling process results in intercalated shells of biomineral, seen in a cross section mature L2 (E18). **a-b**,**e-j**,**i-q** 63-100x; **c-d**,**k** 10-20x. Nuclear DAPI, blue.

Although COL1 is considered the dominant component of dermal bone matrix^20^, 3D analysis finds it confined to cellular sheets traversing L2, but not the surrounding L2 matrix itself (2a-c,f). MMP9, which cleaves fibrillar COL1^17^, occurs only in osteoblastic sheets (Fig.2e,f). The same Runx2/OPN+ NC cells (Fig.2a) secrete Col1 (Fig2d) and its modulator MMP9 (Fig.3 arrows).

Runx2+/OPN+ cells also produce COL2 in L1 and nascent L2 (Fig.2b,i-k) but not inside the D layer. Unlike COL1, COL2 spreads radially into the matrix with time, eventually occupying its entirety (Fig.2i-k). MMP13 cleaves Col2 with high affinity^18^ and is initially present only in L1 and L3 (Fig.2l) and later in all Runx2+ L2 cells (Fig.2n,m). Again, the collagen and its matching remodeller are made by the same cells (Fig.2m). Cells sitting on the surface and penetrating halfway into Calcein+ biomineral are each surrounded by a halo of MMP13 (Fig.2n-p asterisk), thus forming alternating and concentric collagen/mineral/MMP compound shells (2o, q). Such anisotropic protein deployment provides the first *in vivo* evidence for the hypothesis of collagen scaffold modulation as mechanism for biomineral remodelling (Fig.2g): While the Col2/MMP13 system can remodel biomineral volumes^21^, the Col1/MMP9 system appears to coordinate L2 sheet extension. As the c-axis of HAP crystals is aligned with the longitudinal axis of collagen I fibrils^22^, changes in fibril directionality change directionality of crystal growth (see ESM1,2). NC RUNX2+/OPN+ osteoblasts also co-express TRAF6 ^22,23^and RANK^24,25^, direct regulators of HAP crystal resorption lacunae and other behaviours previously attributed exclusively to osteoclasts (see ESM1.7). As there is no evidence of RUNX2+ NC-derived osteoclasts, TRAF6, RANK and MMPs should be reconsidered as agents of resorbing behavior, not as cell lineage markers.

We did not find evidence for OPN or COL1 inside endothelial cells. Instead, Runx2+/OPN+ cells exhibit polarized OPN localization towards the vasculature (Fig.2h) but COL1 facing the biomineral matrix (Fig.2c-f). Polarized OPN deployment prevents/caps biomineralization^26^ in between endothelia and osteoblasts and ensures the integrity of the endothelial-osteoblastic interface during the dramatic biomineral remodelling. This polarization is lost in *Hand2* mutant osteoblasts where inflationary L2 biomineral growth or vasculogenesis does not occur (ESM1.2;1.3 and accompanying paper), despite the presence of endothelial cells and osteoblasts. Given the role of HAND2 in L2 expansion (see accompanying paper), our 3D analysis reveals cell polarity and Collagen/MMP anisotropy as key mechanistic features of intercalary matrix remodelling by invading osteoblasts.

### Postnatal expansion of layer architecture by intercalation

In order to date biomineral and to trace intercalary growth into adult matrix histology we performed pulsed matrix dye injections *in vivo* and collected littermates at late developmental time points (P2,P8) (Fig.3A-H) (see ESM5).The first dye (green), injected at E14, permanently labels the earliest matrix (Jordan et al, accompanying Paper 1). The second dye (red) injection (at E18) stains biomineral deposited between E18 and P2. New biomineral deposited after P2 appears unlabelled (Fig.3A).

At P2 the biomineral deposited after E18 (red) sandwiches the earlier matrix (green) at the L1/L2 and L2/L3 interfaces (Fig.3B,C), consistent with late, scant apposition from L1 and L3 towards L2 (Fig.3f,l, asterisk in 3H), contemporary to intercalary matrix deposition (asterisk in Fig.3C,F-I). Red and green matrices are intersected radially by corridors of later (unlabeled) matrix (stippled line in Fig3BC), consistent with continuing invasion across layers. A surface view onto L1 at P2 shows osteoblastic rosettes (R in b-e) on top of circular collagen matrix, surrounding a central hole (arrow in 3E), providing evidence for a persistent generative architecture.

A littermate of the (E14 + E18 double) injection at P8 reveals significant corridors of unlabelled matrix, emanating from rosettes (* below R in f-h), transecting the earlier (green and red) labeled matrix of L2 (arrows in g,h) with cells sitting at their end (arrowheads in 3H). This provides evidence for continued invasive L2 remodelling into the postnatal period. These cells secrete matrix as we find GFP+ (NC) cells within L2 to be surrounded by secreted GFP+ membrane vesicles (ESM1.5).

At P8, L3 has also undergone a dramatic expansion of biomineral content and cellularity (Fig3B I). Our injections reveal parallel sheets of labeled matrix (red) alternating with later (unlabelled) matrix (3I,compare to uniformly red L3 matrix at P2, Fig.3B). Radial matrix tracts (arrows in 3I) communicate between these alternating sheets. They harbor migrating cells with perpendicular F-actin filopodia from E18 onwards (3J), providing evidence for continued invasive intercalary matrix deposition inside L3.

### Tracing inflationary bone growth into the gnathostome ancestry

Our generative model of inflationary dermal bone growth in mice (Fig.2G,3A) refutes the axiomatic model of appositional bone growth. We attempted to trace its evolutionary origin to the earliest vertebrate skeletons. Though fossil skeletal tissues preserve matrix structure at (sub-)cellular resolution, its growth can be difficult to discern without access to a spectrum of ontogenetic stages. Here, we studied growth stages of the Devonian placoderm *Bothriolepis canadensis* (Fig.3L,Fig 4A-F)– a representative early jawed vertebrate, and an Ordovician pteraspid heterostracan – among the earliest skeletonizing vertebrates (Fig.3M, 4G-L)^27^.

In *Bothriolepis canadensis*, the earliest stage in dermal skeletal development that we investigated comprises a distinct L1 (feature 1 in Fig.4), separated by a gap from the overlying D layer, that is superficial to a thin and poorly mineralized cancellous zone, above a compact L3 (feature 6,Fig.4A,B,C). Subsequent stages of development are observed between individuals and across single bones (Fig A vs B-M). L2 increases in thickness through the extension of vertical sheets of deposited matrix from L1 through L2 into L3 (feature4 Fig4C,D) , communicating across an emerging space just as in mouse. Large cancellae defined by vertical walls receive successive partitions, leading to a fretwork of connections without any visible direct connectivity to overlying dermis vasculature (Fig.4A,D). Only in those L2 areas with the most embellished fretwork do we find connections to dermis vessels that increase in size and number in more mature cancellous areas (Fig.3L).

In heterostracans, three dermal skeletal layers have been recognised historically^28^, but we here distinguish four distinct histological layers: a superficial D-layer of carrying odontodes, overlying a thin L1 (Fig.4G,H), an L2 cancellous/spongy layer, and an L3 plywood layer, all of which are acellular and all bar L1 possess a fibre-rich matrix (Fig.G-L). The initial stage of growth we infer encompasses just the D layer of odontodes, followed by L1, L3 (Fig.3M,4H) and, finally, L2 (Fig.4H)^29^ . L2 develops as in *Bothriolepis*, initially as a branching fretwork spanning L1 through L2 into L3, ultimately circumscribing polygonal cancellae; the branches are spanned subsequently by a fibre-rich matrix that establishes the discrete walls (Fig.3M,4I,J,K). L2 subsequently inflates in thickness through resorption and remineralization^30^. As ontogeny proceeds, cancellae were subdivided in the same way, by establishing branching pillars (Fig.4JI,J) that are spanned subsequently by a fibre-rich matrix (Fig.4H,feature 4). Simultaneously, L3 increases in thickness (feature6, Fig.4L).

The intrinsic ontogenetic fossil data provides *de facto* evidence of the same pattern of inflationary growth that we have observed in mouse and characterized molecularly, incompatible with apposition. We see the same expansion from a bi-layered to a trilayered architecture within the juvenile placoderm (3L) and heterostracan dermal bone matrix (from left to right) (Fig. 3M), capped by an (ossified fourth) D layer (Fig.4M).

**Fig.3.**
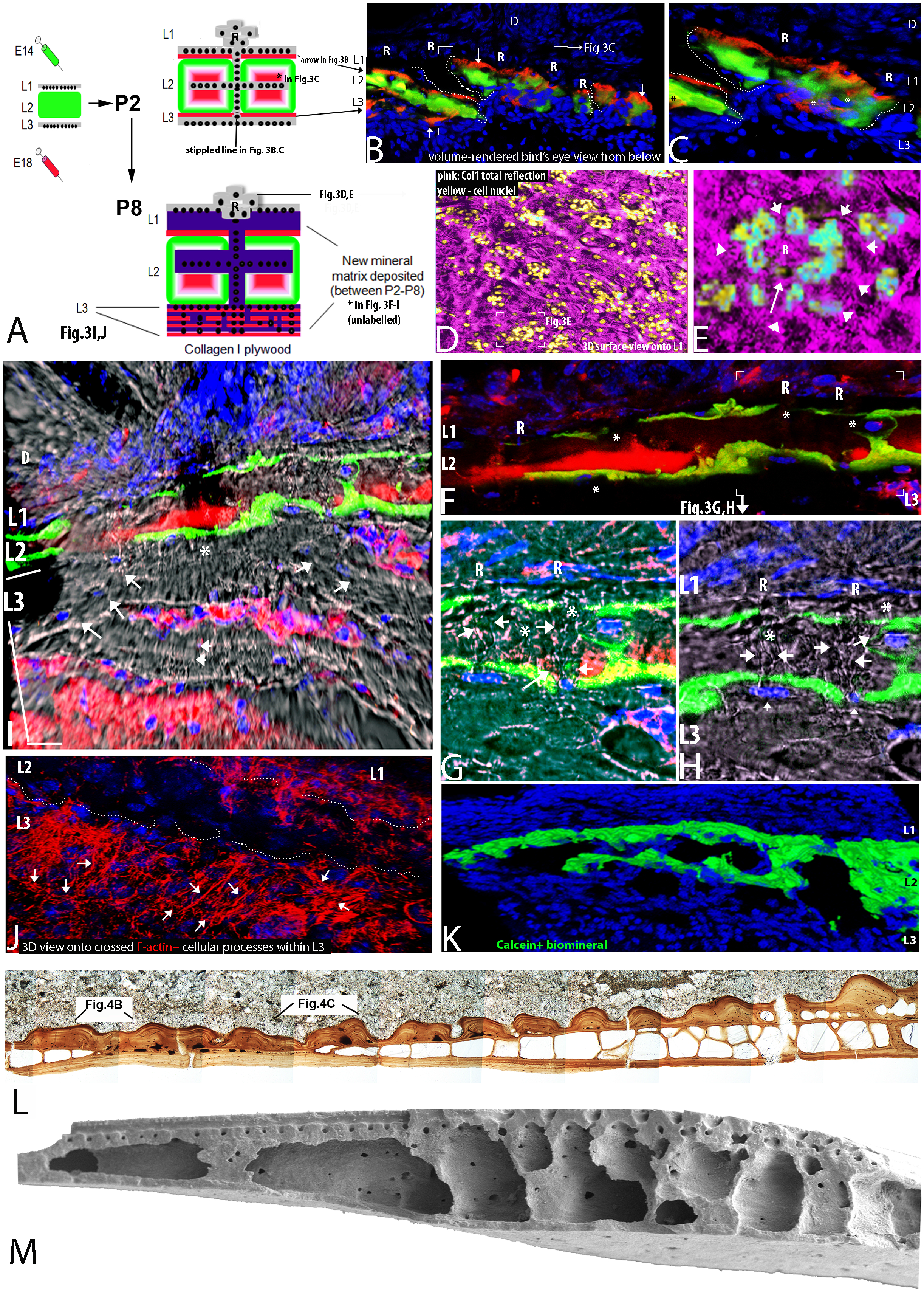
Biomineral remodelling in late mouse and early gnathostome bone ontogeny. (a-j) maintenance of the generative architecture and establishment of final biomineral histology. (**a)** *In vivo* double dye injection regime to label late (postnatal) biomineral deposition (see **ESM5**).(**b, c**) Biomineral deposited between E18 and P2 (labelled red) is incorporated at the core of the oldest mineral (green, *)and additionally encapsulates the entirety of the L2 biomineral (arrows), giving it the appearance of appositional deposition from L1 and 3. However, unlabelled area (b)deposited between P0 and P2 bisects older (green+red) mineral, indicating a continuing invasive process. (**d**,**e**) 3D surface view of flat-mounted calvaria reveal L1 rosettes as a persistent feature throughout P8. Circular birefringent collagen craters underlie each rosette, where a central matrix pore (arrow) is discernible.(**f**,**g**,**h**) Cellular invasion and intercalary matrix deposition continues (at least) til P8 with new matrix (born after P0) is nested within and replacing (red) labelled matrix (deposited between E18 and P0), compare to Fig.3**c**. **(g,h)** Birefringence patterns show late transmigration of individual osteoblasts (*) depositing unlabelled matrix (arrow). (**i**,**j**) Intercalary L3 matrix expansion. Unlabelled (later) matrix intercalates between labelled (red) matrix (compare to Fig. **b**,**c**) indicating a telescoping of L3 Col1 matrix which acquires a plywood histomorphology. This L3 plywood architecture is mirrored by the F-actin+ processes of invading cells (**j**). (**k-m**) Comparative layer morphogenesis and bimineral remodelling in mouse, placoderm and heterostracan dermal bone. k) change in calcein+ biomineral architecture along a growing dermal bone. (**l**) transition from a bi-layered to a tri-layered architecture in dermal bone of a juvenile antiarch placoderm *Bothriolepis canadensis*. (**m**) transition from a bilayered to a trilayered architecture in pteraspid heterostracan. Note in (**l**,**m)** independence of cancellar L2 emergence from overlying dermis vasculature. **b-c**,**e-h**,**j** 63-100x; **d**,**I**,**k** 10-20x. Nuclear DAPI, blue.

### Resolving the paradoxes of early vertebrate skeletal evolution

Knowledge of these processes underlying mouse dermal bone ontogeny provides new insights into the peculiarities of dermal skeletal tissues of the earliest skeletonizing vertebrates^31^ .

First, the honeycomb pattern of cancellar L2 wall formation (Fig.3L,M, 4G,J,K) does not reflect the overlying ridge-like pattern of dermis (D layer) vasculature in these fossils (Fig.3L,M, 4A,D,G). While this fossil evidence suggests independent processes governing L2 and D layer vascularization (Fig.4K), our findings in mouse of *de novo* L2 vasculogenesis and later connectivity to the D layer (Fig.1a-l) can explain this pattern.

Second, we re-discover the anatomically inconspicuous but ontogenetically essential L1 in fossils on the basis of (inwardly) polarized collagen1 secretion, entailing a HAP crystal structure gap at the D-L1 interface (feature 1) (see ESM2,3 for details).

Third, the cellular and tissue-level homology of the acellular aspidin) that comprises much of the dermal skeleton in heterostracans has been the subject of debate for almost two centuries. The radial and tangled arrangement of matrix fibres and fibre lacunae that occur in mineral layers surrounding the putative vascular spaces in L2 have been interpreted variably as acellular bone^32,33^, cellular bone^34^ and dentine^35^. The finding of invasive osteoblasts now provides a first interpretative framework.

Cells migrating across collagen1 matrix are known to organize fibrils along two axes, parallel and orthogonal to their migratory trajectory^36^. Tensile forces organize collagen in straps parallel to the migratory route (feature 2 in Fig.4M). We observe this feature in placoderms (Fig.4A,D,E) and heterostracans (Fig.4I,J), now interpretable as reflecting osteoblastic migration routes. Due to the non-linear properties of collagen meshes, a process called orthogonal mesh distortion leads to reorientation of collagen fibres perpendicular to the migratory direction (feature 3 in ESM2 and Fig.4M)^37^. It is these hallmarks of migration by COL1 secreting cells (Fig.2) that have been misinterpreted as cell lacunae or canaliculae in previous analyses of aspidin.

**Fig.4.**
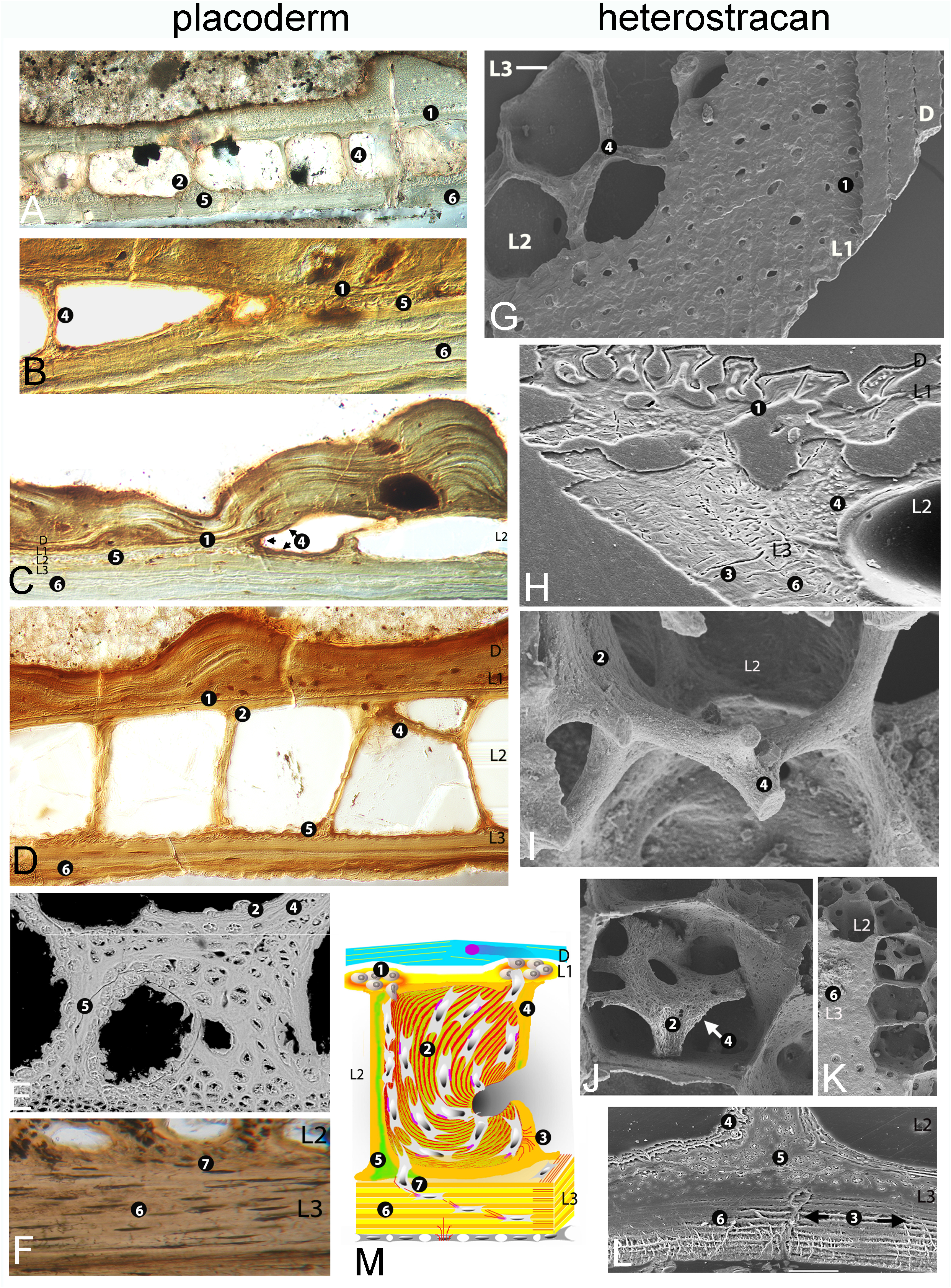
Invasive L2 growth and intercalary biomineralization of L2 and L3 in juvenile antiarch placoderm (*Bothriolepis canadensis*, **a-f**) and a pteraspid heterostracan (**g-l**), and shared generative architecture of dermal bone (**m**). **a**) overall architecture of immature dermal bone in section showing a superficial D layer underlain by an unmineralised L1 and, simple cancellous L2 and plywood like L3 at base. **b-c**) layer 2 expansion inside the same bone by establishment of radial clasps of biomineral (detail of 3L). Contiguity of matrix horizon traversing from L1 into L3 (arrows). **d**) L2 cancellar formation and subsequent horizontal partitioning, lack of vascular connections between D and L1/2, absence of cellularity inside the plywood L3 matrix **e**) L2 biphasic cancellar wall microhistology, consistent with anisotropic secretion of Collagen1 and 2 by Runx2+ cells (see Fig.2). **f**) late cellularity of L3 collagen plywood. **g)** overall architecture of heterostracan dermal bone in external view dissected to show superficial D-layer of odontodes (right), thin fibre-poor L1, underlying fibre-rich L2 cancellar layer and L3 plywood-like basal layer. **h**) transition from a bilayered L1/L3 to a trilayered L1/2/3 architecture. **i**) branching morphogenesis within L2 layer elaboration. (**j-k**) newly forming horizontal cancellar walls (arrow fibrous collagen1), **l**) biphasic L2 walls - sheet-like Collagen1 vs amorphous Col2/Col1 mixtures, as well as L3 isopedine and radial collagen straps indicating orthogonal migratory direction. **m**) Schematic representation of early mouse, juvenile placoderm and adult heterostracan dermal bone anatomy at early stages on the basis of anisotropic Collagen1 (yellow/orange) and Collagen2 (green)secretion by invading osteoblasts. The partnering endothelial cells (see Fig.1) are omitted for clarity. 7 ultrastructural predictions can be made from our extant mouse data that are testable in fossils: **1**.D-L1 Collagen1 matrix discontinuity due to polarized Col1 secretion inside L1 towards L2, this is found in Bothriolepis (A-D) and the pteraspid (j,k,l).**2**. collagen fibre strap formation within L2, found in placoderm (e) and heterostracan (i,o, r) as indicators of the migratory direction of fibripositor (pink) carrying, Collagen secreting (Fig.2) osteoblasts. **3**.Radial fibre orientation as migration marker orthogonal to the direction of migration (e,o,r), **4**.early cancellar walls primarily made of Col1 (Fig.2)deposited in sheets (c,e,i,n,m), **5**. Later biphasic collagen1/collagen2 architecture (see Fig.2 for details)(f,q,r).**6**.Plywood self-organization as hallmark of high concentration of Col1 in layers 2,3 (b,c,h,q,r,s)of invading cells. **7**.ontogenetically later invasion of L3 by cells in mice (Fig.3a-j) and placoderm (i.e. animals with cellular L2 bone). Compare difference in cellularity between early (cell free L3, in c) and late (s)L3 in Bothriolepis, while the collagen1/isopedine sheets of the heterostracan remain acellular. See ESM3,4 for details.

Fourth, the biphasic state of heterostracan (Fig.4G,H,I,L) and placoderm (Fig.4A,D,E) cancellar wall matrix (features 4,5) develops from sheet-like ordered (feature4), to a homogenous core with ordered margins (feature 5). This mirrors the biphasic and anisotropic nature of collagens seen in mouse dermal bone (Fig.2). While early thin cancellar walls are only made of Col1 (confined to L2’s cellular sheets, Fig.2B,C), with time Col2 penetrates the L2 matrix (Fig.2i-k) resulting in homogeneous cores of cancellar walls, bounded by Col1 sheets (feature 5 in Fig.4L).

Fifth, a plywood-like arrangement of Collagen fibres has been described (feature6) and is visible in placoderm (Fig. 4A-F), heterostracan (Fig. 4G-L) and mouse (Fig.3I) L3. Our pulse labelling shows that the stratified appearance of the L3 collagen does not reflect appositional growth. Pure COL1 at very high concentration can self-organize as cholesteric liquid crystals with plywood architecture *in vitro*^38^. We describe high expression of Col1 in L3 cells *in vivo*, which is even retained in HAND2 mutants that do not elaborate L2 (see Fig. 2B here, ESM and Fig. 3J,K,L of accompanying paper). This provides the most parsimonious explanation for the fossil plywood architectures.

Sixth, our ontogenetic comparison reveals an important difference: No cellular lacunae become visible during heterostracan L3 plywood ontogeny (Fig. 4H,L). In contrast, we see an increase in the number of such spaces during placoderm L3 ontogeny (Fig. 4A,C,D versus F). This is consistent with persisting intercalary growth of L3 plywood and the evolutionary emergence of cellular bone (see node 3 in Fig. 5) from a common acellular bone matrix (feature7)^27^. Even in the mouse the polarized osteoblastic-endothelial interface (Fig.2) creates acellular bone matrix in the first place before osteoblasts become surrounded by matrix, forming ‘cellular bone’ later (see ESM4).

**Fig. 5:**
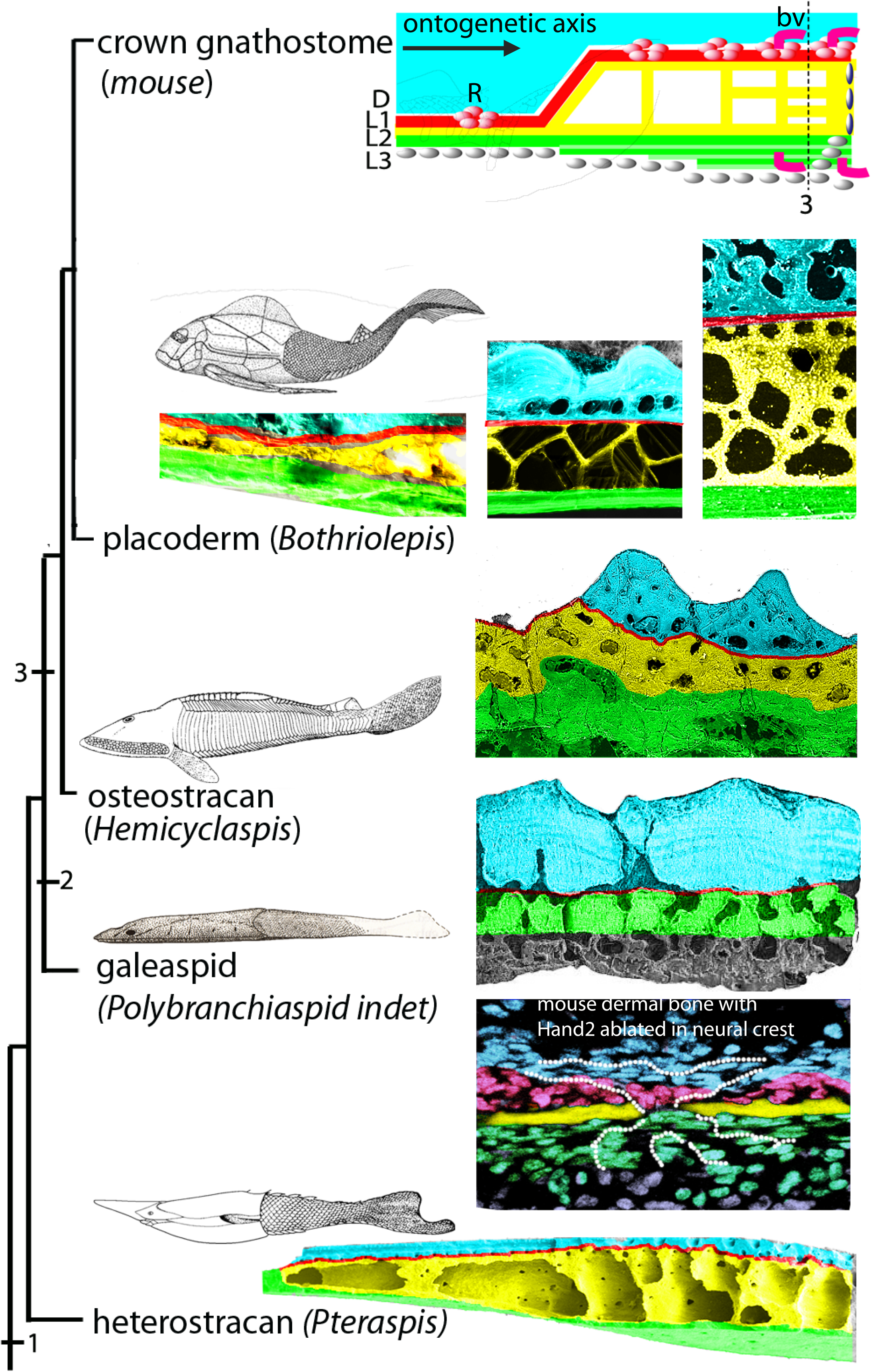
Phylogeny of generative modularity across gnathostome dermal bone development. All species examined here (mouse, juvenile placoderm and adult heterostracan) display a trilayered (L1,2,3) architecture underneath a D layer (blue) and a strikingly similar ontogenetic trajectory. As a well-developed L2 is discernible in heterostracans (Fig.4), this finding phylogenetically brackets all skeletonising vertebrates. While the developmental trajectory of layer elaboration is conserved across total group gnathostomes (left to right), the extent of individual layers differs. Node 1: Origin of dermal bone comprising D (turquoise) and L1-3 (red, yellow,green). Node 2 inferred loss of L2 elaboration in galeaspids. For comparison, *Hand2* neural crest ablation in mouse (see accompanying paper and ESM1,2) abolishes L2 (yellow) expansion and is remarkably similar to a galeaspid morphology. Node 3 appearance of cellular bone, i.e. L2 and L3 cells fully encased by matrix, cellular invasion into L3 matrix. See ESM3 for detailed layer identification criteria in these fossil taxa.

Thus, the striking set of similarities in ontogenetic micro anatomy between mouse, placoderm and heterostracan suggests that the earliest stages of mouse development replicate the juvenile ontogenetic development in members of the gnathostome stem group and indicate a shared histogenetic mechanism that evolved in the earliest skeletonizing vertebrates (Fig. 5, node1). We surveyed the broad phylogenetic spectrum of early jawless and jawed vertebrates (see ESM5), and found that the structure of the dermal skeleton can be rationalized on this same pattern of inflationary growth, independent of the processes that pattern the odontodes in the D-layer. The different parts of the generative architecture are expected to be embellished differently in different groups. For example, L3 expands while L2 elaboration is evolutionarily lost in some clades, such as galeaspids (Fig.5, node2). Their skeletal structure compares well to the phenotype of ablating *Hand2* (or its branchial arch enhancer) in mouse NC, reflecting a genetically modular mechanism.

## Discussion

We describe for the first time the mechanism underlying intercalary biomineralization. The juxtaposition of osteoblasts and endothelia underneath rosettes by common descent (impossible to detect without genetic mosaic labeling) is retained in the adult biomineral-endothelial interface and enables trophic interactions. Independent L2 vasculogenesis ensures that pressure gradients between dura and dermis vessels cannot interfere with L2 histogenesis. Biomineral growth is predicated upon the remarkable ability of OPN+/nRUNX2+ osteoblasts to erect and erode bone matrix. This ability predates ontogenetically and phylogenetically the osteoclasts of the megakaryocyte lineage^39, 40^ that likely emerged with the evolutionary invention of endochondral ossification and bone marrow.

This has significant biomedical and evolutionary implications. Runx2 mutants lack dermal bone^41^. Surprisingly, (PTH) hormone mediated^42^ or Col1-driven^43^ over-expression of RUNX2 does not lead to more bone, as might be expected, but to increased bone erosion with time. We can now resolve this conundrum by showing that biomineral erosion is key mechanism for biomineral expansion through intercalary growth. Runx2 mediates cancer osteomimicry^44^ and its target genes MMP9/13 are part of the invasive programme entailing tumour vasculogenesis^19,45,46^. This might suggest a secondary re-activation of an earlier osteogenetic programme in pathological contexts. Osteoarthritis and osteoporosis might similarly represent reactivations of erosive programmes among resident osteoblasts. Future therapies could benefit from directly targeting these nRUNX2+/OPN+ cells in addition to targeting their interactions with osteoclasts: dermal bone might serve as an interesting model system. *In vitro* bioengineering of dermal bone remains a significant challenge. Our 3D time course analysis reveals osteoblasts erecting highly anisotropic 3D collagen1/2/MMP scaffolds when building up the spongy layer. Biomimetics stimulating such generative architecture and anisotropy might be an attractive new avenue for bone engineering.

Inconsistencies in dermal tissue structure and growth between living and fossil vertebrates have been interpreted to reflect an early phase in the evolutionary establishment of this system, antecedent to a more canalised system present in living jawed vertebrates^31^. However, these inconsistencies were based on an unsubstantiated model of apposition alone. Quite to the contrary, the earliest skeletonizing vertebrates exhibit a pattern of spongiosa growth that is entirely compatible (down to the ultrastructural level) with the inflationary process encountered in living jawed vertebrates such as mouse. This suggests a very rapid establishment of these processes in the deep evolutionary history of gnathostomes. In combination with the D layer odontode theory, our mechanistic model serves as a first comprehensive framework for dermal skeletal development and evolution.

## METHODS SUMMARY

See and accompanying paper (Jordan et al.) for immunohistochemistry, image acquisition and analysis. To enable us to trace the cytoplasmic outlines of neural crest cells, we generated a novel recombinase reporter strain with a membrane bound vGFP (to be published in detail elsewhere). Biomineralization was explored using histological addition to calcein *ex vivo* in combination with multiplex immunohistochemistry (at 5mg/mL concentration), and *in vivo* via labelling experiments conducted under home office licence PPL 70/7178. To look at late aspects of biomineralization, intraperitoneal injections of each labelling agent were conducted on C57BL/6J mice from E11-E18 (with a minimum two day gap between injections) at the following concentrations: calcein 10mg/kg and xylenol orange 90 mg/kg; specimen were isolated for analysis from P2 – P8. Rosette architectures along the surface of L1 were examined in dissected calvaria where the frontal and parietal bones were flat-mounted without removal of the overlying dermis, imaged through their depth by confocal microscopy and reconstructed in 3D segments. Please refer to ESM5 for further details.

The antiarch placoderm *B*.*canadensis* was chosen because it is representative of the diverging lineage of jawed vertebrates^47,48^, its dermal skeleton is representative of ‘placoderms’ generally^49,50^, and because uniquely preserved ontogenetic stages are available. A pteraspid heterostracan from the Canadian Arctic was chosen due to its pristine tissue preservation of ontogenetic stages and its phylogenetic position, bracketing the majority of skeletonized vertebrates.

## Supporting information

Jordan2 Supplements

## Acknowledgements

Dedicated to the memory of Professor Farish A. Jenkins, gentleman, explorer of lost vertebrates, inspiring teacher. We thank Andrew Lumsden, Nick Dale, Jonathan Millar for most helpful comments on the manuscript, Dan White for technical assistance with BioImageXD, Samantha Dixon and Ian Bagley for expert animal husbandry and care, Ian Portman for imaging support. We thank Sylvain Desbiens and Johanne Kerr (MHN Miguasha) for materials and Marilyn Fox, Jay Ague, Ruth Blake, James Eckert, Shun-ichiro Karato, and Danny Rye (Yale Peabody Museum and Department of Geology) for lab access. We gratefully acknowledge grant funding by HFSPO (RGP0029/2007-C), the Wellcome Trust and ConquerChiari (all awarded to GK), NIH R01HD0648240 (HY) NIH grant DE018899 (DEC), NERC NE/G016623/1 (PCJD), NSF, Geological Society of America, Paleontological Society and Yale University (JPD).

## Author contributions

KWJ and GK designed all experiments and wrote the manuscript jointly with PCJD. KWJ performed and analyzed all experiments. DEC and HY and XZ generated and provided essential transgenics. JD and PCJD performed the ultrastructural histological analysis of fossils. GK and PCJD integrated information across fossil and extant data. All authors contributed towards data analysis and approved the final manuscript.

## Author information

The authors declare no competing financial interests. Correspondence and requests for extant material should be addressed to G.K. and fossil work to PCJD.

## References

1. Williamson, W. C. Philosophical Transactions of the Royal Society of London, Series B Investigations into the structure and development of the scales and bones of fishes. 141, 643–702 (1851).

2. Klaatsch, H. Morphologisches Jahrbuch Zur Morphologie der Fischschuppen und zur Geschichte der Hautsubstanzgewebe. 16, 97–196, 209-258 (1890).

3. Goodrich, E. S. Proceedings of the Zoological Society, London On the scales of fish, living and extinct, and their importance in classification. 2, 751–774 (1907).

4. Ørvig, T. in Problems in vertebrate evolution eds S. M. Andrews et al.) 53–75 (Linnean Society Symposium Series 4, 1977).

5. Reif, W.-E. Evolutionary Biology Evolution of dermal skeleton and dentition in vertebrates: The odontode regulation theory. 15, 287–368 (1982).

6. Stensiö, E. A. in Geology of the Arctic Vol. 1 (ed G. O. Raasch) 231–247 (University of Toronto, 1961).

7. Bystrow, A. P. Morphologische untersuchungen der Deckknochen des Schaedels der Wirbeltiere. Acta Zoologica 16, 65–141 (1935).

8. Witzmann, F., Scholz, H., Mueller, J., Kardjilov, N. Sculpture and vascularization of dermal bones, and the implications for the physiology of basal tetrapods Zoological Journal of the Linnean Society, 2010, 160, 302–340

9. Donoghue, P. C. J. & Sansom, I. J. Microscopy Research & Technique Origin and early evolution of vertebrate skeletonization. 59, 352–372 (2002).

10. Morris-Kay, G., Wilkie, A. O.M. (2005) Growth of the normal skull vault and its alteration in craniosynostosis: insights from human genetics and experimental studies. J.Anat. 207, 637–653.

11. Witzmann, F. Comparative histology of sculptured dermal bones in basal tetrapods, and the implications for the soft tissue dermisPalaeodiversity 2: 233–270; Stuttgart, 30.12.2009

12. Couly, G., Coltey, P., Eichmann, A., & Le Douarin, N. M. (1995). The angiogenic potentials of the cephalic mesoderm and the origin of brain and head blood vessels. Mechanisms of development, 53(1), 97–112.

13. Morriss-Kay, G. M. (2001). Derivation of the mammalian skull vault. J Anat 199(Pt 1-2): 143–151.

14. Boot-Handford, R.P. et al. 2003. A Novel and Highly Conserved Collagen (pro_1(XXVII)) with a Unique Expression Pattern and Unusual Molecular Characteristics Establishes a New Clade within the Vertebrate Fibrillar Collagen Family, JBC, Vol. 269. No. 45, 28193–28199,

15. Nishiyama et al. 1994. Type XII and Xiv collagens mediate interactions between banded collagen fibers in vitro and may modulate exracellular matrix deformability , JBC, Vol 269, 11. ,pp 28193–28199

16. Hoang et al. 2003. Bone recognition mechanism of porcine osteocalcin from crystal structure, Nature 425, 977–980.

17. Bigg, H. F., Rowan, A. D., Barker, M. D., & Cawston, T. E. (2007). Activity of matrix metalloproteinase-9 against native collagen types I and III. FEBS Journal, 274(5), 1246–1255.

18. Behonick, D. J., Xing, Z., Lieu, S., Buckley, J. M., Lotz, J. C., Marcucio, R. S.,. & Colnot, C. (2007). Role of matrix metalloproteinase 13 in both endochondral and intramembranous ossification during skeletal regeneration. PLoS One, 2(11), e1150.

19. Ortega, N., Behonick, D. J., & Werb, Z. (2004). Matrix remodeling during endochondral ossification. Trends in cell biology, 14(2), 86–93.

20. Karsenty, G., Kronenberg, H.M., Settembre, C. (2009). Genetic Control of Bone formation. Annual Rev. Cell Dev. Biolo. 209. 25: 629–648.

21. Kikuchi et al. 2004. Biomimetic synthesis of bone-like nanocomposites using the self-organization mechanism of hydroxyapatite and collagen. Composites Science and Technology 64 , 819–825

22. Kim et al. 2005. Osteoclast differentiation independent of the TRANCE–RANK -TRAF6 axis. Journal of Experimental Medicine, Vol. 202, No. 5,. 589–595

23. Kobayashi et al. 2001. Segregation of TRAF6-mediated signaling pathways clarifies its role in osteoclastogenesis. EMBO Journal Vol. 20 . 6, 1271–1280

24. Asagiri, M. & Takayanagi, H. 2007. The molecular understanding of osteoclast differentiation, Bone 40, 251–254

25. Rubin et al. 2005.: Osteoclast: Origin and Differentiation , Bone Resorption , Topics in Bone Biology Volume 2, 1–23

26. George A. , Veis, A. 2008. Phosphorylated Proteins and Control Over Apatite Nucleation, Crystal Growth and Inhibition, Chem Rev. 2008 November ; 108(11): 4670–4693.

27. Janvier, P. 1996. Early Vertebrates, Oxford University Press.

28. Huxley, T. H. Quarterly Journal of the Geological Society, London On Cephalaspis and Pteraspis. 14, 267–280 (1858).

29. Denison, R. H. Palaeontographica (Abt. A) Growth and repair of the shield in Pteraspididae (Agnatha). 143, 1–10 (1973).

30. White, E. I. Palaeontographica Abt. A Form and growth in Belgicaspis (Heterostraci). 143, 11–24 (1973).

31. Smith, M. M. et al. Modern Geology ‘Teeth’ before armour: the earliest vertebrate mineralized tissues. 20, 303–319 (1996).

32. Agassiz, L. Recherches sur les Poissons Fossiles. (Imprimerie de Petitpierre, 1833–43).

33. Ørvig, T. Arkiv för Zoologi Palaeohistological notes 2: certain comments on the phylogenetic significance of acellular bone in early lower vertebrates. 16, 551–556 (1965).

34. Halstead Tarlo, L. B. Nature Aspidin: the precursor of bone. 199, 46–48 (1963).

35. Denison, R. H. Fieldiana Geology Ordovician vertebrates from Western United States. 16, 131–192 (1967).

36. Petroll WM et al. 2004. Dynamic Three-Dimensional Visualization of Collagen Matrix Remodeling and Cytoskeletal Organization in Living Corneal Fibroblasts, Scanning VOL. 26, 1–10.

37. Sawhney R & Howard J. 2002 Slow local movements of collagen fibers by fibroblasts drive the rapid global self-organization of collagen gels The Journal of Cell Biology, Volume 157, Nr. 6, 1083–1090.

38. Giraud-Guille et al. 2003. Liquid crystalline assemblies of collagen in bone and in vitro systems. Journal of Biomechanics 36, 1571–1579

39. Geissmann, F., Manz, M. G., Jung, S., Sieweke, M. H., Merad, M., & Ley, K. (2010). Development of monocytes, macrophages, and dendritic cells. Science, 327(5966), 656–661.

40. Asagiri, M. & Takayanagi, H. 2007. The molecular understanding of osteoclast differentiation, Bone 40, 251–254.

41. Otto et al. (1997).Cbfa1, a Candidate Gene for Cleidocranial Dysplasia Syndrome, Is Essential for Osteoblast Differentiation and Bone Development, Cell ,Vol. 89, 765–771,

42. Bellido, T., Ali, A. A., Plotkin, L. I., Fu, Q., Gubrij, I., Roberson, P. K., … & Jilka, R. L. (2003). Proteasomal degradation of Runx2 shortens parathyroid hormone-induced anti-apoptotic signaling in osteoblasts: a putative explanation for why intermittent administration is needed for bone anabolism. Science Signalling, 278(50), 50259.

43. Geoffroy, V., Kneissel, M., Fournier, B., Boyde, A., & Matthias, P. (2002). High bone resorption in adult aging transgenic mice overexpressing cbfa1/runx2 in cells of the osteoblastic lineage. Molecular and cellular biology, 22(17), 6222–6233.

44. Akech, J. et al. 2010. Runx2 association with progression of prostate cancer in patients: mechanisms mediating bone osteolysis and osteoblastic metastatic lesions, Oncogene 29, 811–821.

45. Ahn, GO and Brown, JM. 2008 Matrix metalloproteinase-9 is required for tumor vasculogenesis but not for angiogenesis: Role of bone marrow-derived myelomonocytic cells Cancer Cell. March ; 13(3): 193–205

46. Chou, J., Lin, J. H., Brenot, A., Kim, J. W., Provot, S., & Werb, Z. (2013). GATA3 suppresses metastasis and modulates the tumour microenvironment by regulating microRNA-29b expression. Nature Cell Biology, 15(2), 201–213.

47. Brazeau, M. D. Nature The braincase and jaws of a Devonian “acanthodian” and modern gnathostome origins. 457, 305–308 (2009).

48. Davis, S. P. et al. Nature Acanthodes and shark-like conditions in the last common ancestor of modern gnathostomes. 486, 247–250 (2012).

49. Downs, J. P. & Donoghue, P. C. J. Journal of Morphology Skeletal histology of Bothriolepis canadensis (Placodermi, Antiarchi) and evolution of the skeleton at the origin of jawed vertebrates. 270, 1364–1380 (2009).

50. Giles, S. et al. Journal of Morphology Histology of “placoderm” dermal skeletons: Implications for the nature of the ancestral gnathostome (2013).

