## Supplementary material for "Dermal bone remodelling by invasive osteoblasts evolved in stem gnathostomes": Jordan2 Supplements

#### Supplemental materials:

Jordan et al.. Osteoblastic dermal bone remodelling is a stem gnathostome feature.

##### 1. Generative model of dermal bone formation and its ultrastructural consequences

- 1.1. A novel highly invasive osteoblastic cell type
- 1.2. Hand2 dependent cellular polarity of osteopontin secretion
- 1.3. A common origin of endothelia and osteoblasts
- 1.4. Relationship between this invasive cell type and biomineral features
- 1.5. Intercalary matrix secretion
- 1.6. Hydroxyapatite crystals follow Collagen patterns
- 1.7. Other novel resorptive molecular features of Runx2+ osteoblasts within L2.

##### 2. Ultrastructural predictions of collagen anisotropies – usable in extant and fossil taxa

###### 2.1. Biomineral features:

1. D-Layer-L1 discontinuity of matrix
2. Collagen fibre strap formation within L2.
3. Radial fibre orientations underneath migrating cells - orthogonal amplification of collagen mesh distortion.
4. Early Collagen1 composition of cancellar walls
5. Later Collagen1/collagen2 biphasic (sheetlike-amorphous) architecture of cancellar walls
6. 7. Plywood self-organization of Collagen1 in layer 3 and 2
- Later cellular invasion of L3 and intercalary matrix growth of the compacta.

##### 3. Reconciling growth of the dermal skeleton in the earliest skeletonising vertebrates

Heterostraci

Galeaspida

Osteostracans

Placodermi

##### 4. Acellular versus cellular L2 biomineral in ontogeny and evolution

##### 5. Detailed Methods

### 1. Generative model of dermal bone formation and its ultrastructural consequences

In the accompanying paper we use mosaic analysis, genetic fate mapping, 3D reconstruction at single cell level in combination with in vivo labelling of nascent matrix in order to understand radial growth of dermal bone in the mouse. We redefine generative layers, find molecular evidence for a D, L1 separation, L2, L3 and observe that while L1 and L3 stay contiguous, it is L2 which grows substantially in thickness. Our findings are not compatible with a standard appositional model but necessitate growth of L2 and L3 by an invasive cellular and intercalary biomineral deposition process.

#### 1.1. A novel highly invasive osteoblastic cell type

In the present paper we reveal key molecular aspects of the cell populations at work in invading and remodelling layer 2 and 3, while layer 1 is retained in morphology and molecular anatomy throughout. These invading cells are characterized by nuclear Runx2 and Hand2 immunoreactivity. Abzhanov et al. purifying cells from biomineral and then performing double in situ hybridization were detecting a Col1+/Col2+ cell type, which they termed CLO cells (Abzhanov et al, 2007). However as only RNA in situ hybridization was used in isolated cells, the anisotropic collagen1/2 distributions that we discover in vivo could not be discovered nor was it clarified whether Runx2 and OPN can coexist inside the same cells. We clarify here that they do (see Fig. 2f,g,h accompanying paper, Fig. 2 this paper).

Runx2 and its downstream target genes MMP9 and 13 (**Fig. ESM11,12**) are part of a generic invasive signature used in other regions of the body (during osteomimicry of breast/prostate cancers, Jimenez et al 1999, Akech et al 2010 or in carcinomas Ahn and Brown 2008). Our findings imply that such signature might have evolved in the phylogenetically older context of dermal bone development and is being reactivated in the context of tumorigenesis.

These Runx2+ cells also carry Periostin (POSTN), member of the ancient Fasciclin II family which is responsible for sheet migration (Lindsley et al., 2004) (as is Hand2 which directly controls POSTN, Holler et al. 2010). Moreover, periostin itself blocks biomineralization (Saito et al, 2002). Their neighbouring cells within L2 are positive for Undulin (Collagen XIV of the FACIT family) which facilitates sliding of the Collagen1 cellular sheets of in relation to the biomineral that is encased by Collagen XIV+ cells (**Fig ESM03, 05**) (Boot-Handford et al. 2003). This is inferred from the known function of undulin to mediate extracellular matrix deformability (Nishiyama et al. 1994).

Most notably, osteopontin (OPN) and Osteocalcin (both previously considered to be 'late' osteoblastic markers) (Abzhanov et al. 2007) are found on the Runx2+ cells at the earliest stages of L2 formation (**Fig ESM02,04,06**). Both OPN and OC cap hydroxyapatite crystals (Hunter et al. 1994, George and Veis 2008, Ge et al. 2003, Hoang et al. 2003). This now places for the first time the *in vitro* functionality of OPN and OC in capping biomineralization into an *in vivo* context. Thus the polarized secretion of OPN (away from the biomineral, in between osteoblasts and emerging blood vessels/endothelial cells) that we observe here ensures continued communication between endothelia and osteoblasts and their nourishing relationship (**Fig ESM06 left and right**), while OPN and OC both keep the invasive layer within L2 malleable and prevent that cells will become firmly encased prematurely – a phenotype found inside the small thin sheet of biomineral found in Hand2 mutants (Fig.4m,n,o of the accompanying paper, **FigESM08**). At early stages of L2 elaboration OC+/Runx2+/OPN cells are in direct contact with the biomineral front and no Undulin+ cells cover the biomineral (**Fig ESM01,02**).

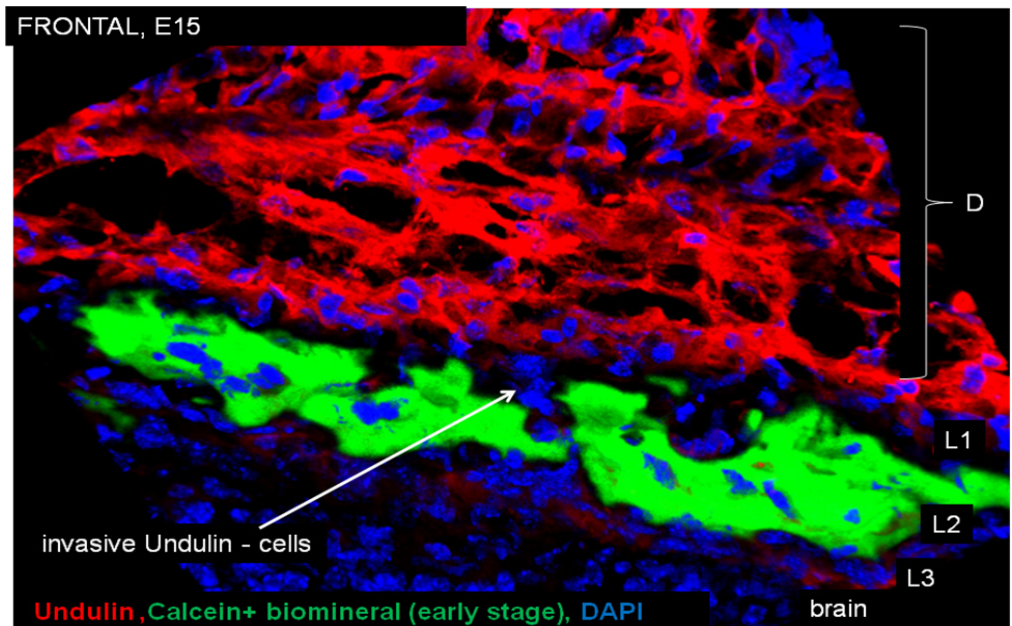

**Fig ESM01**

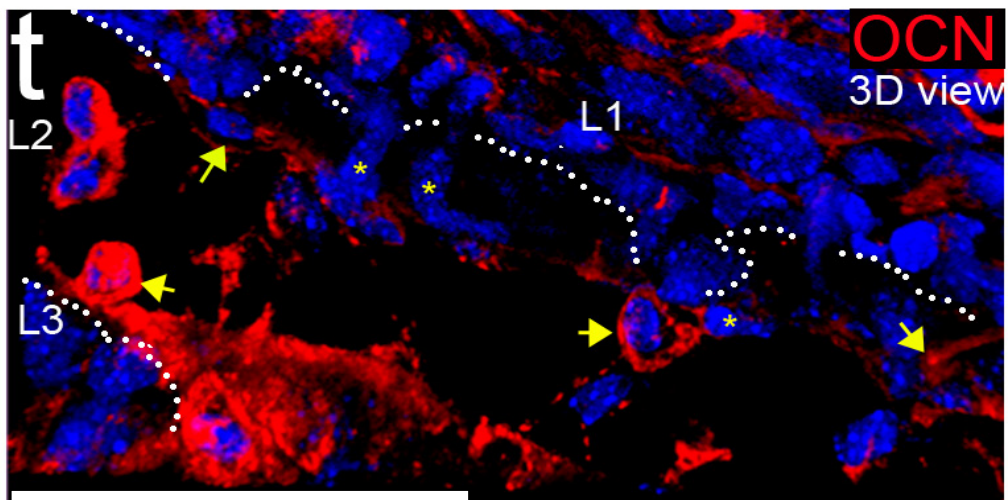

**Fig ESM02**

The two figures (**FigESM01 and 02**) above represent stage i) below, prior to the emergence of the Undulin+ cells (blue in j,k of **FigESM03** below) encasing the biomineral (orange).

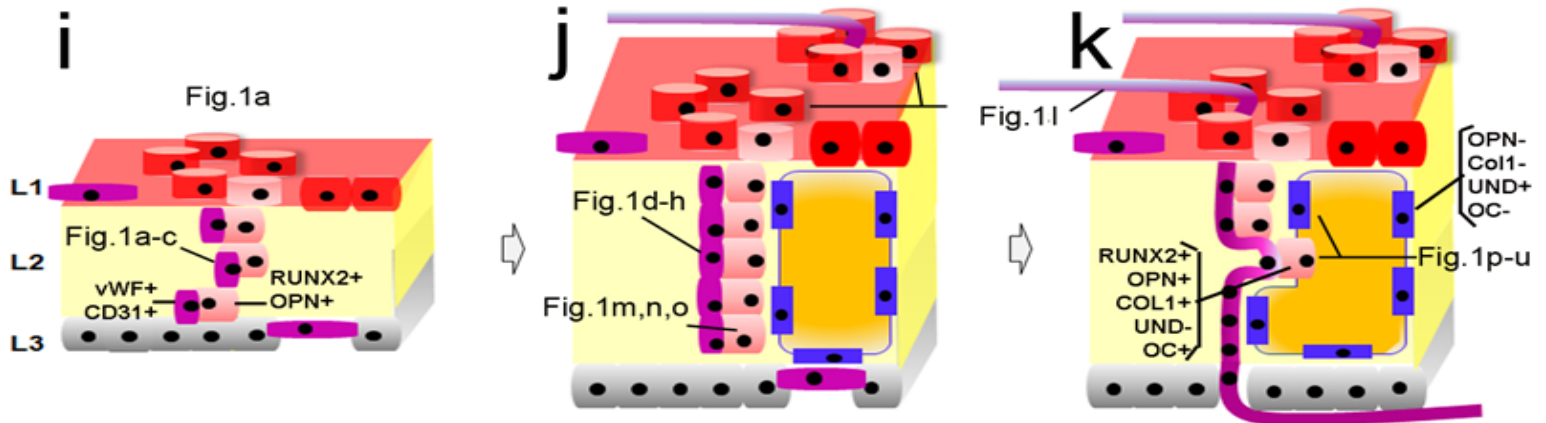

**Fig ESM03 – emergence of the undulin+ population**

The following two figures (**FigESM04,05**) represent state j,k: a thin sheet of OC negative (Undulin+ cells, yellow arrows) in between the OC+invasive osteoblasts (see R for rosettes) and the biomineral (green) on the other side (**FigESM04**).

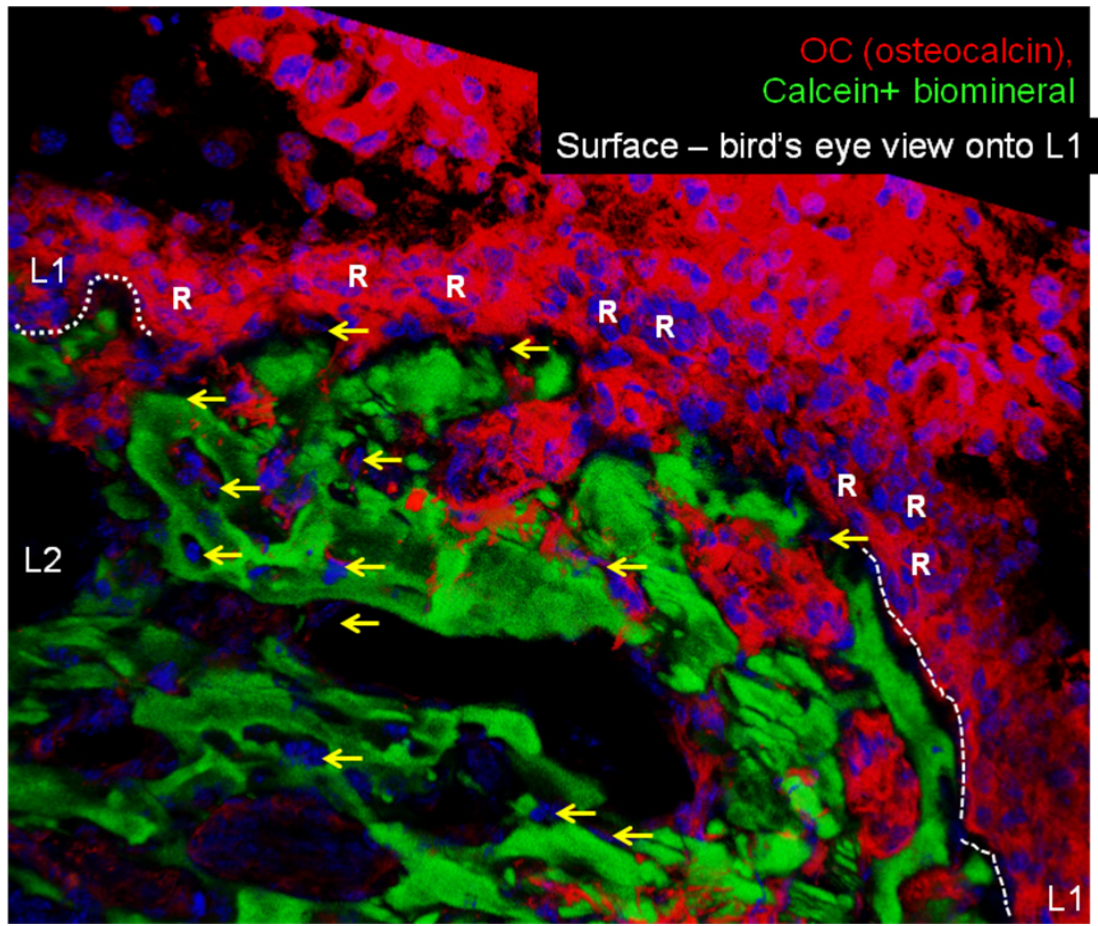

**Fig ESM04. Invasive osteocalcin+ cells**

Complementarily to that we find Undulin+ cells directly surrounding and encasing the biomineral (see previous manuscript and **FigESM05** below).

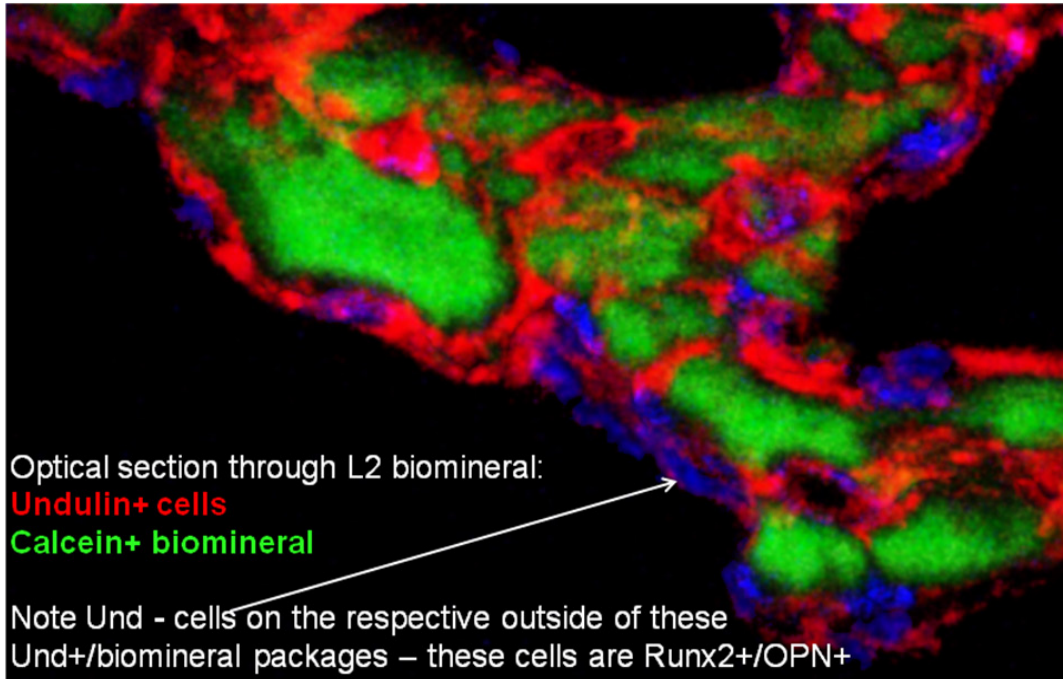

**Fig ESM05 – Undulin+ sheets around more mature (cancelled) biomineral**

#### 1.2.Hand2 dependent cellular polarity of osteopontin secretion

As we describe in the accompanying previous paper, Hand2 mutant (neural crest) cells lose the ability to polarize, remain blast-like and, consequently, all cells become positive for nuclear Runx2 and OPN has lost its polarization in such blast-like cells and fully surrounds these cells. Compare in **Fig ESM06** below the OPN+ polarity in flattened Runx2 wild-type cells (left side) with the circumferential OPN immunoreactivity in Hand2 mutant cells (LacZ+, red below) that display a blast-like morphology (right side of **FigESM06**).

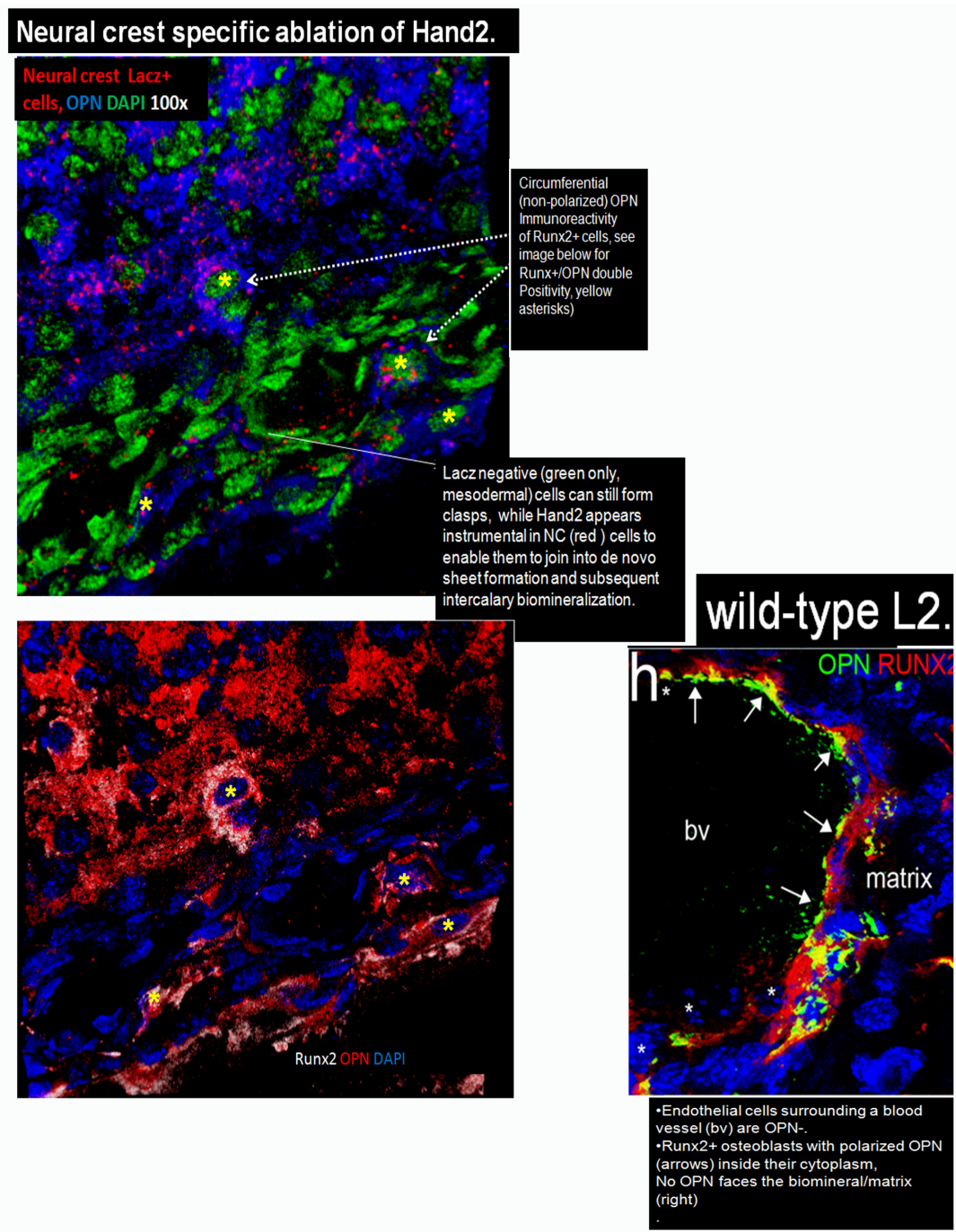

**Fig ESM06 – Osteopontin polarity and its loss in Hand2-/- RUNX2+ osteoblasts**

##### 1.3. A common origin of endothelia and osteoblasts

Our 3D analysis reveals that the elaboration of layer 2 involves a Hand2 dependent de novo sheet formation process: cells percolating from newly discovered rosette architecture from layer 2 are polarising, flattening up into contiguous sheets within L2 that are elaborating further with time. In the present paper, we show, surprisingly, that neural crest cells not only give rise to Runx2 + osteoblasts but also to a significant fraction of cd31 VWF+ endothelial cells inside cranial dermal bones (**Fig.1a-i** in main paper). Most interestingly, we observe pairs of neural crest derived CD31/vWF+ endothelial cells and Runx2+ osteoblasts that might suggest a common lineage origin. Only part of the frontal, clavicle (and nasal) bone examined is mesodermal (see **Fig.2A main paper** for non-GFP+ osteoblasts (i.e. mesodermal) directly lining the dark matrix. Because of this mosaic nature of the dermal bones examined finding pairs of endothelial/osteoblastic cells can either be explained by 1. Secondary meeting of these distinct cell types during the invasion process (but this would need to be NC lineage specific..) or 2. A common lineage origin of the two – and we see these cells to remain juxtaposed during development: while the endothelial cells join forces to establish bone vasculature vasculogenetically, the osteoblasts establish biomineral in the way we outline (**Fig4** accompanying paper and **Fig3,4 main paper**). During these two processes, the juxtaposition of osteoblasts and endothelia ensures nourishment of the biomineralizing layer as soon as the L2 vessels become connected up to the dermis and meningeal vasculature (first discernable at E17 in the frontal bone). This juxtaposition of endothelial and osteoblastic cells is not discernable in Hand2 neural crest mutant cells of the frontal bone any longer, however VWF+ endothelial cells are present within the L1/2 hybrid layer (**FigESM07**). This suggests that Hand2's function does not affect (endothelial) cell fate choices but the ability of nascent L2 pairs of endothelial and osteoblastic cells to coordinate their action.

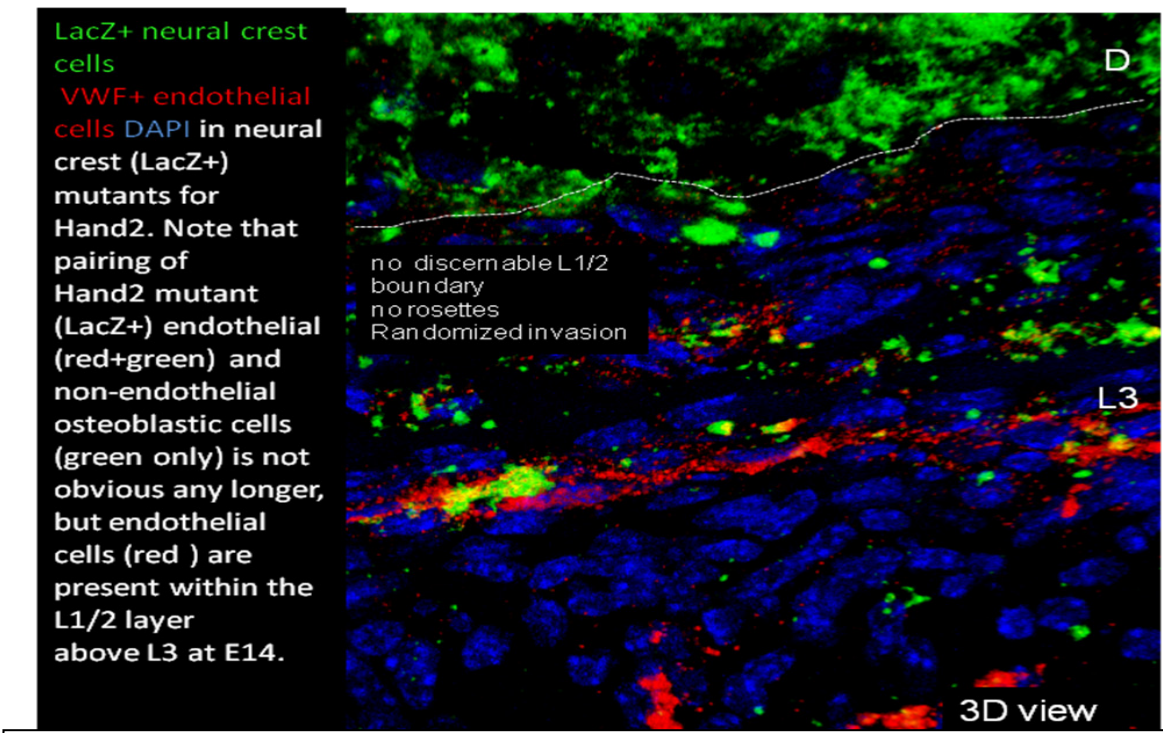

**Fig ESM07 – Loss of endothelial-osteoblastic pairing in Hand2 NC mutant cells**

Finding neural crest derived endothelial cells was very surprising, as all endothelial cells in the body has previously been considered to be solely of mesodermal origin.. A recent study had shown that neural crest cells in bone marrow in the adult are able to form endothelia inside the body postnatally (Nagoshi et al. 2008). We extend this finding into the earliest stages of dermal bone formation from E14.5 onwards. The notion that only mesodermal cells give rise to endothelia has emerged from chick quail grafting studies in the Le Douarin and other labs in the past (Couly et al. 1995, Etchevers et al. 2001), on the basis of the use of the so-called QH1 antibody which has been raised against an unknown endothelial epitope in quail. In the light of our findings, it is possible that this epitope might not be found on all endothelia such as the cranial ones that we have found to be CD31 +/VWF+. Moreover, antibodies against Runx2 were not available at the time to discover this ‘twinning’ of osteoblasts and endothelial stages – even if the earliest stages of dermal bone formation had been examined. Future work using the chick quail system and employing different antibodies will hopefully resolve why neural crest derived endothelia were so far not detected in the avian system.

###### 1.4.Relationship between this invasive cell type and biomineral features - discovering the anisotropy of two collagen networks (rather than one) within L2.

We discover a remarkable anisotropically distributed Collagen-MMP network generated by RUNX2+/OPN+ cells (**FigESM08**). Our single cell analysis reveals that these cells secrete collagen in a highly directional/polarized fashion: Collagen1 in the plane of cellular sheets forming de novo within L2 (**FigESM09**) and Collagen2 perpendicular to these sheets – into pre-existing matrix (**Fig2i-k**, main paper).

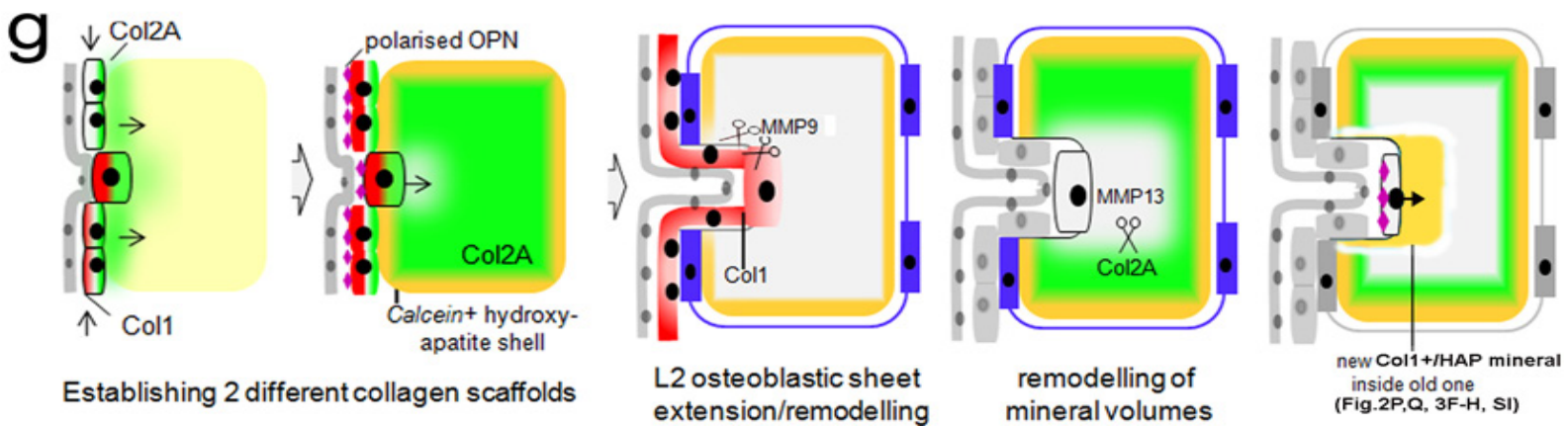

**Fig ESM08. Schematic of two anisotropic Collagen/MMP networks**

While collagen1 is confined to cellular sheets within L2, MMP9 which cleaves Collagen1 is also confined to these sheets and excluded from the matrix volumes (blue below ) (**FigESM09**).

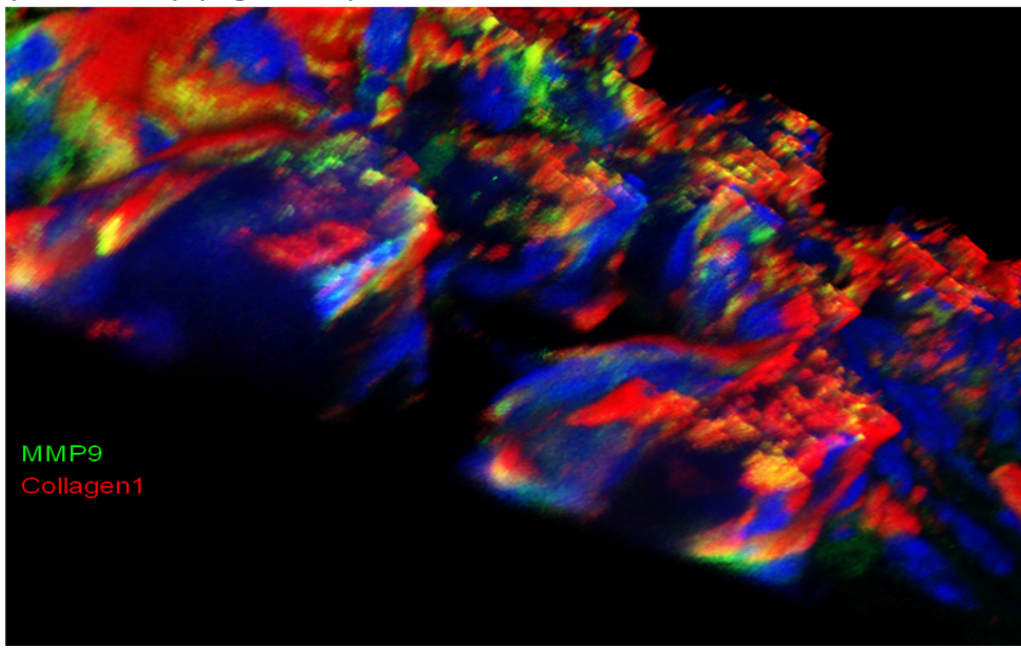

**Fig ESM09. 3D view of L2 matrix: Collagen1 and its respective MMP9 do not penetrate the matrix (blue) but stay confined to elaborating clasps.**

Collagen2 is also made by the Runx2+ invading cells (see **FigESM10**) as is MMP13 (**FigESM11**). Note that endothelial cells are not secreting collagen2 (front blue cells ensheathing the L2 blood vessel).

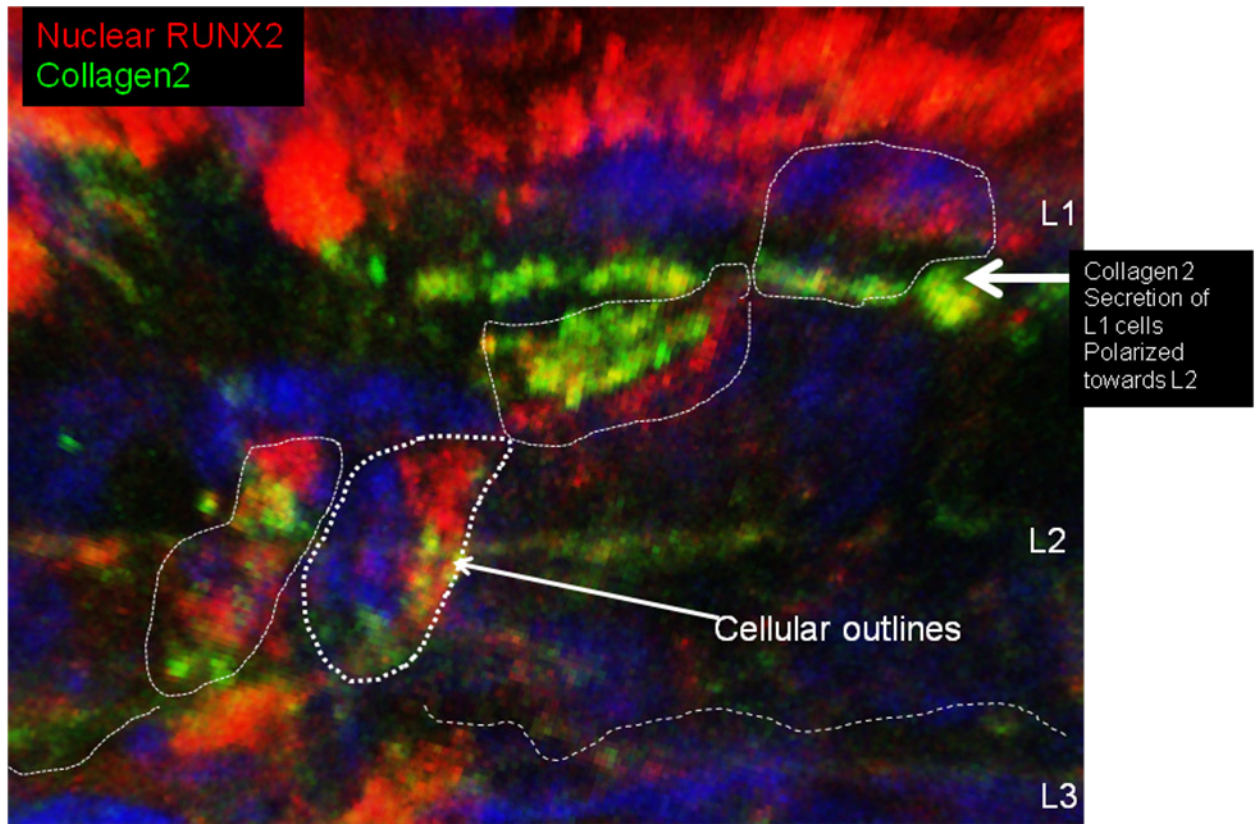

**Fig ESM10. Volume reconstruction of RUNX2+/Collagen2A positive cells**

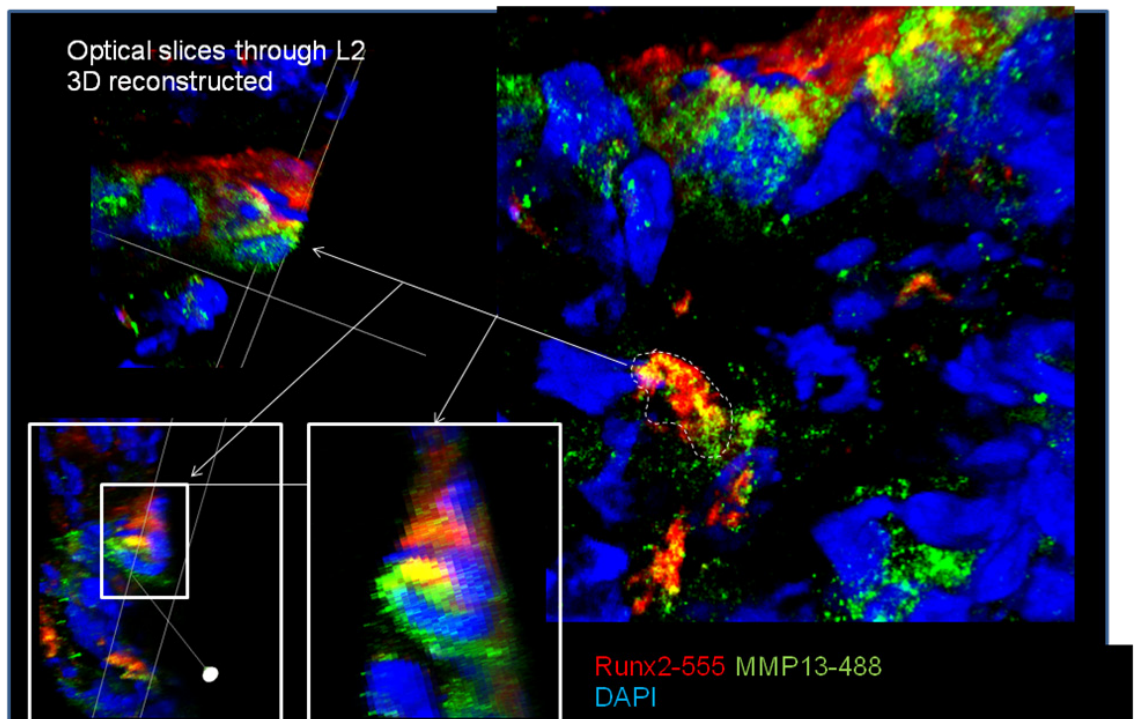

**Fig ESM11. These RUNX2 cells also secrete MMP13 which cleaves Col2.**

At later stages, the secreted MMP13 fills the L2 biomineral volumes in their entirety, rendering the colocalized Col2 more malleable:

Green – Calcein+ biomineral, red MMP13 (see **FigESM12** below), 3D view onto L2 upwards.

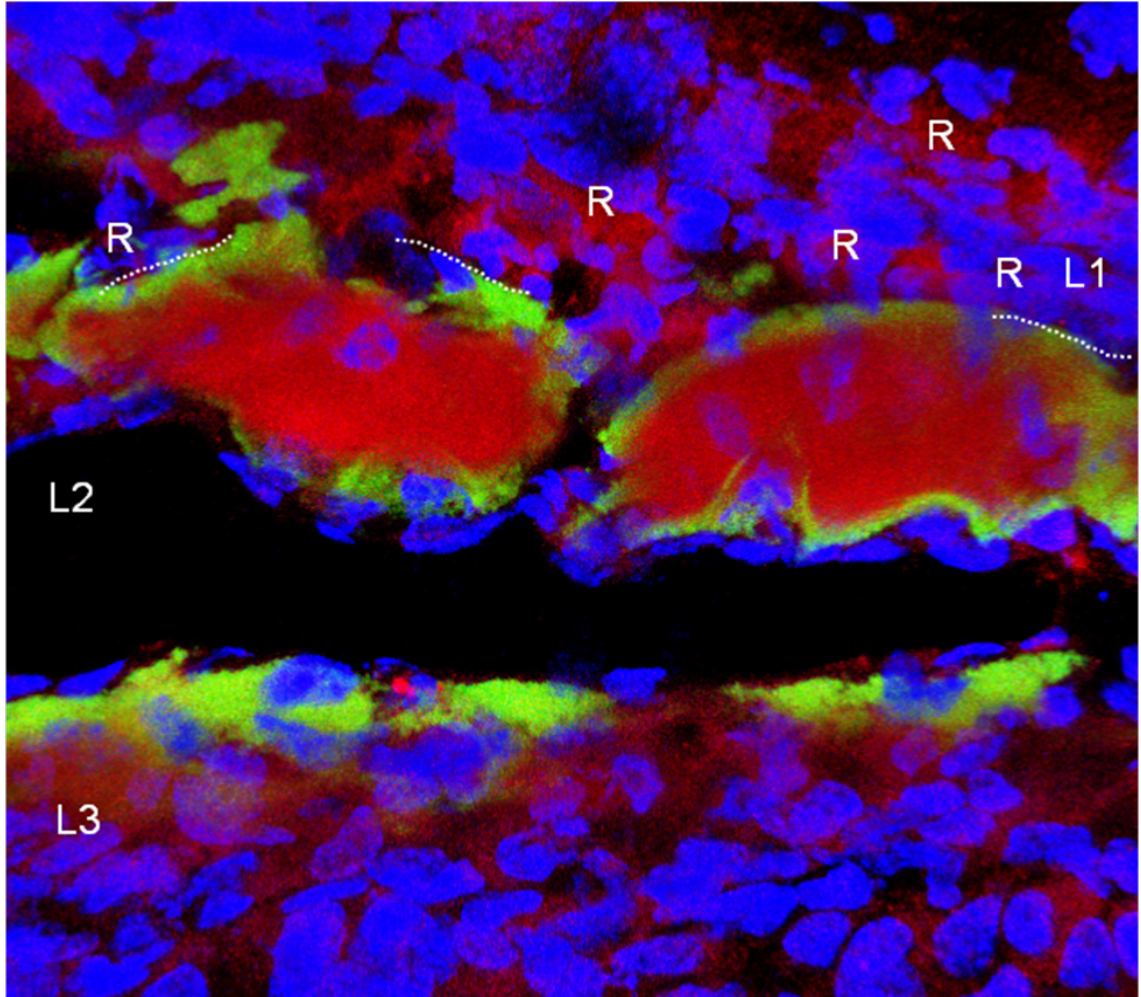

**Fig ESM12. MMP13 on the inside of Calcein+ matrix 'shells'.**

**FigESM13** strikingly shows that Collagen1 and 2 do not overlap when reconstructed in 3D (see Fig. 2b – and FigESM13 below, note no yellow areas of overlap !) – providing first ever evidence for a high degree of anisotropy of these two collagen networks. While the collagen1/MMP9 system is restricted to developing cellular sheets within L2, the collagen2/MMP13 is restricted to the centres of biomineral volumes. Recent *in vitro* studies had shown a role in fibroblasts of MMP13 in reshaping 3D collagen gel volumes (Toriseva et al. 2007), but its *in vivo* role in cranial dermal bone formation has not been assessed through colocalization studies at single cell resolution. The two systems communicate at those places where invasive OPN+/RUNX2+ cells interrupt the undulin 'wraps' surrounding the biomineral. This would not have been visible without 3D reconstruction at single cell resolution.

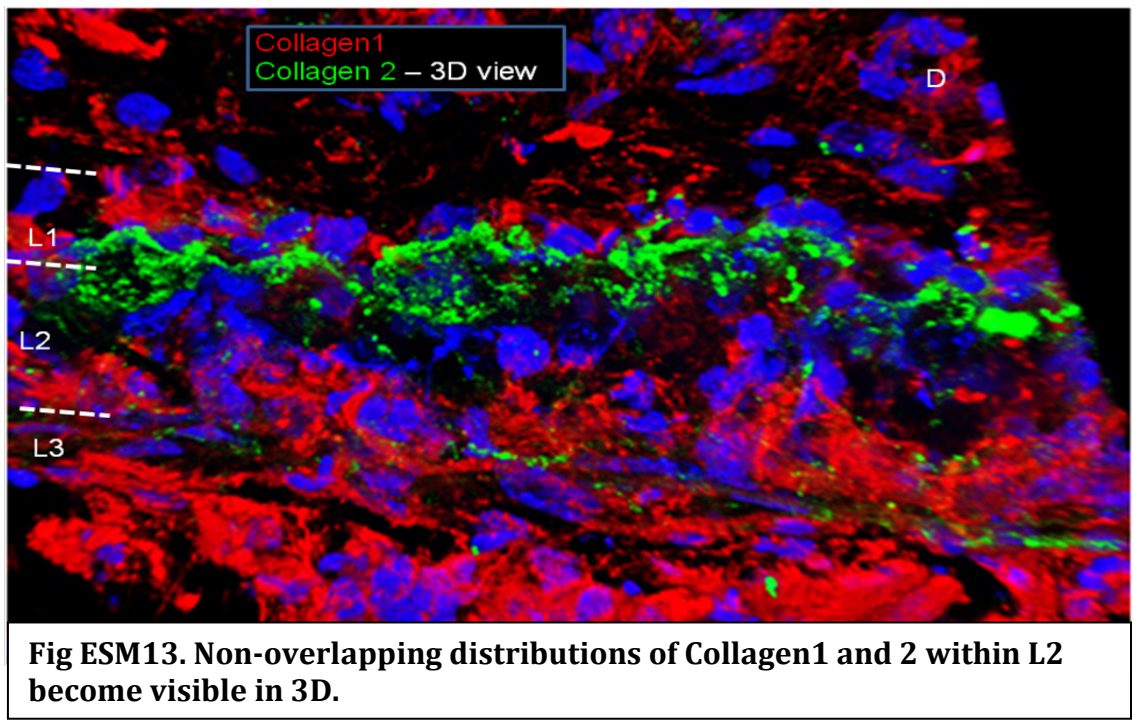

##### 1.5 Intercalary matrix secretion

We provide *in vivo* fluorescein pulse labelling experiments as direct evidence for new biomineral being deposited in an intercalary fashion (Fig.4 accompanying paper, Fig.3 present paper). To provide additional direct evidence for this unexpected finding we posited that vesicular membranes of GFP+ neural crest cells inside the matrix should be visible in a transgenic context. Indeed, **FigESM14** below shows GFP+ matrix vesicles secreted by neural crest derived osteoblastic cells within L2, forming halos of label around vGFP+ neural crest cells. This visualization was possible as we used a membrane-tethered vGFP as a novel (wnt-1-Cre) recombinase reporter mouse (to be described elsewhere). Current models of biomineralization assume secretion of collagen and additional factors as part of an exocytotic process involving, in this transgenic background vesicles with vGFP+ plasma membrane components (white arrows).

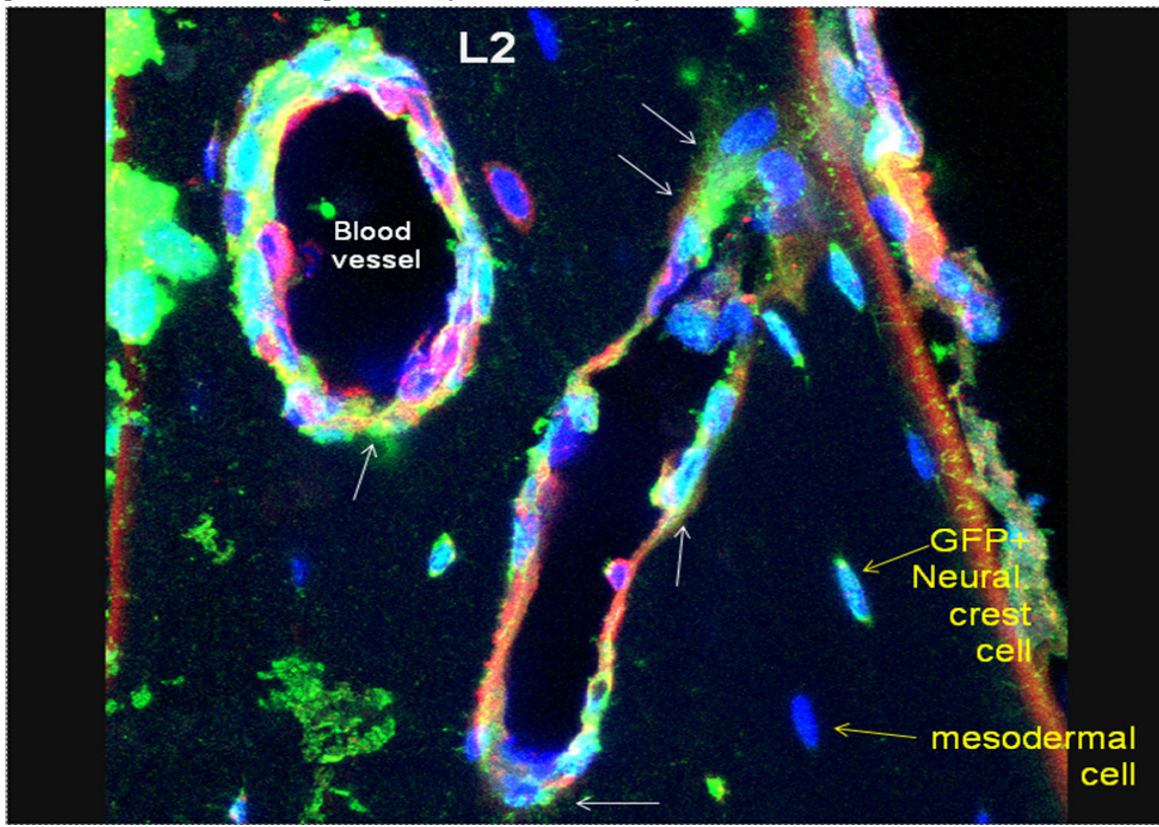

**Fig ESM14. Intercalary secretion/deposition of vGFP+ matrix vesicles within L2 of the more mature frontal at E18.**

#### 1.6. Hydroxyapatite crystals follow Collagen patterns

As hydroxyapatite crystallizes on collagen1 scaffolds, the biomineral formed provides direct evidence for the directionality of the underlying collagen networks and the cells making them, both in extant species such as mouse as well as in fossils. The figure below shows the well-established spatial relationship between collagen fibre bundles and the longitudinal (D) axis of HAP crystal orientation (**FigESM15**).

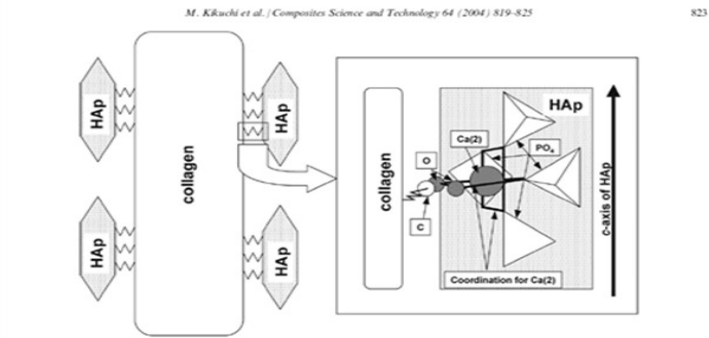

Fig. 5. Schematic drawing of the relation between self-organization (directional deposition of HAp on collagen) and interfacial interaction in the composite. Direction of interaction between HAp and collagen is restricted by covalent bond between COO and Ca(2) to maintain regular coordination number of 7.

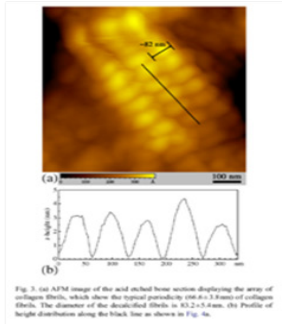

Fig. 3. (a) AFM image of the acid etched bone section displaying the array of collagen fibrils, which show the typical periodicity $63.2 \pm 5.3$ nm of collagen fibrils. The diameter of the fibrils is $167.2 \pm 5.3$ nm. (b) Profile of height distribution along the black line as shown in Fig. 3a.

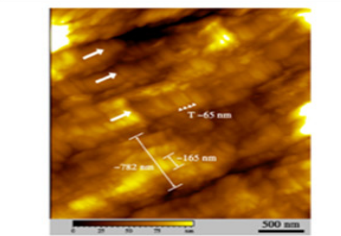

Fig. 4. Typical AFM image obtained from the unstained bone sections. The preferential orientations of the fibrils are demonstrated by the white arrows. The periodical cross-orientation pattern along the fibrils are pointed out by the arrow heads. The periodicity is $63.2 \pm 5.3$ nm. As indicated, several mineralized fibrils, with $167.2 \pm 6.8$ nm in diameter, aggregate into thicker fibers, with $743.8 \pm 68.7$ nm in diameter.

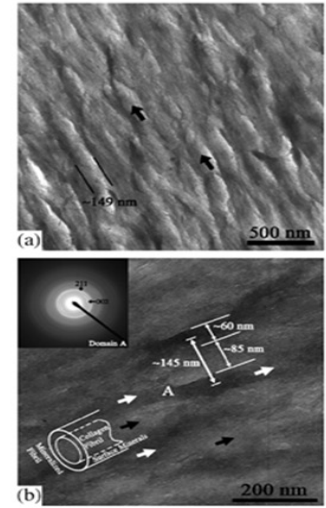

Fig. 2. TEM images of the unstained bone sections. The general image in (a) indicates the mineralized collagen fibrils, which are closely arrayed preferentially along the circumferential orientation of the bone wall as pointed by the black arrows. The diameter is $151.2 \pm 13.5$ nm. (b) presents the detailed structure of the fibrils at high resolution. The black arrows demonstrate the collagen fibrils with $85.8 \pm 5.3$ nm in diameter, while the white ones indicate the surface minerals that deposit along the fibrils with $60.6 \pm 5.6$ nm in thickness. The measurements to the characteristic sizes of the fibrils and the minerals illustrated that sum of the diameter of the fibrils and the thickness of the surface mineral layer matches the diameter of the mineralized fibrils. The SAD pattern (insert) in domain A displays that the crystals are HA in nature and with a preferred (002) crystallographic orientation over small regions examined.

J. Ge et al. / Materials Science and Engineering
C 27 (2007) 46–50

**Fig.ESM15, taken from Kikuchi et al. 2004 (left) and Ge et al. 2007 (right).**
**Registration of Col1 fibres and HAP crystals as seen by AFM in native tissue**

Furthermore reshaping of the entire L2 biomineral system – which we show in a developmental time course in the accompanying paper (**Fig. 3g,4**) and the present one (**Fig. 3K**) can now be seen to be accomplished by reshaping the collagen scaffold system, using the anisotropic collagen/MMP secretion mechanism we discover and describe in Fig.2.

##### 1.7. Other novel resorptive molecular features of Runx2+ osteoblasts within L2.

Moreover, collagen remodelling is one of the key aspects of biomineral remodelling, already deposited (Calcein+)HAP crystals need to be resorbed as well. To test this novel notion further we examined the Runx2+ osteoblastic cells for key molecules involved in the processes of biomineral erosion. Indeed, we discover inside the Runx2+/OPN+ neural crest osteoblasts TRAF6 and RANK (**FigESM16 and 17**).

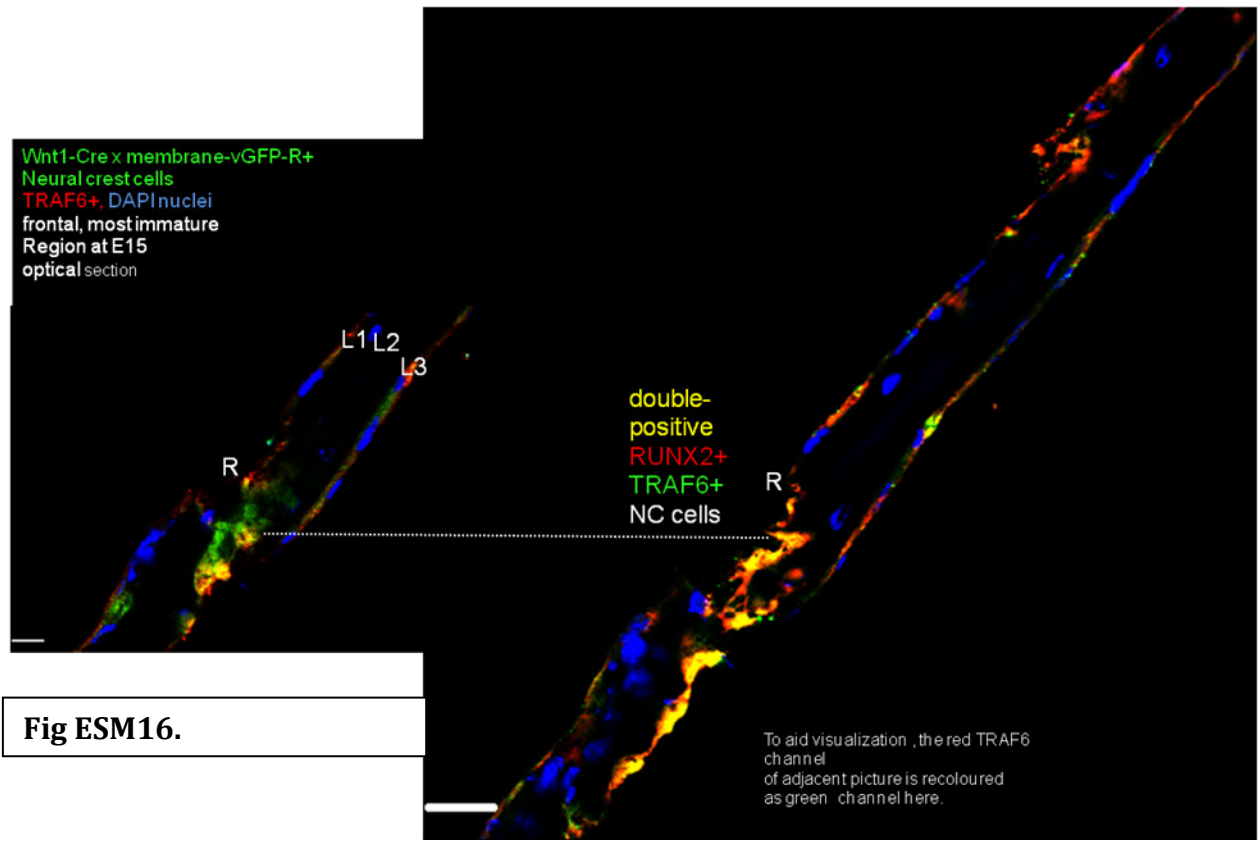

**Fig ESM16.**

Both RANK and its downstream effector/adaptor TRAF6 are critical for the mechanism of establishing podosomes and resorption lacunae (Kobayashi et al, 2001; Kim et al. 2005). Previously RANK and TRAF6 were only considered to be found on/inside osteoclasts (Asagiri and Takayanagi, 2007; Rubin et al 2005, Aeschlimann&Evans 2004, Darnay et al 2007, Dougall et al 1999). We now find these molecules already present at E15 – the earliest stages of dermal bone formation, (and L2 maturation) before the osteoclastic/monocytic lineage that relies on the formation of bone marrow even exists.

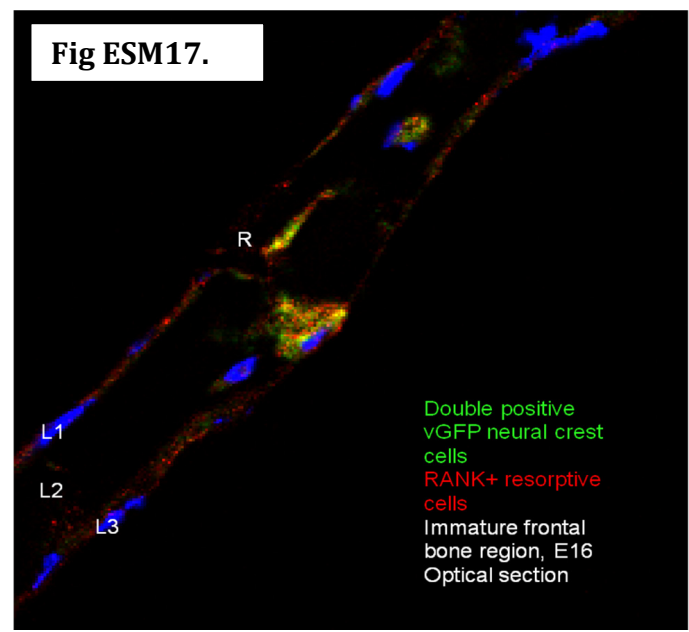

The optical sections show that these molecules are found *inside* (wnt1-Cre X membrane-vGFP reporter + green ) neural crest cells (that do not give rise to osteoclasts) and that are RUNX2+ osteoblasts (red in **FigESM17**), enabling these (now yellow, double positive) cells to establish podosomes and other resorptive cytological features.

#### 2.Ultrastructural predictions of collagen anisotropies – usable in extant and fossil taxa

We can identify 7 histological/microstructural hallmarks of the intercalary invasive process responsible for L2 and L3 expansion (see **FigESM18**, below) on the basis of the expression and properties of collagen fibres:

expression
biophysical
of collagen

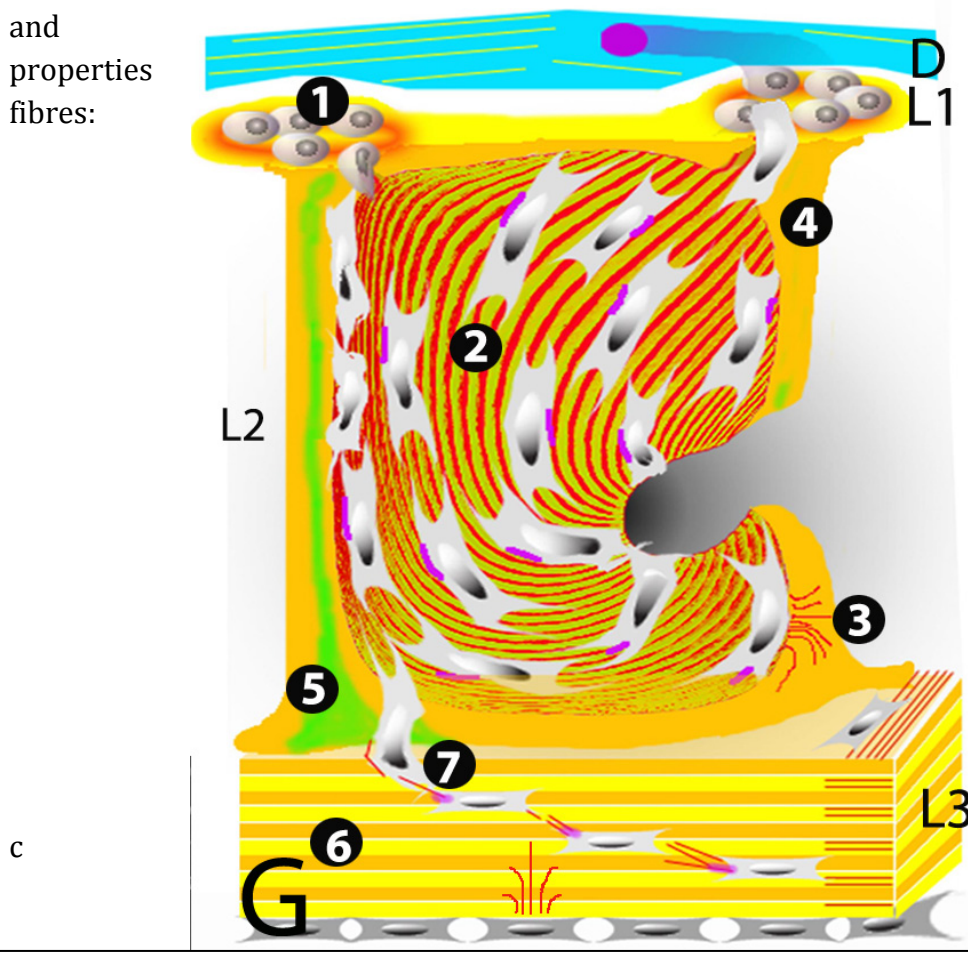

**Fig ESM18.** Schematic representation of early mouse, heterostracan and placoderm dermal bone generative anatomy, see below for details. Endothelial layer is omitted for reasons of clarity. D-layer in turquoise, osteoblast fibroblasts in pink, collagen yellow/orange, Collagen Fibre direction in red, Collagen2 green.

- 1.D-Layer-L1 discontinuity of matrix
- 2.Collagen fibre strap formation within L2.
- 3.Radial fibre orientations underneath migrating cells - orthogonal amplification of collagen mesh distortion.
- 4.Early Collagen1 composition of cancellar walls
- 5.Later Collagen1/collagen2 biphasic (sheetlike-amorphous) architecture of cancellar walls
- 6.Plywood self-organization of Collagen1 in layer 3 and 2
- 7.Later cellular invasion of L3 and intercalary matrix growth of the compacta.

Below we will provide detailed explanations of each feature.

#### 2.1. Biomineral features

1. **D layer-L1 discontinuity of matrix.** Collagen1 secretion by L1 cells is polarized towards L2 while collagen secretion within the D (dermis) layer is not. This would mean that cells in the dermis are surrounded from all sides by (biomineralized) collagen matrix, while there is a discontinuity of matrix just above L1. L1 cells are highly polarized in terms of cytoplasmic PERIOSTIN, HAND2, RUNX2 and cellular and extracellular Collagen1 deposition towards L2. Underneath the rosettes we find circularly organized collagen1 fibres surrounding a central hole adjacent to the invading nuclear RUNX2+/OPN+ cell (**Fig.3D,E**, arrow and arrow heads, main text). This provides an identifier for rosette architectures even in fossil material. Most importantly, this and the accompanying paper describe for the first time molecular marker distributions at the protein level that are not shared between cells in the D layer and Layer 1: Our single cell co-expression analysis find Collagen2 only to be made and polarized by L1 (and L2) cells towards the L2 biomineral, but not inside D layer cells, providing another molecular separation between D layer and L1 cells (**FigESM10**, white arrow.).

These collagen distributions provide the first evidence for a cellular and molecular separability of D and L1 that enabled us to identify L1 as layer in between D and L2 (or L3) deep into gnathostome ancestry.. We find this both in heterostracan and placoderm samples (see below and **Fig.4, feature 1**).

##### Layer 2:

2. **Collagen fibre strap formation within L2.** Extensive literature has shown

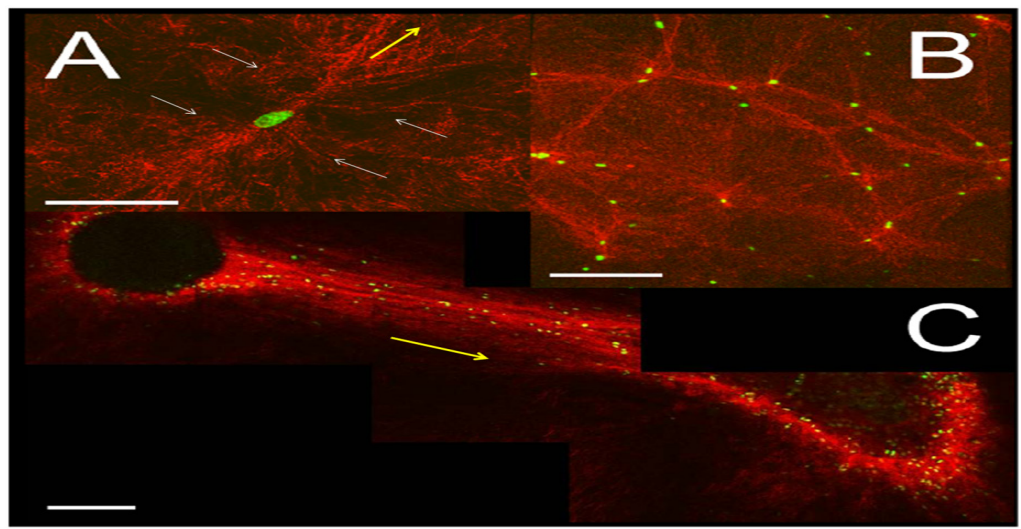

**Figure 1. Collagen gel morphological changes induced by presence of cells.** (A) Single U87 glioblastoma cell in a collagen network 10 hours after gel polymerization. bar = 50 μm. (B) Several U87 cells on the surface of a collagen gel 10 hours after gel polymerization. bar = 200 μm. (C) Two cell colonies embedded in a collagen matrix 48 hours after gel polymerization. bar = 200 μm. Fibers (artificial red color) are imaged through confocal reflectance; cell nuclei (green) are labeled with a GFP-histone heterodimer. doi:10.1371/journal.pone.0005902.g001

that cells migrating on top of (or through) an amorphous collagen1 matrix (red in **FigESM19** above) are able to organize this matrix along the long-axis of their filopodia/migration, leading to the formation of straps (yellow arrow in **FigESM19,20**) ( Vader et al. 2009, Petroll et al 2004).

Thus if straps can be seen in extant or fossil data this is a direct indicator of an underlying migratory process. We find this in both heterostracan and placoderm samples (**Fig.4, feature 2**).

3. **Radial fibre orientations underneath migrating cells - orthogonal amplification of collagen mesh distortion.** As a consequence of the non-linear properties of collagen networks cell migrating over a Collagen1 surface 'pull' collagen fibres into their path in a way that is orthogonal (white arrow, **FigESM20,21**) to their axis of migration (yellow arrow).

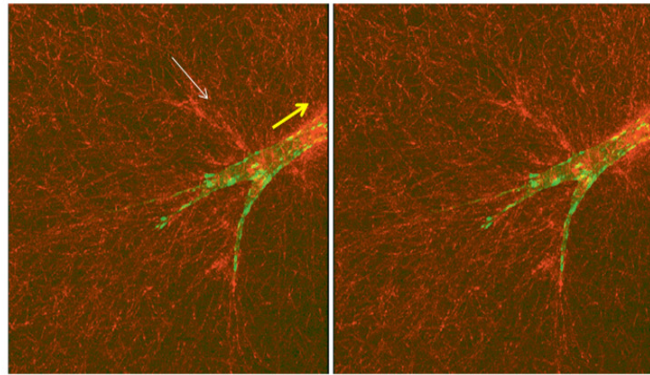

Fig. 3 Stereo pair reconstruction of a corneal fibroblast (TRK-36-zyxin) inside a three-dimensional collagen matrix generated using maximum intensity projections at $-5^\circ$ (a) and $+5^\circ$ (b). Reflected light (red) and fluorescent (green) images were obtained using visible light confocal microscopy. Horizontal field width $\approx 120 \mu\text{m}$.

**Fig ESM20, orthogonal collagen fibre traction, taken from Petroll et al, 2004.**

This non-linear phenomenon, termed orthogonal amplification of mesh distortion, has been well described in the literature studying cell-matrix interactions (Sawhney & Howard 2002 and literature therein). Once we recognize (as our experiments show) who produces Collagen1 at high concentration and the direction of invasion (**Fig.2** main paper), this biophysical phenomenon can explain radial/orthogonal tracks of collagen fibres towards the surface in aspidine as well as in cellular bones of osteostracans, placoderms or extant species. These have been called 'Sharpey fibre systems' in older literature, oriented perpendicular to

**Fig ESM21. Orthogonal matrix/mesh distortion. Taken from Sawhney & Howard 2003).**

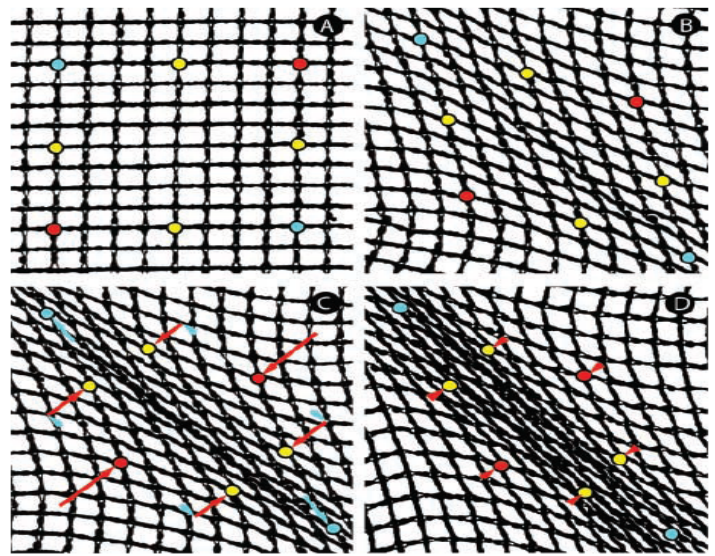

Figure 7. **Physical model of matrix movements.** A flexible yet inextensible mesh was imaged on a flatbed scanner. (A) Passive mesh, with reference points marked. (B) As tension was added from single points at the upper left and lower right corners, the mesh pattern became distorted, with square shapes elongating into diamonds as fibers aligned. (C) Further tension produced an aligned strap. Arrows show the mesh movements relative to the strap axis (parallel, blue; perpendicular, red). The tensile load resulted in a relatively small tensile strain, but a larger compressive displacement at a right angle. We term this feature of mesh geometry the orthogonal amplification of mesh distortion. (D) Pulling from multiple points adjacent to the axial corners increased the lateral draw of mesh and produced a broader strap.

bony layers and were previously only considered as part of muscle attachment systems. This is likely to be a misperception, based on the perceived static nature of biomineral, a concept rendered obsolete by our findings. Cells walking across collagen gels already generate this phenotype, a finding that supports reconstructing cells on the outside of a collagen layer on the basis of such orthogonal fibre tracks, even if no lumina (for cell bodies) can be discerned inside collagen matrix.

4. **Early Collagen1 composition of cancellar walls.** We find anisotropic secretion of collagen1 fibres in the plane of the newly forming cellular sheets/clasps within L2. Recent work on collagen fibre secretion mechanisms by the Kadler lab have revealed fibripositors (**pink in FigESM18**) as cellular structures of preferential collagen processing and deposition (Kadler et al 2008; Kapacee et al. 2008). Fibripositors are found at the lagging edge of cells - in contrast to previous concepts of secretion along the flat side of the cells. The existence of these cytological structures and the directionality of collagen1 secretion adds to the evidence **against** an appositional collagen1 secretion process (which by its very name implies **radial** collagen 'piling up' with oldest at the bottom and later collagen further up). Cells migrate and leave collagen behind at their lagging edge. Obviously when new cells arrive they will start to 'walk' across these collagen1 floors, generating the post-hoc impression of appositional piling up while actually the depositional process follows a horizontal vector. As collagen1 does not percolate into early L2 matrix but stays confined to L2 clasps/sheets its concentration is higher along cellular sheets.
5. **Later Collagen1/collagen2 biphasic (sheetlike-amorphous) architecture of cancellar walls.** We observe Runx2+/OPN+ cells also to be positive for Collagen2 and see that collagen2 positivity increases with time (and elaboration of L2) (**Fig.2i-k**, main paper). Most interestingly, Collagen2 is secreted perpendicular to the L2 sheets deep into pre-existing matrix of L2. This and its cleavage by MMP13 which is injected into the biomineral volumes by Runx2+ cells leads to amorphous collagen matrix at the centre of more mature (older) cancellar/biomineral walls, encased by sheetlike Collagen1 on the surface of cancellar walls. We see this morphology both in mice, heterostracans and placoderms.

##### **Layer 3:**

6. **Plywood self-organization of Collagen1 in layer 3 and 2.** As discovered by the laboratory of Giraud-Guille, Collagen1 has the remarkable biophysical property to self-organize into liquid crystals of a cholesteric geometry without the presence of cells or accessory presence (**FigESM22 and 23**, below), leading to plywood microstructures (Giraud-Guille et al. 2003, 2005, 2008). This property is dependent upon very high concentration of

Collagen1 and provides a minimal/parsimonious explanation for both extant and fossil plywood architectures found in L2 and – dominantly in L3.

A remarkable transition in the self-organizational properties of Collagen1 that solely depends on concentration has been found (**FigESM22**).

*M.M. Giraud-Guille et al. / Current Opinion in Colloid & Interface Science 13 (2008) 303–313*

311

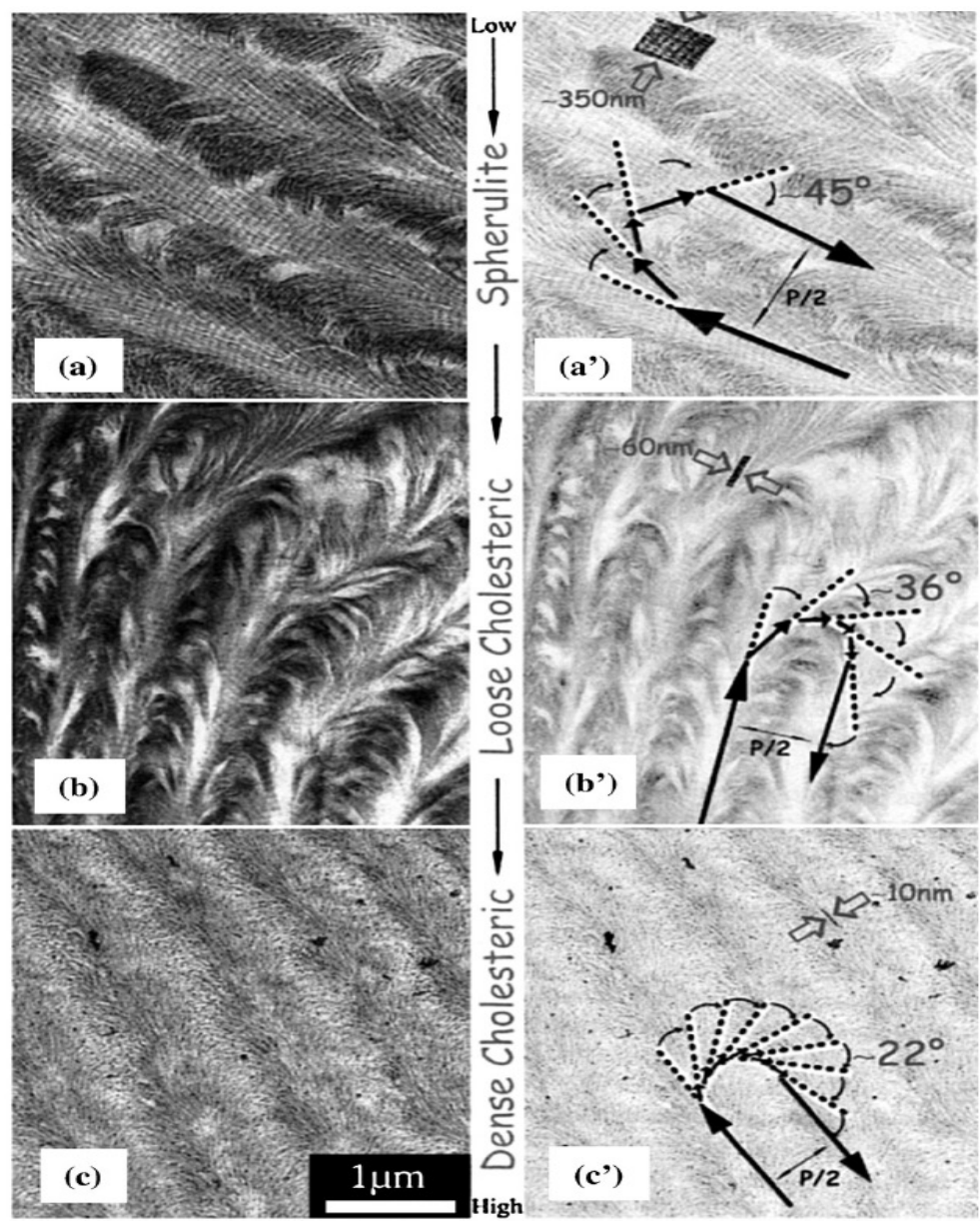

**Fig. 7.** Collagen fibrillar arrangements in cell free assembled matrices. (a,b,c) Ultrathin sections showing dense tissue-like collagen packing at different concentration dependant levels in a micro-chamber device. (a') Discontinuous rotation angles and large fibrils at a lower concentration. (c',d') Regular series of arced patterns showing smaller fibril diameters and rotation angles as the concentration increases; from reference [8\*] permission requested. a,b,c: bar= 1 μm.

**Fig ESM22 – Concentration-dependent self-organization of Collagen1 in vitro into plywood architecture, taken from Giraud - Guille et al. 2008.**

We show that in wildtype the Runx2+/OPN+ cells within L2 express and secrete Collagen1 at high concentrations, as do (OPN+) L3 cells. L3 in the mouse is characterized by plywood architecture (**Fig.3i** PMT image), a structure previously termed isopedine in fossil early gnathostomes. Please see below (**FigESM24**) an image of a heterostracan Pteraspis L3, which displays the same cholesteric collagen1 architecture seen in **Figs ESM22,23b,c**).

**Fig.ESM23. Self-organization into cholesteric Collagen1 crystals from above. Taken from Giraud-Guille et al. 2005.**

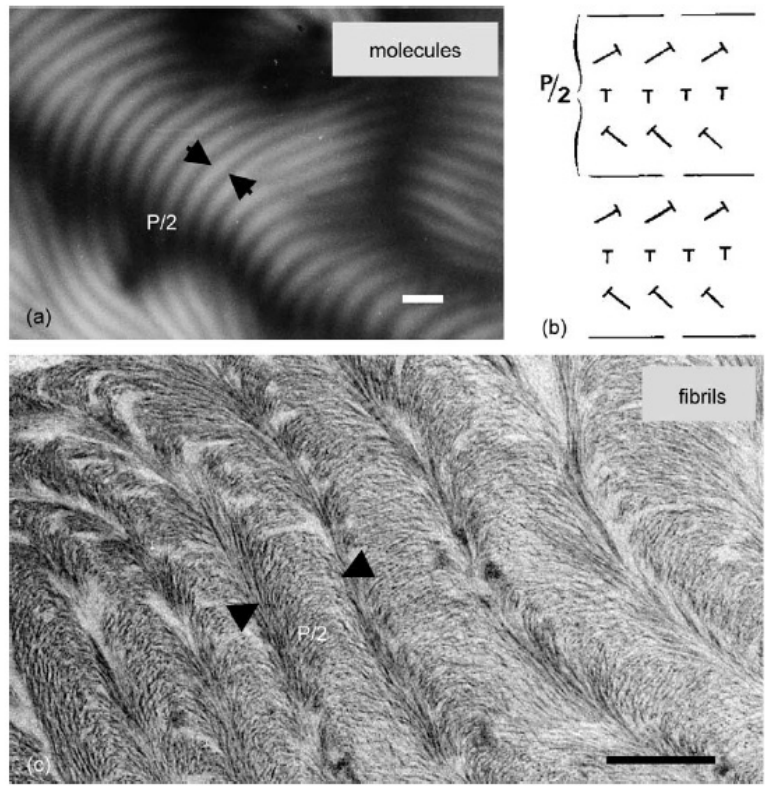

Fig. 2. Self assembled collagen at the molecular and fibrillar level: (a) acid soluble collagen solution at high concentration assembled in a liquid crystal of the cholesteric type, characterised by typical fingerprint patterns in polarised light microscopy, bar=5 $\mu\text{m}$; (b) diagram of a cholesteric geometry where nails represent molecules oblique to the observation plan, $P/2$ corresponds to a $180^\circ$ rotation of the molecular direction; (c) stabilised liquid crystal at neutral pH inducing collagen fibrillogenesis. The overwhole cholesteric geometry, evidenced by regular series of arced patterns, is maintained. TEM, bar=0.5 $\mu\text{m}$.

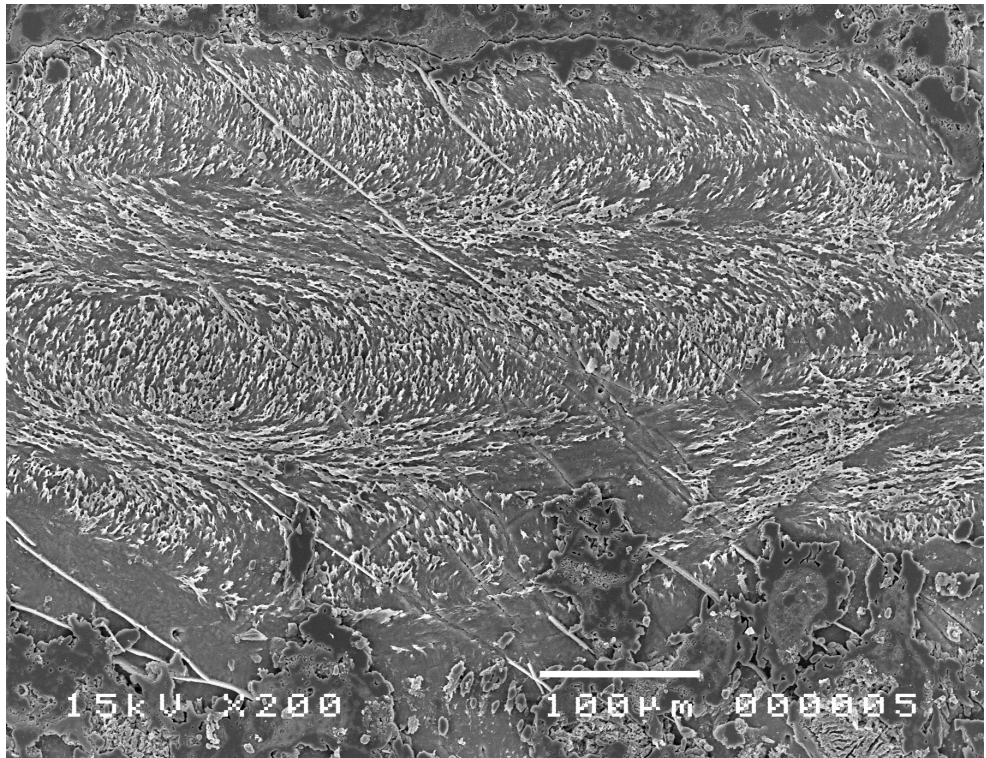

**Fig ESM24. Collagen1 plywood liquid crystals in Pteraspis L3 isopedine (Donoghue, unpubl.)**

Thus high expression/concentration of Col1 is sufficient to generate plywood patterns. This informs paleontological interpretations, as one cannot a priori posit that a collagen sheet closer to the margins is 'younger' than a more internal one – which an appositional model might posit. As orthogonal matrix distortion would take place when L3 cells are sitting underneath the L3 plywood architecture, one would expect to find radial matrix tracks in this region. We find these both in various heterostracans, placoderms and mice.

A significant collagen1 expression and plywood layer 3 is even present in Hand2 mutant animals in which L2 does not get elaborated (Fig.ESM25, below, see accompanying paper, Fig.3).

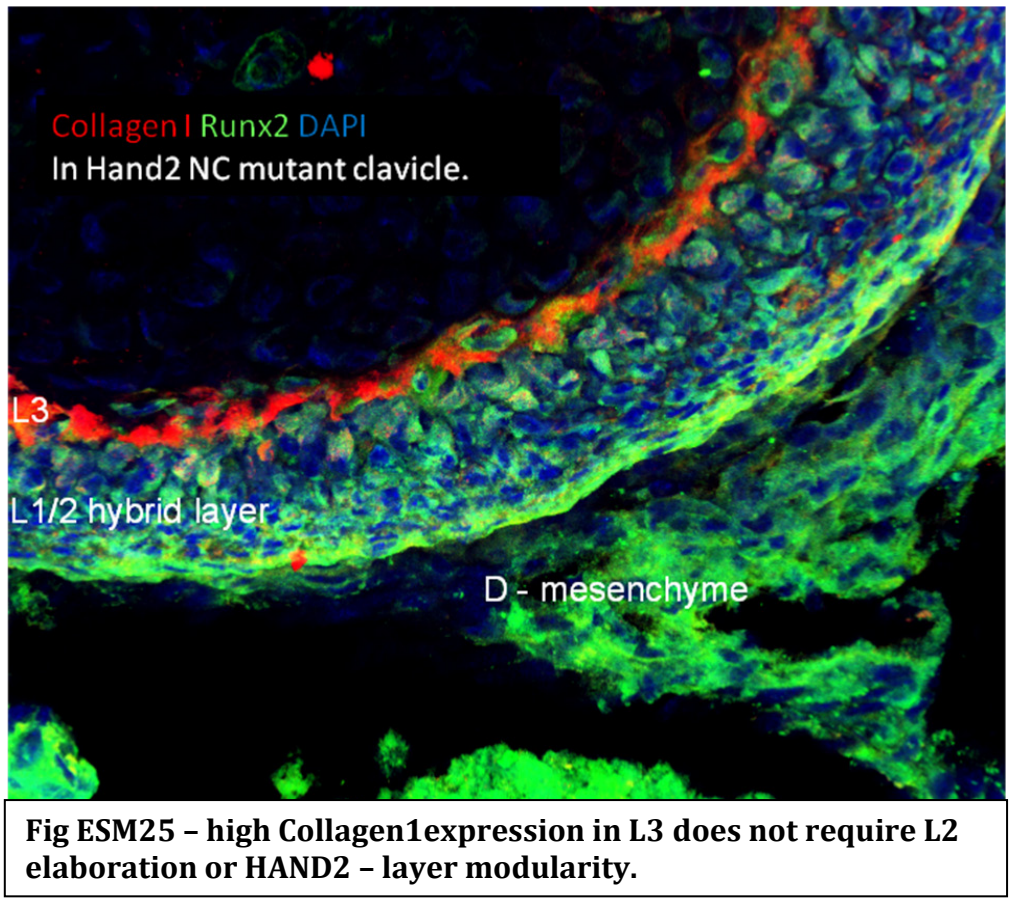

This extant example provides evidence for the functional modularity of the layer organization – each can be separately elaborated in ontogeny or in phylogenetic time (or both). Our experiments suggest that subtle changes in RUNX2 expression and protein localization as the molecular target of these fundamental historical differences.

###### 7. Later cellular invasion of L3 and intercalary matrix growth of the compacta.

Indeed, at later stages of mouse L3 growth we see that the thick collagen1 plywood becomes intersected by radial tracks of cells migrating in, both in a radial and horizontal orientation (**Fig.3i,j**). Our in vivo biomineral labelling experiments show that later deposited matrix (unlabelled) intercalates in between earlier (red) labelled matrix: yet another example of an intercalary matrix growth that only experimentation can reveal (**Fig.3b,c,f,g,h**). In early mouse development very few cells are positioned within the collagen1 plywood architecture, in later development that number increases, followed by invasion of vasculature into this plywood matrix (**Fig.3i**).

This phenotype is replicated during placoderm ontogeny of *Bothriolepis*, compare L3 in early larvae vs mature dermal bones (**FigESM18**, feature 7

and in **Fig.4C vs 4S**). As we do not find cellular lacunae within L3 plywood of heterostracans (**Fig.4h,i,q,r**), but we do find them in osteostracans (see **FigESM33 and 34**, below), we infer that such invasive feature of L3 growth is related to the evolutionary emergence of cellular bone.

Remarkably, corneal collagen1 matrix displays a similar plywood architecture as L3 and it was recently reported that Osteopontin is required for cellular invasion into such plywood structure (Miyazaki et al,2008). We predict a shared role of OPN in osteoblasts that invade into the L3 plywood matrix which future experiments could reveal. Currently, the L3 cellular population is not genetically trackable.

##### **Deciphering the temporal axis of matrix deposition and remodelling**

Plywood architecture of Collagen1 can not a priori be interpreted as evidence for appositional growth due to the capability of collagen1 to self-organize (**FigESM22,23**). However, our in vivo biomineral labelling shows that if labelled (earlier deposited) matrix is intersected by unlabelled matrix in a radial fashion (**Fig.3i**), this must have been caused by a later invasion of cells. Thus, if we find in fossils horizontal plywood architectures intersected by cellular lacunae in a radial direction – or even by vasculature (**FigESM33,34**) – this process would have taken place after the deposition of the plywood Collagen1 matrix. Radial fibre direction alone (without cellular lacunae) can only be used as indicators of orthogonal collagen mesh distortion and as proxy for cells positioned at and migrating across the surface matrix layer.

The feature 1-7 outlined above provide criteria for layer identification in fossils and – as our ontogenetic data in mouse shows a development from a L1/L3 to a L1/L2/L3 configuration, this provides an ontogenetic axis by which we can interpret fossil ontogenetic material: We find the same transition from a bilayered to a trilayered bone in placoderms and heterostracans, bracketing all gnathostomes.

##### **3.Reconciling growth of the dermal skeleton in the earliest skeletonising vertebrates**

Despite considerable interest in the early evolution of the vertebrate dermal skeleton, attempts to resolve the pattern of development of the dermal skeleton in any given clade have proven problematic. Most effort has been expended in explaining the growth and patterning of the dermal odontodes, giving rise to the Ørvig and Stensiö's 'lepidomorial theory' and Ørvig and Reif's 'Odontode theory', but this attempts to explain only the most superficial aspect of the dermal skeleton. Circumference parallel lines have been observed in the superficial layer, suggestive of a pattern of marginal appositional growth, in placoderms, osteostracans and heterostracans (for summary see Janvier 1996). However, when the deeper layers within the dermal skeleton have been investigated, they have shown no evidence of marginal appositional growth. It has been an expectation that the original structure of the dermal tissues should be preserved and merely augmented through appositional growth, and that this should be preserved in the skeletal tissues. This perspective is supported by the view that dermal tissues show little evidence of remodelling in stem-gnathostomes. However, these facts cannot be reconciled with the fact that the skeletal elements occur in a (sometimes very great) range of sizes and thicknesses, reflecting ontogenetic growth (White and Toombs, 1983). The solution to this conundrum is that the dermal skeletal tissues were much more dynamic in vivo than has been perceived hitherto. The growth rings within the superficial odontode layer do indeed reflect a pattern of margin appositional growth, but the deeper layers within the dermal skeleton were in flux through ontogeny, resorbed and expanded through intercalary inflation, destroying the record of gradual marginal growth of dermal skeletal elements.

Below we review the dermal skeletal histology of the component clades of stem-gnathostomes, including any evidence there is for the ontogeny of this skeletal system.

###### **Heterostraci**

Heterostracans exhibit a variety of dermal skeletal architectures, from tessellate arrangements of scales or plates, to a small number of large fused or unfused plates, to groups in which the dermal skeleton is comprised of just one or two large ossifications (Halstead 1973). In any instance, the composition of the dermal skeleton has been interpreted to be the same, comprised of a superficial layer of scales, an intermediate cancellar layer overlying a basal lamellar layer (Janvier 1996). However, we recognise a distinct histological layer intermediate of the superficial layer and the cancellar layer that develops independently of the surrounding tissue layers. This division of the heterostracan dermal skeleton can be distinguished in all other heterostracans that we have investigated and, thus, we consider it to be general for the clade.

The superficial layer of dermal tubercles composed of dentine and enameloid comprise the D layer. The thin underlying layer, superficial to the cancellar layer, appears to lack the fibre matrix that is so conspicuous in the underlying tissue layers, and exhibits a regular series of holes, approximately 20 $\mu\text{m}$ in diameter, that connect the overlying dermal tubercles to the underlying cancellar spaces; we interpret this thin tissue layer as L1. The cancellae, which comprise L2, are defined by walls perpendicular to layers 1 (overlying) and 3 (underlying), composed of a rich complex fibre matrix. The honeycomb-like architecture of these cancellar spaces is distinct and independent from the planar arrangement of dermis vasculature. This suggests independent morphogenesis and secondary registration of the two systems. After initial construction, the cancellae in L2 are subdivided by cross walls that have the same structure and, eventually, the spaces are partially filled by centripetal appositional growth. The matrix of the initial cancellar walls is initially structured but this diminishes in the core of the walls during development. The basal lamellar layer represents L3 is comprised of a plywood-like arrangement of sheets of acellular but fibre-rich tissue, permeated by oblique to orthogonal fibre bundles that have traditionally been interpreted as aspidinocytes (Halstead Tarlo 1963) or Sharpey's fibres (Ørvig 1965). However, these structures are more readily reconciled as collagen I fibre bundles that arise through (a) self-organisation of Collagen I at high concentration (see **features 4, 5, 6 in FigESM18 and Fig.4G**, above), and (b). Orthogonal/radial distortions of the collagen matrix as a consequence of cell migration (**feature 3** above). The pattern of establishment and subdivision of the cancellar spaces is compatible with the process of mineralization of L2 in mouse.

Interpretations conflict on the sequence of development of these component tissue layers. All studies agree that the superficial layer of odontodes forms first, and that the deeper tissues form latterly, with the progressive development of the basal lamellar layer and the vertical walls of the overlying cancellar layer (Denison 1964, 1973; White 1973). Earlier, Fahlbusch (1957) had speculated that the innermost areas of a given spongy bone area are the most mature and that what we would now term L3 could give rise to material contributing to L2 and L3 plywood could grow appositionally from the bottom.

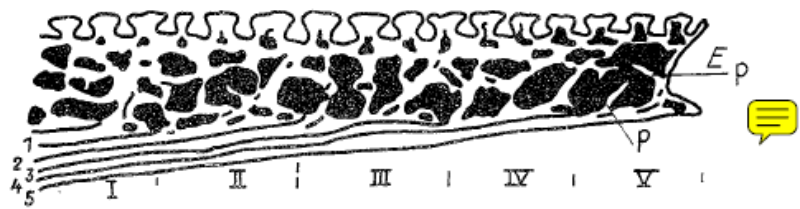

Abb. 4. Querschnitt durch den Plattenrand einer Dorsalplatte. Stark schematisiert. 1—5 die einzelnen Basalschichtlagen, 1 die älteste, 5 die jüngste. I—V die einzelnen Zuwachszyklen, I der älteste, V der jüngste. E = randlicher Gewebesaum. P = Primäranlage der Spongiosabälkchen.

**Fig ESM26. Pteraspis dorsal plate, inferred growth sequence, taken from Fahlbusch 1957.**

However without any experimental foundations established here, many different mechanisms could possibly lead to the same final pattern and the self-organizational properties of collagens were unknown at the time that can equally explain the growth of L3 plywood. Moreover, the difference (which we now understand as ontogenetic change) in the architecture of the L2 cancellar spaces could not be understood in any appositional sense. More recently Pernegre described (without providing histology) a ventral plate of the pteraspidifom heterostracan *Woodfjordaspis felixi* which displays the most immature area without L2 cancellae centrally, surrounded by cancellar (L2) areas (Fig.3 in Pernegre 2006).. Such architecture (with the more mature regions surrounding a less mature one) would render any appositional growth at the plate margin impossible, but no experimentally tested framework was available to explain the malleability of L2 biomineral at those early stages of cancellar formation.

Thus, no (fossil) data – without an experimentally proven process to compare against- could discriminate the relative timing of development or mineralization of the cancellar and basal lamellar layers. There is scant evidence to suggest that the trabeculae of the cancellar layer form first (Denison 1964, 1973), but also that the cancellar and basal lamellar layers form together (White 1973), and that the basal lamellar layer forms before the overlying cancellar layer (Greeniaus and Wilson 2003), but it is difficult to discriminate between preservational artefacts and true underlying processes. These differences might reflect differences in timing of mineralization, rather than development of the tissues. Nevertheless, it is clear that the spongy (now termed L2) and lamellar layers (now termed L1,L3) continue to develop through ontogeny, the lamellar layer growing through apposition at its base and the cancellar layer increasing in thickness. The pattern of inflation of L2 has not been documented although a later resorption in L2 has been documented (Donoghue and Sansom 2002).

The structure of the heterostracan dermal skeleton and the pattern of development that it exhibits is directly compatible with the processes seen in dermal bone development in mouse. Indeed, the structure of L2 – the classic ‘aspidin’ tissue of heterostracans, is also readily interpretable in light of our mouse model for dermal bone development. The nature of aspidin has been the subject of much debate, with interpretations polarised between interpretations as a fibre-rich acellular bone (Agassiz 1833-43; Pander 1856; Huxley 1858; Powrie 1870; Stensiö 1927; Gross 1930, 1935; Ørvig 1951; Bystrow 1955; Ørvig 1958b, 1958a; Gross 1961; Denison 1963; Ørvig 1965; Denison 1967; Ørvig 1967; Moss 1968a, 1968b; Ørvig 1968), to a cellular bone in which the spindle-shaped spaces reflect ‘aspidinocytes’ – fusiform osteocytes (Rohon 1893; Obruchev 1941; Halstead Tarlo 1963, 1964; Obruchev 1964; Halstead Tarlo 1965; Halstead 1969, 1973, 1974). Halstead (1969, 1973) identified three main types of structure within aspidin: coarse fibre bundles at the inner surface of the skeleton that he identified as Sharpey's fibres, spindle-shaped spaces that occupy the area between the vascular canals, interpreted as aspidinocyte cell spaces, and fine-calibre tubes that are radially arranged about the vascular canals, that he interpreted as spaces for cell-processes. The opposing view, that putative 'aspidinocyte' cell spaces represented the site of collagen fibres, or more specifically, Sharpey's fibres, was at the same time maintained by Gross (1935), Bystrow (1955), Ørvig (1958a, 1965, 1967, 1968), and Moss (1968b). These competing interpretations can now be reconciled readily since comparable structures develop within the collagen matrices of developing dermal bone as a consequence of osteoblast migration, distorting the collagen matrix and leaving behind tracks recording the path of migration. These are the radial canals that occur within the osteon-like tissue layers lining cancellae within L2. In a sense, these do indeed represent cell spaces, albeit osteoblast cell spaces, not 'aspidinocytes'.

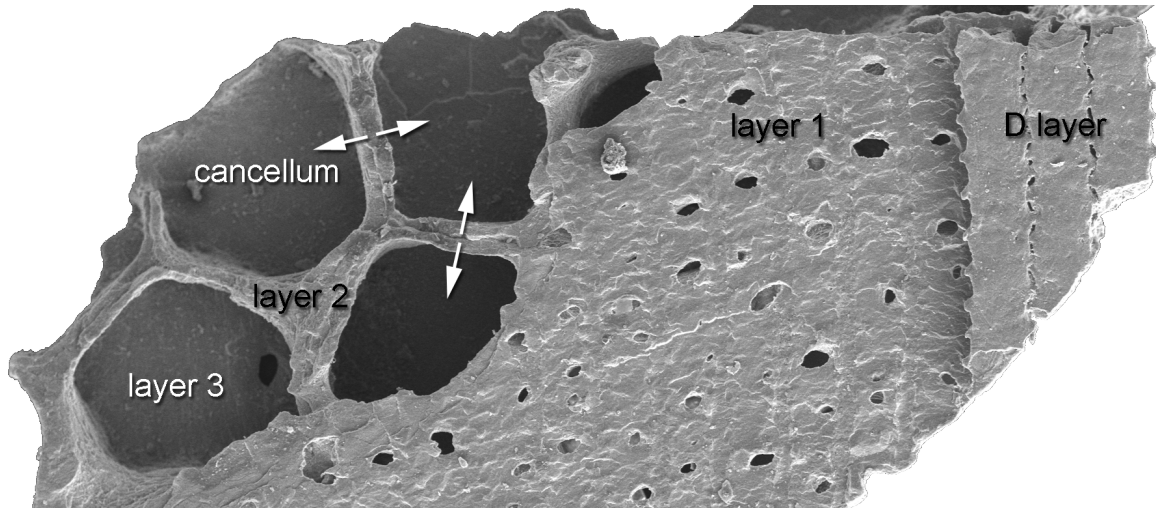

**Figure ESM27:** The four-layered structure of the pteraspiderostracan dermal skeleton, external view. Fragment of the dermal skeleton of a unnamed pterapid from the Lower Devonian of Prince of Wales Island, Canadian Arctic (NRM.PXXXX). The surface tubercles (D-layer) occur to the right, aligned vertically; they have been removed from the rest of the fragment, revealing a distinct thin structureless underlying layer (L1). Below L1, occurs the honeycomb-like cancellar layer (L2); note that the organization of the cancellae is distinct from that of the overlying tubercles of the D-layer. L2 is floored by the basal lamellar layer (L3) which exhibits a plywood-like structure. Note that the vertical walls of the cancellae are bilayered, reflecting the centripetal growth from the core of the wall; arrows indicate the direction of growth.

**Figure ESM28:** The four-layered structure of the pteraspid heterostracan dermal skeleton, viewed in cross-section. Fragment of the dermal skeleton of a unnamed pterapid from the Lower Devonian of Prince of Wales Island, Canadian Arctic (NRM.PXXXX). The D-layer is comprised of longitudinally arranged ridges of dermal tubercles and their associated vasculature. This is separated by a thin layer of structureless tissue (L1), below which occur the radial (with respect to the body axis) cancellae occur (L2); the plywood-like L3 occurs at the base. Note that the arrangement of the dermal tubercles and their associated vasculature (D-layer) are not coordinated with the L2 cancellae, indicating distinct patterning.

This arrangement is typical of pteraspid heterostracans (**Fig.ESM27,28**), but other clades within Heterostraci are variations on this theme. The dermal skeleton of cyathaspids such as *Anglaspis* (**FigESM29**) appear to have grown in a single episode but the overall structure remains the same.

**Figure ESM29:** The four-layered structure of the cyathaspid heterostracan dermal skeleton. Fragment of the dermal skeleton of *Anglaspis macculloughi*, a cyathaspid, from the Lower Devonian of Earnestry Brook, Shropshire (NHM.PXXXX). The D-layer is comprised of dermal tubercles, separated from the cancellae that comprise L2 by a thin structureless L1. The cancellae show evidence of centripetal appositional layers but the core of the walls is structureless. Some of the cancellae have been bisected by cross-walls. The basal layer exhibit a plywood-like structure characteristic of L3.

Tessellate heterostracans, the phylogenetic position of which within Heterostraci is unresolved (Janvier 1996), show the most significant differences in structure of the dermal skeleton. *Lepidaspis* has a structure that is simplified with respect to pteraspids (**FigESM30**). The overall architecture is the same, but L1, L2 and L3 are comparatively underdeveloped, equivalent to an early stage of dermoskeletal development in pteraspids. We identify a thin L1 (identified as an unmineralised matrix discontinuity at the base of the D-layer, see Point 1 above); the walls of the cancellae that comprise L2 show the typical structure of 'aspidin' with a meshwork of fibre-like spaces that show a radial organisation with respect to the cancellae. These are the same structures that we interpret as osteoblast migration tracks, above. However, the cancellae show evidence of subdivision through development of cross-walls, as seen in pteraspids (above) and mouse (see our other paper).

**Figure ESM30:** The four-layered structure of the dermal skeleton in the tessellate heterostracan *Lepidaspis*, from the Lower Devonian of Prince of Wales Island, Canadian Arctic (NRM.PXXXX). Upper figure: dermal tubercles (D-layer), separated by a discontinuity comprised of a structureless matrix (L1), overlying a cancellar layer (L2) with a rich fibre matrix comprising the primary walls and cross-walls, overlying a basal lamellar layer (L3). Lower left: dermal tubercle with dentine core and enameloid cap; lower middle: structureless L1 separating D-layer (above) from L2 (below); lower right: radial fibres in the primary walls of the cancellae, formed through distortion of the collagen matrix as a consequence of cell movement.

Other tesserates, such as *Tesseraspis* (**FigESM31**) and *Corvaspis*, exhibit a distinct dermal skeletal structure, obeying the same four-part structure seen in pteraspids, differing most significantly in the develop of L2 which is hugely expanded but, nevertheless, shows evidence of the same pattern of growth, with the initial establishment of the fretwork of walls from a fibre-rich matrix, followed by subsequent centripetal infilling of the remaining spaces.

**Figure ESM31:** The four-layered structure of the dermal skeleton in the tessellate heterostracan *Tesseraspis tessellata*, from the Lower Devonian of Earnestry Brook, Shropshire (NHM.PXXXX). The superficial D-layer (upper left, right) is comprised of sparse dental tubercles, separated by a thin structureless layer (L1) from the underlying spongy layer with centripetally-lined canals, some of which cross-cut the fabric of the matrix (lower left, right), reflecting resorption of the original matrix. The radial fabric centered on the matrix in the walls surrounding the vascular spaces in L2 (middle right) represent distortions in the collagen matrix occurring as a consequence of cell movement. The basal layer L3 has a rich fibre matrix exhibiting much evidence of distortion by cell movement (upper left, lower left).

#### Galeaspida

The dermal skeleton of galeaspids is composed of a fibre rich matrix organised into thick plywood-like layers that, though aligned parallel to the body wall, alternate in their orientation between successive layers (**FigESM32**). In our interpretative framework, this occurs through high expression of Collagen I in the D-layer. Delimiting this internally we observe a matrix discontinuity that we interpret as a remnant of L1. Beneath this discontinuity, the matrix exhibits a separate, plywood architecture characteristic of L3, intersected by a later developing vasculature that cross-cuts the original fabric of the matrix. The radial fabric of fibres within the matrix of the D-layer and L3 reflects the distortion of the Collagen I matrix as a consequence of cell migration (Point 3 above). A similar juxtaposition of D-layer and L3 is found in the Hand2 mutant mouse (**Fig.5 main paper, FigESM25 above**), implying that a Hand2 mediated process (of controlling Runx2 and elaborating L2) might have been secondarily lost in galeaspids, because heterostracans phylogenetically bracket all other skeletonised vertebrates.

**Figure ESM32:** Dermal skeleton of a polybranchiaspid indet. (IVPP V12603). A plywood-like structure occurs throughout (note that the lowest division represents the cartilaginous neurocranium-scapula ossification). Upper figure: there is a conspicuous discontinuity midway in the thickness of the dermal skeleton that represents the position of L1; L2 is not developed. The radial fibre orientation (vertical in this aspect; lower left, lower right) reflects distortions in the collagen matrix that occur as a consequence of cell movement.

#### Osteostraci

Prior to our discovery of the generative L1 in an extant taxon the osteostracan dermal skeleton has classically been interpreted as three-layered with a superficial layer of dermal tubercles (D-layer) overlying a middle vascular spongy bone layer and a basal lamellar layer. In the majority of species investigated, the bulk of the thickness of the dermoskeleton is comprised of a plywood-like arrangement of layers of cylindrically-shaped fibre-rich bundles that are parallel to the bodywall, but alternate in orientation within this plane, comparable to isopedin, characteristic of L3. The bases of the dermal tubercles (D-layer) are associated with a discontinuity that we can now interpret as L1. L2 intercalates L1 and L3, and expands in thickness through ontogeny, as reflected in a transect from the margin

(new growth) to the core (site of initial skeletal growth) of dermal plates (**Fig.ESM33** compare left and right side of the same plate). L3 undergoes resorption to open a network of vacuities that extends superficially and to the base of the dermal skeleton (**FigESM33, bottom right**) and, in some taxa, shows evidence of secondary mineralization by centripetal infilling (e.g. *Hemicyclaspis*, **FigESM33**). Thus, the regular plywood arrangement is broken down and replaced by a more chaotic arrangement of different generations of mineralization within L2, increasing the thickness of this division.

**Figure ESM33:** Dermal skeleton of the osteostracan *Hemicyclaspis*. Upper: a cranial dermal plate extending from margin (right) to core (left). The D-layer comprises a significant amount of the thickness of the plate (upper, lower right). Each of the dermal tubercles are separated from the underlying tissue by a thin structureless layer (L1). L2 occurs beneath this, comprised of a thin haphazardly structured layer at the margin, expanding in thickness towards the core (earliest formed part) of the dermal plate (lower left). L2 shows evidence of multiple episodes of resorption of a pre-existing matrix, opening vacuities, many of which are later infilled through deposition of new matrix. The expansion in thickness of L2 occurs at the expense of both the overlying D-layer and the underlying plywood-like L3 layer. Fibre bundles oriented oblique or orthogonal to the principal orientation of the plywood matrix reflect matrix distortions from cell movement.

*Tremataspis* is a representative of a derived clade of osteostracans, the tremataspids, and its histology is among the most thoroughly documented of all osteostracans (Denison 1947, 1951) (**Fig.ESM34**). The bulk of the dermal skeleton is again composed of a plywood-like arrangement of alternating sheets of like-aligned fibre-bundles within which there are vacuities that cross-cut the plywood structure; these vacuities are not secondarily remineralised. These vacuities connect to what have been interpreted as vascular spaces in an overlying cellular bone layer, which is overlain in turn by a continuous sheet of dentine and enameloid. There is some evidence of the ontogeny of the dermal skeleton in *Tremataspis*, but it remains unclear whether the evidence should be interpreted as a pattern of progressive mineralization (Denison 1947), perhaps of an otherwise unmineralised matrix, or of resorption (Denison 1951). The superficial enameloid/dentine layer represents the D layer and the cellular bone layer beneath it, L1; the plywood structure represents L3, but as we cannot identify a D-L1 discontinuity within the matrix, there is no discernable L2. The vacuities within L3 are similar to those found in L3 of late mouse dermal bone. As in the Hand2 mutants, the phenotype of *Tremataspis* shows independence and modularity of the underlying generative architecture.

**Figure ESM34:** Dermal skeleton of the osteostracan *Tremataspis*. The dermal skeleton is comprised mostly of D-layer and plywood-like L3, separated by a structural discontinuity (L1). There is no L2, although the vacuities within L3 (upper image and magnified successively in lower left and lower right) cross-cut the primary matrix and therefore represent resorption.

#### Placodermi

The structure of the dermal skeleton in placoderms is not well known. The antiarch *Bothriolepis canadensis* is by far the most thoroughly understood taxon (Downs and Donoghue 2009), known from articulated remains representatives of a spectrum of growth stages (Werdelin and Long 1986). The degree to which this taxon is representative of other placoderm species (Young 2010), but it exhibits a structure common to other placoderms that have been investigated, with the exception that it lacks the dermal tubercles that appear to have been a primitive features of placoderm families. Since the dermal tubercles are not our concern, *B. canadensis* remains a suitable model for understanding the structure and growth of the placoderm dermal skeleton. This is a helpful outcome since there is no alternate taxon represented by remains of comparable quality and quantity. Most importantly, we can observe all stages of L2 elaboration *within the same bony plates*, necessitating a malleable characteristic of biomineral that cannot be inferred easily from adult material. These early stages of L2 growth, hitherto unknown, also provide a facile comparison to the heterostracan morphology (and a similar L2 elaboration that was hitherto uninterpretable).

The study material consists of a growth series of thirteen fossil specimens of *Bothriolepis canadensis* (MHNM 02-52, 02-255, 02-265, 02-616, 02-640, 02-930, 02-1490, 02-1711, 02-2320, 02-2355, 02-2363, 02-2507, 02-2617) from the Late Devonian (Frasnian) deposits of the Escuminac Formation at the Miguasha field site in Quebec, Canada.

The dermal skeleton of *Bothriolepis canadensis* exhibits a four-layered structure, including a superficial D-layer comprised of continuous sheets of a bone-like tissue that is developed into a surface ornament resembling, but not representing, distinct dermal tubercles. The D layer exhibits evidence of external appositional growth based on the cross-cutting relationship between the incremental layering in the tissues and resorption horizons. There is a significant structural discontinuity, which we now identify as Layer 1, between the D-layer and the underlying cancellar Layer 2. The base of the dermal skeleton is comprised of a plywood-like L3 that includes cell lacunae flattened within the plane of the plywood structure.

Through studying the dermal skeleton in representatives of different ontogenetic stages we were able to resolve the development of this structure. Although we studied a range of dermal plates, the mixilateral (turquoise MxL in **Figure ESM35**) proved representative and our description of the developmental of dermoskeletal structure focuses on evidence from this dermal plate as it is well known to grow isometrically during *Bothriolepis* ontogeny (see scale bar below **FigESM35**).

**Figure ESM35:** The cranial dermal skeleton of *Bothriolepis canadensis* showing plate homologies and growth allometry (from Werdelin and Long 1986). The plate highlighted is the mixilateral (MxL) which was the focus of our analysis of the development of the dermal skeleton.

In the earliest growth stages that we studied (The external thoracic skeleton of the smallest individuals in the study sample (MHN 02-2355, 02-2617; AMD lengths of 10mm and 15mm, respectively), the dermal skeleton is comprised approximately half of D-layer and half of Layer 3; the structural discontinuity of L1 is evident and in MHN 02-2617 L2 is represented by a poorly structured zone within which small vacuities have begun to open (**Figure ESM36a**). These circular vacuities (see **Fig.4A**, arrow, main paper) are now interpretable as Col2/Col1 spherules surrounding invading cells. Layer 2 develops subsequently through inflation, where the existing mineral tissue in D-layer and L3 are separated progressively through the development of cancellae within L2 (**Figure ESM36b-d**). This must occur through a dynamic process of resorption and new growth of mineral, and the fabric of the tissue is dominated by spheritic mineralization (**Figure ESM36g,h**) that has been interpreted to represent rapid growth (Downs and Donoghue 2009) and can now be explained molecularly: As the invading Runx2+/OPN+ cells are initially blast-like and secrete Collagen2 (**FigESM10, above**), they could easily be mistaken for ‘chondrocytes’ while they are actually invasive osteoblasts (see **ESM chapter 1.1** above).

In MHN 02-2617, closer to the margins of the element, the vertical chambers of layer 2 are subdivided by vertical and horizontal sheets of bone connecting the walls to one another. In the sections taken, the smallest individual (MHN 02-2355)

shows only few of these horizontal sheets in the mixilateral element and these appear only in the thickened lateral corner of the element. In the same individual, vertical walls appear at the lateral and medial margins of the mixilateral, but the majority of the element is lacking any layer 2 development.

The lack of layer 2 is observed in only the smallest two specimens in the growth series and this is true of a much wider area in the very smallest individual (MHNM 02-2355). Sheets of bone connecting the vertical walls of layer 2 are infrequent in MHNM 02-2355 and are only observed in the deepened lateral corners of the element; these are more common in the slightly larger individual represented by MHNM 02-2617 and typify the entirety of layer 2 in all larger individuals in the sample. In the smallest individuals, mineralized spherites are rare but collect primarily at the base of layer 2. In larger individuals, spherites appear throughout layer 2 and are densely aggregated.

Vacuities develop subsequently within D-layer (close to the L1 interface) through resorption and secondary growth through centripetal appositional growth (**Figure ESM36c,d**). This results in a spongy or cancellar structure both above and below L1 (**Figure ESM36e**), sometimes obliterating the original L1 structural distinction (**Figure ESM36e,f**) and making comparisons harder between these late stages and heterostracan/mouse material.

This sequence of development, obtained from comparing homologous dermal bones across an ontogenetic spectrum of articulated individuals, is reflected also within individual dermal bones from late ontogenetic stage individuals. **Figure ESM36i** shows a transect through the structure of a mixilateral dermal plate, from the centre to the plate margin, which demonstrates the pattern of growth from the initial development of L2 (left; centre of dermal plate) through to the full developmental extent of L2 and the inflation of the D-layer (right, margin of dermal plate).

Juvenile placoderm specimens used in this study:

**Fig ESM36:**

- A: MHNM 02-2617, mixilateral
- B: MHNM 02-2355, mixilateral
- C: MHNM 02-2617, mixilateral
- D: MHNM 02-2617, mixilateral
- E: MHNM 02-616, mixilateral
- F: MHNM 02-616, nuchal cranial plate
- G: MHNM 02-616, anterior ventral lateral
- H: MHNM 02-616, mixilateral
- I: MHNM 02-2617, mixilateral

**Figure ESM 36:** Development of the structure of the cranial dermal skeleton in *Bothriolepis canadensis*. A. Initial growth stage in which the D-layer and L3 dominate; the structural discontinuity

of L1 is evident and L2 has begun to manifest as a structureless layer overlying L3. B. The dermal plate increases in thickness through inflation of L2 by resorption and development of struts that will eventually circumscribe cancellae or osteons. C, D. The cancellar walls extend in height and the D-layer also begins to increase in thickness through the development of secondary osteons. E, F. The mature dermal skeleton exhibits a cancellar or spongy structure within both L2 and D-layer; the structural distinction of L1 is often obliterated by resorption and new mineral growth. Nuchal cranial plate. G, H. Inflationary growth in L2 and D-layer is often characterised by spheritic mineralization; note the flattened cell lacunae in L3. I. This ontogenetic pattern is reflected also in the transect from plate core to plate margin.

In the more developed areas a second horizontal subdivision of cancellar spaces becomes obvious, which is remarkably similar to what we find in heterostracans (Fig.3M) and mouse (Fig.3K): Thus the temporal sequence of

- **1. layer2 expansion**, (Fig4B) with collagen1 trails traversing from L1 through L2 into L3 (Fig.4B arrows, feature 4)
- **2. Establishment of vertical cancellar walls** (Fig.ESM36c, Fig4B,C),
- **3. Establishment of further horizontal cancellar subdivisions** ( Fig4C left, 4E,4F, Fig ESM36d,e,g,i and Fig ESM37 below) is visible within the same bony structures of each taxon.

**FigESM37, taken from Jordan et al (accompanying paper), displays the temporal sequence of biomineral (yellow) remodelling within L2 by intercalary invasion of growing cellular sheets (grey lanes), secreting an anisotropic network of collagen scaffolds (Fig 2 this paper).**

This sequence of biomineral modification requires a substantial malleability that is entirely incompatible with an appositional process alone – now explained at the mechanistic level on the basis of single cell analysis and double pulse labelling of matrix deposition (FigESM37 taken from Fig.4g of the accompanying paper). The complex set of similarities across these three taxa (Fig.ESM18, features 1-6) points to a common underlying ontogenetic process of osteoblastic remodelling and intercalary matrix deposition that must have evolved among the ancestors of all skeletonised vertebrates.

#### 4. Acellular versus cellular L2 biomineral in ontogeny and evolution

Our in vivo double labelling experiment at later stages (**Fig.3** main paper) reveal details of mineral deposition and its spatial temporal sequence in L3 that could not be ‘inferred’ in any intuitive way. We see that labelled matrix is interspersed by later matrix sheets like in an accordion-like fashion / intercalary process. Thus, invasion and continuous deposition of coll I by osteoblasts appears to continue even into late more mature stages of dermal bone formation (**Fig.3G-J**). In those stages single cells albeit being polarised are fully surrounded by collagen matrix that either they or their predecessors have previously deposited. If such structures were to fossilise, one would recognise cellular lacunae inside plywood like matrix. In L3 regions that are most immature in the juvenile *Bothriolepis* we see very few lacunae within the plywood matrix (**Fig.4C**), while an adult *Bothriolepis* L3 carries many lacunae (**Fig.4S**). This suggests an invasion process in this cellular bone of placoderms, similar to what we find in postnatal stages of mice. Interestingly we see no lacunae within the plywood architecture of *Pteraspis* material (**Fig.4Q,R**), suggesting that invasion into L3 might be an ontogenetically and phylogenetically later process, linked to the emergence of cellular bone.

In the phylogenetic tree, we observe acellular bone first, and cellular bone later (**node 3, Fig.5**). The acellular bone matrix previously called aspidine, could hitherto not be understood by using a simple appositional mechanism. If the early stages of dermal bone formation in the mouse corresponds to early and late stages of dermal bone formation in basal gnathostomes, one would never expect to find cellular bone in L2: polarised cells would secrete coll I only in 1 direction, these cells would be bounded by the other side by endothelial and would not leave a deeper trace in the underlying collagen. Their respective migration paths would be reflected by the organisation of coll I fibres in L2, these coll I fibres that cover cancellar walls would sit on an amorphous matrix comprising coll II / coll I it has previously been shown that mixtures of coll I with other collagens or accessory factors can transform collagen fibre direction from an organised into an amorphous one. Osteostracans are the first basognathostomes with cellular bone, a feature that is retained in placoderms and all crown gnathostomes.

The striking similarity between early mouse, placoderm and heterostracan bones provides an evolutionary scenario where the emergence of L2 cellularity (i.e. cells being fully surrounded by matrix) happened by an ontogenetic ‘addition of an end stage’ (stippled line, top of **Fig.5**, node 3): A simple extension of invasive behaviour and the ability of single cells to leave sheets within L2 and L3 and migrate through (instead of only on top of) such collagen matrix would render a dermal bone cellular. This is because every cell would be bounded by collagen by all sides. Fortunately, the juvenile dermal bones of placoderms provide a remarkable match to the mouse material as regards to the elaboration of L3. In regions of the dermal bone, where L2 is not or poorly cancellated, we cannot discern cellular lacunae within L3. When we move in the same bone, in more mature regions L3 becomes increasingly populated by cells. In fact, in the adult placoderm, many flat lacunae have previously been described in the L3 plywood architecture, however, these were not interpretable as cells because the generative architecture that we outline here was unknown. High expression / concentration of coll I and invasive properties of osteoblasts into self-organising collagen I plywood meshworks would provide a parsimonious explanation for such morphological traces. We posit that this cellular feature / invasion into collagen matrices (would be a prerequisite for the emergence of what was hitherto cellular bone: bony matrix that would surround osteoblasts from all sites. We show such osteoblasts in mice at later stages, within L3 and L2, **figure 3G-J**).

#### Literature

- Abzhanov et al, 2007. Regulation of skeletogenic differentiation in cranial dermal bone, *Development*, 134: 3133-3144.
- Aeschlimann, D. & B. A. J. Evans (2004) The vital osteoclast: how is it regulated? *Cell Death and Differentiation*, 11, S5-S7.
- Agassiz, L. 1833-43. *Recherches sur les Poissons Fossiles*. Imprimerie de Petitpierre, Neuchâtel.
- Ahn, GO and Brown, JM. 2008 Matrix metalloproteinase-9 is required for tumor vasculogenesis but not for angiogenesis: Role of bone marrow-derived myelomonocytic cells *Cancer Cell*. March ; 13(3): 193–205
- Akech, J. et al. 2010. Runx2 association with progression of prostate cancer in patients: mechanisms mediating bone osteolysis and osteoblastic metastatic lesions, *Oncogene* 29, 811-821.
- Asagiri, M. & Takayanagi, H. 2007. The molecular understanding of osteoclast differentiation, *Bone* 40, 251-254.
- Boot-Handford, R.P. et al. 2003. A Novel and Highly Conserved Collagen (pro\_1(XXVII)) with a Unique Expression Pattern and Unusual Molecular Characteristics Establishes a New Clade within the Vertebrate Fibrillar Collagen Family, *JBC*, Vol. 269. No. 45, 28193-28199,
- Bystrow, A. P. 1955. The microstructure of the shields of the jawless vertebrates from Silurian and Devonian periods(eds). *Berg. Mem. vol. Acad. Nauk SSSR*. pp. 472-523.
- Couly et al. 1995. The angiogenic potentials of the cephalic mesoderm and the origin of brain and head blood vessels. *Mech Dev* 53, 97-112.
- Darnay, B. G., et al. (2007) TRAFs in RANK signaling. *Tnf Receptor Associated Factors* (Traf), 597, 152-159.
- Denison, R. H. 1947. The exoskeleton of *Tremataspis*. *American Journal of Science* 245: 337-365.
- Denison, R. H. 1951. The exoskeleton of early Osteostraci. *Fieldiana Geology* 11: 199-218.
- Denison, R. H. 1963. The early history of the vertebrate calcified skeleton. *Clinical Orthopaedics and Related Research* 31: 141-152.
- Denison, R. H. 1964. The Cyathaspididae: a family of Silurian and Devonian jawless vertebrates. *Fieldiana Geology* 13: 309-473.
- Denison, R. H. 1967. Ordovician vertebrates from Western United States. *Fieldiana Geology* 16: 131-192.
- Denison, R. H. 1973. Growth and repair of the shield in Pteraspididae (Agnatha). *Palaeontographica (Abt. A)* 143: 1-10.
- Donoghue, P. C. J. and Sansom, I. J. 2002. Origin and early evolution of vertebrate skeletonization. *Microscopy Research & Technique* 59: 352-372.
- Dougall et al. 1999 RANK is essential for osteoclast and lymph node development. *Genes & Development*, 13, 2412-2424
- Downs, J. P. and Donoghue, P. C. J. 2009. Skeletal histology of *Bothriolepis canadensis* (Placodermi, Antiarchi) and evolution of the skeleton at the origin of jawed vertebrates. *Journal of Morphology* 270: 1364-1380.

- Etchevers H. et al. The cephalic neural crest provides pericytes and smooth muscle cells to all blood vessels of the face and forebrain, *Development*, 128, 1059-1068.
- Fahlbusch, K., 1957. Pteraspis Dunensis Roemer, Eine Neubearbeitung der Pteraspidenfunde von Overath, *Paleontographica Abt.A*, 108, Liefg.1-4, 1-56 (Stuttgart)
- Ge et al. 2003. Nucleation of apatite crystals *in vitro* by self-assembled dentin matrix protein 1, *Nature Materials*, Vol2, 552-558.
- George A. , Veis, A. 2008. Phosphorylated Proteins and Control Over Apatite Nucleation, Crystal Growth and Inhibition, *Chem Rev.* 2008 November ; 108(11): 4670-4693.
- Giraud-Guille et al. 2003. Liquid crystalline assemblies of collagen in bone and in vitro systems. *Journal of Biomechanics* 36, 1571-1579
- Giraud-Guille et al 2005. Bone matrix like assemblies of collagen: From liquid crystals to gels and biomimetic materials. *Micron* 36, 602-608
- Giraud-Guille et al 2008. Liquid crystallinity in collagen systems in vitro and in vivo *Current Opinion in Colloid & Interface Science* 13 , 303-313
- Greeniaus, J. W. and Wilson, M. V. H. 2003. Fossil juvenile Cyathaspididae (Heterostraci) reveal rapid cyclomorial development of the dermal skeleton. *Journal of Vertebrate Paleontology* 23: 483-487.
- Gross, W. 1930. Die fische des mittleren Old Red Süd-Livlands. *Geologische und paläontologische Abhandlungen* 18.
- Gross, W. 1935. Histologische Studien am Aussenskelett fossiler Agnathen und Fische. *Palaeontographica (Abt. A)* 83: 1-60.
- Gross, W. 1961. Aufbau des Panzers obersilurischer Heterostraci und Osteostraci Norddeutschlands (Geschiebe) und Oesels. *Acta Zoologica (Stockholm)* 42: 73-150.
- Halstead, L. B. 1969. Calcified tissues in the earliest vertebrates. *Calcified Tissues Research* 3: 107-134.
- Halstead, L. B. 1973. The heterostracan fishes. *Biological Reviews* 48: 279-332.
- Halstead, L. B. 1974. *Vertebrate hard tissues*. Wykeham Science Publications Ltd., London.
- Halstead Tarlo, L. B. 1963. Aspidin: the precursor of bone. *Nature* 199: 46-48.
- Halstead Tarlo, L. B. 1964. The origin of bone. In H. J. J. Blackwood (eds). *Bone and Tooth*. Pergamon Press. pp. 3-17.
- Halstead Tarlo, L. B. 1965. Psammosteiformes (Agnatha) - a review with descriptions of new material from the Lower Devonian of Poland II - systematic part. *Palaeontologia Polonica* 15: 1-168.
- Huxley, T. H. 1858. On *Cephalaspis* and *Pteraspis*. *Quarterly Journal of the Geological Society, London* 14: 267-280.
- Hoang et al. 2003. Bone recognition mechanism of porcine osteocalcin from crystal structure, *Nature* 425, 977-980.
- Holler, K.L., et al., 2010. Targeted deletion of Hand2 in cardiac neural crest-derived cells influences cardiac gene expression and outflow tract development, *Dev. Biol. Developmental Biology*, Vol 341, issue1, 291-304.
- Hunter et al. 1994. Modulation of crystal formation by bone phosphoproteins: structural specificity of the osteopontin-mediated inhibition of hydroxyapatite formation *Biochem. J.* 300, 723-728
- Janvier, P. 1996. *Early Vertebrates*. Oxford University Press, Oxford.

- Jimenez et al. 1994. Collagenase 3 is a target of Cbfl, a transcription factor of the *runx* gene family involved in bone formation, *Molecular and Cellular Biology*, 19:6, 4431-4442.
- Kadler et al. 2008. Collagen fibrillogenesis: fibronectin, integrins, and minor collagens as organizers and nucleators *Current Opinion in Cell Biology*, 20:495-501
- Kapacee Z et al. 2008 Tension is required for fibroblast formation. *Matrix Biology* 27, 371-375
- Kikuchi et al. 2004. Biomimetic synthesis of bone-like nanocomposites using the self-organization mechanism of hydroxyapatite and collagen. *Composites Science and Technology* 64 , 819-825
- Kim et al. 2005. Osteoclast differentiation independent of the TRANCE-RANK - TRAF6 axis. *Journal of Experimental Medicine*, Vol. 202, No. 5., 589-595
- Kobayashi et al. 2001. Segregation of TRAF6-mediated signaling pathways clarifies its role in osteoclastogenesis. *EMBO Journal* Vol. 20 .6, 1271-1280
- Lindsley et al. 2004. Comparison of the four mouse fasciclin-containing genes expression patterns during valvuloseptal morphogenesis *Gene Expression Patterns* 5 (2005) 593-600
- Miyazaki et al. 2008. Corneal wound-healing in Osteopontin-deficient mouse, *Investigative Ophthalmology & Visual Science*, 8, Vol. 49, No. 4
- Moss, M. L. 1968a. Bone, dentin, and enamel and the evolution of vertebrates. In P. Person (eds). *Biology of the mouth*. American Association for the Advancement of Science, Washington DC. pp. 37-65.
- Moss, M. L. 1968b. The origin of vertebrate calcified tissues. In T. Ørvig (eds). *Current Problems of Lower Vertebrate Phylogeny*. Almquist & Wiksell, Stockholm. pp. 359-371.
- Nagoshi 2008. Ontogeny and Multipotency of Neural Crest-Derived Stem Cells in Mouse Bone Marrow, Dorsal Root Ganglia, and Whisker Pad, *Cell Stem Cell* 2, 392-403.
- Nishiyama et al. 1994. Type XII and XIV collagens mediate interactions between banded collagen fibers in vitro and may modulate extracellular matrix deformability , *JBC*, Vol 269, 11. ,pp 28193-28199
- Obruchev, D. V. 1941. Devonian fishes from the Minusinsk Basin [in Russian]. *Trudy paleontologicheskogo Instituta* 8: 23-48.
- Obruchev, D. V. 1964. Fundamentals of paleontology. Vol. XI Agnatha, Pisces, edited by Y. A. Orlov. Moscow: Nauka.
- Ørvig, T. 1951. Histologic studies of ostracoderms, placoderms and fossil elasmobranchs 1. The endoskeleton, with remarks on the hard tissues of lower vertebrates in general. *Arkiv för Zoologi* 2: 321-454.
- Ørvig, T. 1958a. *Pycnaspis splendens*, new genus, new species, a new ostracoderm from the Upper Ordovician of North America. *Proceedings of the United States National Museum* 108: 1-23.
- Ørvig, T. 1958b. Tanderna och tandvavnaderna genom tiderna (the teeth and their hard tissues through the ages). *Zoologisk Revy* 1958: 46-63.
- Ørvig, T. 1965. Palaeohistological notes 2: certain comments on the phylogenetic significance of acellular bone in early lower vertebrates. *Arkiv för Zoologi* 16: 551-556.

- Ørvig, T. 1967. Phylogeny of tooth tissues: evolution of some calcified tissues in early vertebrates. In A. E. W. Miles (eds). *Structural and Chemical Organisation of Teeth*. Academic Press, New York and London. pp. 45-110.
- Ørvig, T. 1968. The dermal skeleton: General considerations. In T. Ørvig (eds). *Current Problems of Lower Vertebrate Phylogeny*. Almquist & Wiksell, Stockholm. pp. 374-397.
- Pander, C. H. 1856. *Monographie der fossilen Fische des silurischen Systems der Russisch-Baltischen Gouvernements*. Akademie der Wissenschaften, St Petersburg.
- Pernegre V. 2006. Un nouveau ptéraspidiforme (Vertebrata, Heterostraci) du Dévonien inférieur du Spitsberg : nouvelles données paléo-ontogéniques, *Biodiversitas*, 28(2), 239-248.
- Petroll WM et al. 2004. Dynamic Three-Dimensional Visualization of Collagen Matrix Remodeling and Cytoskeletal Organization in Living Corneal Fibroblasts, *Scanning* VOL. 26, 1-10.
- Powrie, J. 1870. On the earliest known vestiges of vertebrate life; being a description of the fish remains of the Old Red Sandstone rocks of Forfarshire. *Edinburgh Geological Society Transactions* 1: 284-301.
- Rohon, J. V. 1893. Die Obersilurischen fische von Oesel. *Memoires l'Academie des Sciences de St.-Petersbourg, VII<sup>e</sup> Serie* 41: 124.
- Rubin et al. 2005.: Osteoclast: Origin and Differentiation , Bone Resorption , *Topics in Bone Biology* Volume 2, 1-23,
- Saito et al. 2002. A cell line with characteristics of the periodontal ligament fibroblasts is negatively regulated for mineralization and Runx2/Cbfa1/Osf2 activity, part of which can be overcome by bone morphogenetic protein-2 *Journal of Cell Science* 115, 4191-4200
- Sawhney R & Howard J. 2002 Slow local movements of collagen fibers by fibroblasts drive the rapid global self-organization of collagen gels *The Journal of Cell Biology*, Volume 157, Nr. 6, 1083-1090.
- Stensiö, E. A. 1927. The Downtonian and Devonian vertebrates of Spitsbergen. Part 1. Family Cephalaspidae. *Skrifter om Svalbard og Nordishavet* 12: 1-391.
- Toriseva, M.J. 2007. Collagenase-3 (MMP-13) Enhances Remodeling of Three-Dimensional Collagen and Promotes Survival of Human Skin Fibroblasts, *Journal of Investigative Dermatology*, Volume 127, 49-59.
- Vader D, Kabla A, Weitz D, Mahadevan L (2009) Strain-Induced Alignment in Collagen Gels. *PLoS ONE* 4(6): e5902. doi:10.1371/journal.pone.0005902
- Werdelin, L. and Long, J. A. 1986. Allometry in the placoderm *Bothriolepis canadensis* and its significance to antiarch evolution. *Lethaia* 19: 161-169.
- White, E. I. 1973. Form and growth in *Belgicaspis* (Heterostraci). *Palaeontographica Abt. A* 143: 11-24.
- Young, G. C. 2010. Placoderms (armored fish): dominant vertebrates of the Devonian Period. *Annual Review of Earth and Planetary Sciences* 38: 523-550.

#### 5. Detailed methods

See accompanying paper Jordan et al. 2013 (An intercalary mechanism governs radial growth of dermal bone).for immunohistochemistry, image acquisition and analysis.

To enable us to trace the cytoplasmic outlines of neural crest cells, we generated a novel recombinase reporter strain with a membrane bound vGFP (to be published in detail elsewhere). Biomineralization was explored using histological addition to calcein *ex vivo* in combination with multiplex immunohistochemistry (at 5mg/mL concentration), and *in vivo* via labelling experiments conducted under home office licence PPL 70/7178. To look at late aspects of biomineralization, intraperitoneal injections of each labelling agent were conducted on C57BL/6J mice from E11-E18 (with a minimum two day gap between injections) at the following concentrations: calcein 10mg/kg and xylene orange 90 mg/kg; specimen were isolated for analysis from P2 – P8. Rosette architectures along the surface of L1 were examined in dissected calvaria where the frontal and parietal bones were flat-mounted without removal of the overlying dermis, imaged through their depth by confocal microscopy and reconstructed in 3D segments.

##### Generation of transgenic animals

**XZ-DR**, bred and maintained by X. Zhang, University of Warwick. The XZ-DR transgenic is a Cre-reporter strain of mice generated by Dr Xintao Zhang. XZ-DR mice carry transgenic floxed resistance-pA cassettes in front of membrane bound-vGFP under the control of the Ubiquitin C promoter (to be described elsewhere). *Wnt1-Cre* mice were crossed with the XZ-DR-reporter yielding offspring in which the floxed cassettes are subject to Cre-mediated excision. The neural crest cells of these *Wnt1-Cre* x XZ-DR offspring are permanently labelled and can be directly visualised under fluorescence or by labelling with an anti-GFP antibody. Specimen that had been isolated, and embedded fresh (no fixation treatment).

##### Immunohistochemistry

Samples were prepared as above, with the following modifications: specimen from embryonic stage E16 onwards were fixed overnight in fresh 4% PFA at 4°C. All specimen for immunohistochemistry from embryonic stage E16 onwards were cut to a thickness of 12-17 µm onto charged slides, from E10-E14 specimen were cut to a thickness of 10 µm. Multiplex immunohistochemistry was performed on sections of embryos from all crosses listed above. Slides were defrosted at room temperature for 5 min, fixed in 4% paraformaldehyde for 15 min at room temperature, rinsed 3 x in PBS and blocked with permabilisation for 1 hour at room temperature. As a default, 20% Roche Western Block Solution (WBS) (Roche) + 0.1% Triton in PBS was used as the default blocking solution. Following blocking, primary antibodies were incubated at 4°C overnight in the blocking solution. The next morning samples were washed for 1-2 hours(s) with the wash solution (10% blocking solution without any detergent), with changes every 5-15 min. Secondary and conjugated antibodies were then incubated for 1 hr at room temperature in the Blocking Solution, followed by 1-2 hour(s) of washes (with frequent changes) and a PBS rinse. Slides with tyramide amplification (β-Galactosidase (AbCam) and Hand2 (R&D) antibodies) had the following additional steps: quenching of endogenous peroxidases with 0.3% H<sub>2</sub>O<sub>2</sub> during blocking; tyramide amplification (Perkin Emler Kit) with 5 µL conjugated (FITC, Cy3 or Cy5) tyramide in 500 µL Ampli-Buffer for 10-15 min; stopping of amplification reaction with 4% PFA or 0.01 N HCl for 10 min and 3 x PBS rinse. All slides were counterstained with DAPI (Invitrogen), administered either at 1:1000 for 5 min, or 1:2000 for 20 min with Rhodamine Phalloidin (Chemicon 1:250). Slides were then rinsed a final time in PBS and mounted in Mouwiol + DABCO. Slides were cover-slipped and sealed with nail varnish. Details of primary antibodies used can be found below. All secondary antibodies were obtained from Invitrogen and used at a concentration of 1:200.

##### PRIMARY ANTIBODIES

| Protein | Host | Working Dilution | Company | Reference |
| --- | --- | --- | --- | --- |
| α-SMA-FITC conjugated | Mouse, IgG2a | 1:200 | AbCam | ab8211 |
| α-SMA-cy3 conjugated | Mouse, IgG2a | 1:200 | Sigma | C6198 |
| β-Catenin | Mouse, IgG1k | 1:100 | Millipore | 05-665 |
| β-Catenin | Mouse, IgG1 | 1:100 | BD | 610153 |
| β-Galactosidase | Chicken, IgGY | 1:500+Amp1:1500 | AbCam | ab9361 |
| CD31 (PECAM) | Rabbit, IgG | 1:20 | AbCam | ab28364 |
| Collagen I | Mouse, IgG1 | 1:200 | AbCam | ab6308 |
| Collagen I | Rabbit, IgG | 1:100 | AbCam | ab21286 |
| Collagen I | Rabbit, IgG | 1:100 | AbCam | ab59435 |
| Collagen II | Goat, polyclonal | 1:100 | AbCam | ab24128<br>H0000737-M01 |
| Col14A1 (Undulin) | Mouse, IgG1k | 1:250 | Strattech |  |
| CRABP1 | Mouse, IgG2b | 1:100 | AbCam | ab2816 |
| Fibronectin | Rabbit, IgG | 1:100 | Sigma | F3648 |
| GFP | Chicken, IgGY | 1:400 | AbCam | ab13970 |
| Hand2 | Goat, polyclonal | 1:50+Amp1:1500 | AbCam |  |
| MMP9 | Mouse, IgG1 | 500 | R&D | AF3876 |
| MMP9 | Mouse, IgG1 | 1:200 | AbCam | ab58803 |
| MMP9 | Rabbit, IgG | 1:200 | AbCam | ab38898 |
| MMP13 (VIII A2) | Mouse, IgG1 | 1:100 | AbCam | ab52128 |
| MMP13 (Hinge-Region) | Rabbit, IgG | 1:100 | AbCam | ab39012 |
| Notch (activated) | Rabbit, IgG | 1:100 | AbCam | ab8925<br>NB600-1528 |
| Osteocalcin | Mouse, IgG1 | 1:100 | Strattech |  |
| Osteopontin | Rabbit, IgG | 1:100 | AbCam | ab63856 |
| Periostin (POSTN) | Rabbit, IgG | 1:100 | AbCam | ab14041 |
| Runx2 | Mouse, IgG2a | 1:200 | AbCam | ab76956 |
| von Willibrand Factor | Sheep, polyclonal | 1:500 | AbCam | ab11713 |

##### Ex vivo labelling

Matrix Labelling: histological addition of matrix mineral labelling agents in conjunction with immunohistochemistry was conducted prior to slide mounting following the immunological staining or in its absence. Mineral matrix of bone was stained with calcein made into solution with PBS, with 5 mg/50mL an effective concentration to allow visualisation of labelling. Slides were dipped solution for 10 sec – 45 sec, rinsed several times with PBS then mounted as described above.

***In vivo* labelling experiments (controls described in the detailed Materials of the accompanying paper XX)**

Animals obtained by continued use were used to map a time-course of bone mineralisation. Labelling agents were chosen that could be visualised without any treatment to enhance signal in emission areas with non-overlapping spectra such that they could be concomitantly analysed (for controls and details of image acquisition, see below). Concentrations of labelling agents were chosen such that the labelling was expected to be conferred to the dam and her offspring, without any pharmacological side effects, as determined in a host of previous studies on long bones (Tam and Anderson 1980, Simmon et al. 1981): calcein 10 mg/kg and xylene orange 90 mg/kg. Labelling agents were administered via intraperitoneal injection to mouse dams. Dosages were sufficient to confer the agents to the fetuses to achieve embryonic labelling. Mineralised bone was permanently labelled at 2 time-points between E11 and P0. Dosages were established from literature values on studies of bone-repair in long bones (Pautke et al. 2005, O'Brien 2002, Sun 1992, Vogt 2008) as above, and were prepared in Phosphate Buffered Saline (PBS, 137 mM NaCl, 2.7 mM KCl, 8 mM Na<sub>2</sub>HPO<sub>4</sub>, 1.4 mM KH<sub>2</sub>PO<sub>4</sub>, pH 7.4) and sterilised prior to administration. Specimen subjected to the protocol were terminated via an appropriate Schedule 1 method and the tissues were harvested (no sooner than 3 days after the last injection). Samples were fixed in fresh 4% PFA overnight, embedded in OCT (Optimal Cutting Temperature) media (TissueTek, VWR) and stored at -80°C until cryosectioning. Cryosectioning was performed on OTF 5000 Cryostat (Bright, UK) at a thickness of 15-17 µm. Samples were counterstained with 1:1000 DAPI (Invitrogen) and Image Acquisition and Analysis were conducted as per details below.

**Image Acquisition:** Confocal microscopy was performed using Leica TCS SP2 and SP5 systems using 10x, 20x, 40x, 63x, and 100x lenses. Alexa Fluor 488 was excited at 488 nm and emission was measured around 500 nm on the FITC channel; Alexa Fluors 546, 555, RPh and Cy dye 3 were excited at 543 nm and emission measured around 555 nm (TRITC channel). Alexa Fluor 647 and Cy dye 5 were excited at 633 nm and emission measured around 655 nm. DAPI stain was excited at 364 nm and emission measured around 400 nm. Reflectance images were acquired by exciting with the 488 laser and collecting using PMT. The measurement range was optimised to guarantee no overlap or bleed-through between the channels. All image acquisition occurred in the x, y and z-planes, resulting in a z-series. The average z-series comprised a step size of ~0.4-0.9 µm and the entire depth of the section was imaged. Brightfield images were acquired under a dissecting microscope, in a single optical plane.

**Image Acquisition: Matrix Labelling**

Samples that had been subjected to mineralised matrix labelling were imaged with the following parameters to ensure no spectral overlap occurred.

| Labelling | Excitation/Emission | SP5 Laser | SP2 Laser | Overlaps: |
| --- | --- | --- | --- | --- |

**Agent**

|  |  |  |  |  |
| --- | --- | --- | --- | --- |
| Calcein | 495 / 517 | Ar 100 mW<br>(488) | Ar 100 mW<br>(488) | FITC |
| Xylenol<br>orange | 440/570 / 610: 555 | DPSS 561 | HeNe 1.5 mW<br>(543) |  |

**Image Analysis**

**Image Analysis, 2D:** 2D image reconstructions were completed using LeicaLiteAS, ImageJ and Adobe Photoshop software. Images were either of individual optical sections from a z-series, or collapsed maximal or average projections of z-series. Image files from the confocal were converted into colour and channels were superimposed. Adjustments for brightness and contrast were made, in a linear fashion to the entire image, as needed.

**Image Analysis, 3D:** All 3D reconstructions of z-series were created using the freeware BioImageXD developed by the Universities of Jyväskylä and Turku in Finland and the Max Planck Institute of Germany: World Wide Web http://www.bioimagexd.net.
